# Multiple host species maintain genetic diversity in herbivore populations

**DOI:** 10.64898/2026.09.28.755086

**Authors:** Dilêr Haji, Erika R. LaPlante, Niels Agerbirk, Julien F. Ayroles, Joy Bergelson, Andrew D. Gloss, Noah K. Whiteman

## Abstract

How natural populations maintain genetic variation for fitness remains a central problem in evolutionary biology. Theory predicts that opposing selective pressures across environments can preserve fitness-associated polymorphism, but how interactions among species contribute to this process remains poorly understood. Here we show that host diversity preserves standing genetic variation within sexual populations of the leafmining fly *Scaptomyza flava*. We evolved twelve populations for ten generations on either of two naturally co-occurring host plants or on both, pooling survivors from the two hosts before mating. Single-host populations evolved reciprocal fitness trade-offs, whereas two-host populations maintained relatively even performance across hosts. Whole-genome time series dynamics identified a linked region in which alleles increased on one host and decreased on the other, while showing little net change under two-host conditions. Wild flies reared from the two hosts showed corresponding allele-frequency differences within this region. Across the genome, two-host populations retained more rare variants than populations evolving on either host alone. These findings show that selection imposed by different host species in sympatry can maintain standing genetic diversity within an herbivore population.

## Introduction

Evolutionary theory predicts that fitness-associated genetic variation arises by mutation and is largely removed by purifying natural selection and genetic drift as populations approach phenotypic optima (*1*, *2*). However, natural populations carry more genetic variation for fitness than these evolutionary forces alone predict (*3*, *4*). How fitness variation is maintained is a long-standing question in evolutionary biology (*4–6*), and critical for predicting the capacity of populations to adapt under rapid environmental change (*7*). One hypothesis for how fitness variation can be maintained is that balancing selection preserves polymorphisms when alternative alleles are favored in different environments *sensu lato* (*8–14*). Examples range from long-term negative frequency dependent selection (*15*) resulting in trans-species polymorphisms (*16*), to spatially varying selection across distinct habitats (*17*) and between fine-grained spatial scales within habitats (*18*) favoring large-effect alternative alleles at single genes. While these demonstrate that balancing selection can leave strong signatures at individual loci, considerably less is known about how balancing selection driven by environmental heterogeneity shapes the maintenance of standing genome-wide variation over space and time (*19*, *20*).

Evolve-and-resequence experiments in outcrossing, sexual species provide a powerful approach for testing how heterogeneous selection acts on standing genetic variation because they allow selective environments to be experimentally controlled while genome-wide allele-frequency change is followed directly over successive generations (*19*, *21*). Studies conducted in drosophilid flies have provided important new insights into the genetic basis of rapid adaptation from standing genetic variation for fitness (*22*, *23*). They have also revealed patterns of genome-wide diversity change driven by abiotic variation in the temporal (*24–26*) and spatial (*27*) environment, with populations maintained under heterogeneous regimes showing correspondingly higher inbreeding depression (*28*).

One potentially widespread form of balancing selection that could drive the maintenance of fitness-associated genetic variation within panmictic populations––particularly those involving biotic interactions––is a type of spatially-varying selection first formalized theoretically by Levene (1953). Consider a single population of an herbivorous insect species that attacks two different host plant species in sympatry. Each immature herbivore individual completes development on only one host plant individual and therefore experiences only that coarse-grained environment (*29*). Yet, that herbivore population, as a whole, encounters both host species. If the herbivores mate at random after they emerge from their host plants, this configuration will match the conditions required under models that result in protected polymorphism, provided alternative alleles are favored on the alternative hosts and each host species contributes a comparable share of the survivors (*10*). This general framework has been extensively elaborated upon (*30–34*) and studied in a handful of cases involving differing coarse-grained abiotic environments (*35–37*). However, whether and to what extent Levene-type heterogeneity generated by co-occurring biotic environments can maintain genetic variation for fitness, and the specific selective conditions that are sufficient to maintain genetic diversity under such a selective configuration at the genome scale, remains largely unknown.

We tested the prediction that Levene-type selective heterogeneity can maintain genetic diversity from standing genetic variation by combining a laboratory evolve-and-resequence experiment with a field genomic comparison in the leaf-mining drosophilid fly *Scaptomyza flava*. We evolved replicate populations of *S. flava* on alternative natural host-plant species in the laboratory and then compared the resulting patterns of allele-frequency change and genetic diversity with host-associated genetic variation in the field population from which the experimental populations were founded. *S. flava* is unusual among drosophilids because its immature stages develop entirely as endoparasitic, leafmining herbivores within living mustard leaves, including those of *Arabidopsis thaliana* in nature––they are culturable only on living plants (**Fig. 1A**). Although each immature fly typically experiences only the environment of a single host-plant individual, *S. flava* attacks more than 40 species of mustards and their relatives (Brassicales) (*38–42*). Here, we focused on *Barbarea vulgaris* and *Turritis glabra*, two locally abundant and sympatric host species of similar local abundance in sympatry (211 *B. vulgaris* and 215 *T. glabra* flowering stalks along a transect at the founder locality; Methods), which occur in fallow fields and other disturbed mesic habitats in New Hampshire, USA. The two hosts had distinct glucosinolate defense profiles (Supplementary Text), indicating that the two species represent ecologically distinct host environments for *S. flava*. Flies at our sampling sites attack both host species and are not known to form host races on them. Therefore, this system approximates the ecological selective conditions of the Levene model and enables a test of the extent to which antagonistic selection across alternative host environments can maintain genetic variation within a single herbivore population.

**Figure 1.**
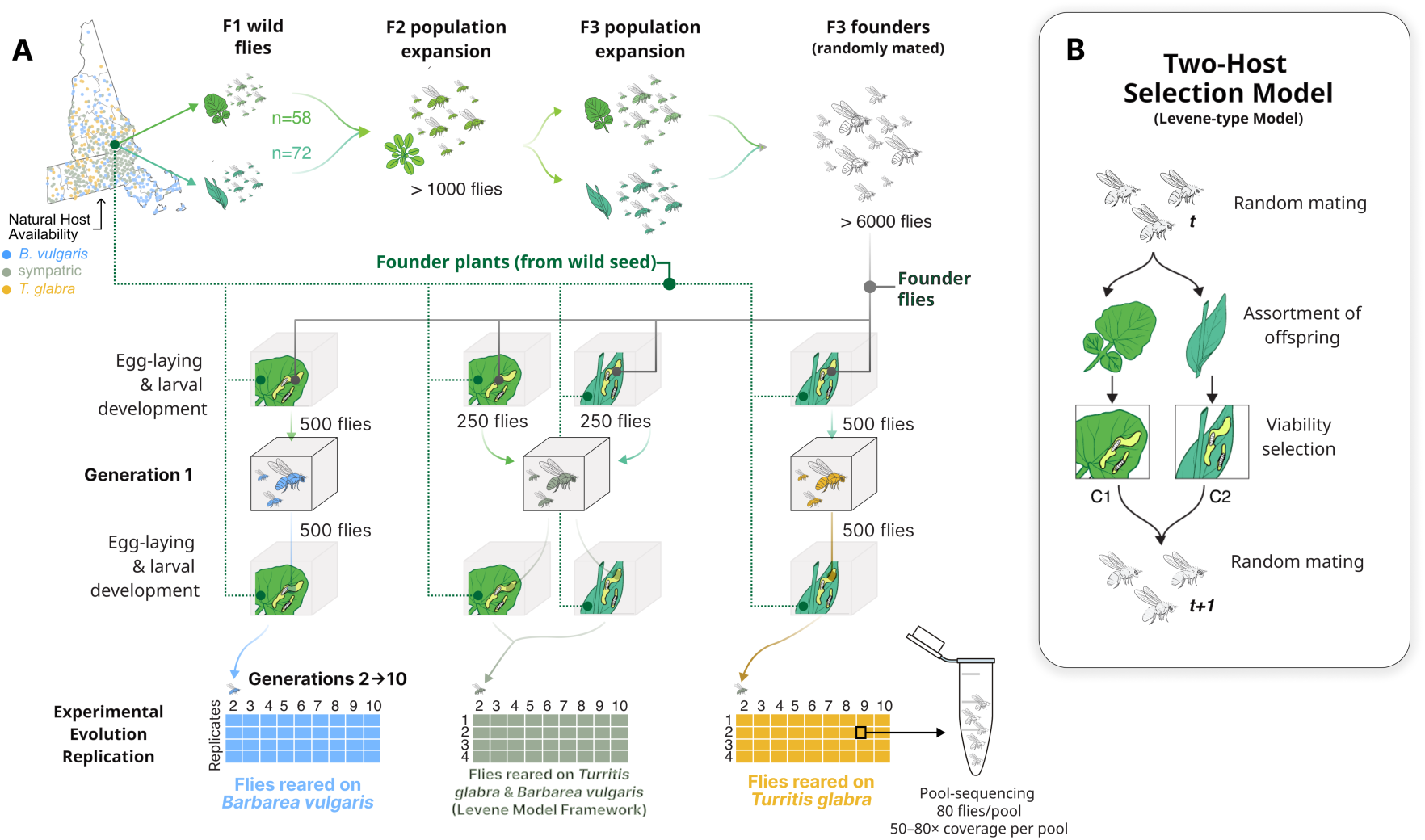
Experimental design for testing the maintenance of fitness-associated genetic variation through selection by multiple host species. **(A)** Wild *Scaptomyza flava* were collected from naturally infested *Barbarea vulgaris* (n = 58) and *Turritis glabra* (n = 72) at a site where both hosts co-occur in similar abundance (211 versus 215 flowering stalks along a transect); these 130 flies founded a laboratory colony expanded for two generations (*Arabidopsis thaliana*, then both hosts) before random mating. Wild-seed-derived stocks of both hosts were established from the same sites as the wild flies were collected in parallel. Four replicate populations evolved for 10 generations in three environments: *B. vulgaris* alone **(B)**, *T. glabra* alone (**T**), or both (**B+T**). In the two-host treatment, mated adults oviposited on a mixture of both hosts; infested plants were separated by host so that larvae developed within one host environment and 250 adults emerging from each host were pooled and randomly mated to start the next generation. Pool-sequencing example is shown. (**B**) Levene-type two-host selection model motivating the **B+T** treatment: after random mating, offspring are distributed among alternative hosts and experience host-specific viability selection before survivors rejoin a common mating population. The hosts differ in glucosinolate defenses (Figs. S2, S3; Tables S1–S4). Illustrations by Julie Johnson.

We generated a founder colony of *S. flava* in a laboratory growth chamber from 130 flies reared from naturally infected *B. vulgaris* (n = 58) and *T. glabra* (n = 72) leaves collected at a single locality. This colony was maintained for one generation on the permissive mustard host plant *Arabidopsis thaliana* (Col-0) and then for one generation on a combination of *B. vulgaris* and *T. glabra* (**Fig. 1A, S1**). In parallel, we established local, wild seed-derived stocks of both host plant species from the same locality. We then evolved flies for 10 generations under serial infection to model single-host and two-host species environments: single-host (uniform) selection driven by *B. vulgaris* (**B**) or *T. glabra* (**T**) and a two-host environment driven by both plant species available to flies under a Levene-type selection process (**B+T**). We evolved four replicate populations in each environment. Within each fly generation, we infected plants by allowing 500 mated adults to oviposit, collected surviving adults that emerged from leaves, and then used those adults to infect a new set of fresh plants. In the two-host environment, the population oviposited on a mixture of both host species; infested plants were then separated by host species into two sub-cages for larval development and 250 adults emerging from each host were retained and pooled into a single mating cage to initiate the next generation. Single-host populations were handled identically, with plants divided between two sub-cages, so that procedures did not differ among treatments. This design modelled coarse-grained selection within each generation (*10*, *29*). Each larva completed development inside leaves of a single host species and all survivors were allowed to then mate at random within each of their respective host plant treatments of origin (**B**, **T**, or **B+T**) (**Fig. 1B**).

## Results

### Experimental flies evolved reciprocal fitness trade-offs

If selection on each host favors different alleles in *S. flava* populations, we reasoned that ten generations of selection by a single host should improve performance on that host, whereas a population exposed to both hosts should improve less. We addressed this hypothesis first at the level of whole populations, counting the total number of flies surviving from egg to adult in the 2nd, 3rd and 10th generations (*t*_2_, *t*_3_ and *t*_10_). Viability increased in both single-host environments relative to the two-host environment (**Fig. 2A**). Relative viability **T** ÷ [**B+T**] rose to 128% of its starting value by *t*_10_ (Cohen’s *d* = 1.46, *p* = 0.031, one-sided), while **B** ÷ [**B+T**] rose by a similar amount but less consistently among replicates (Cohen’s *d* = 0.35, *p* = 0.27, one-sided). Taking both contrasts together, six of eight single-host cages increased, to 127% of starting values on average, relative to **B+T** (*p* = 0.087, one-sided). Realized heritability of the **T** contrast was high, at 60–69%, indicating that the response drew on additive genetic variance rather than on a change in rearing conditions (**Supplementary Text**). Single-host populations therefore evolved improved performance at faster rate than the two-host populations.

**Figure 2.**
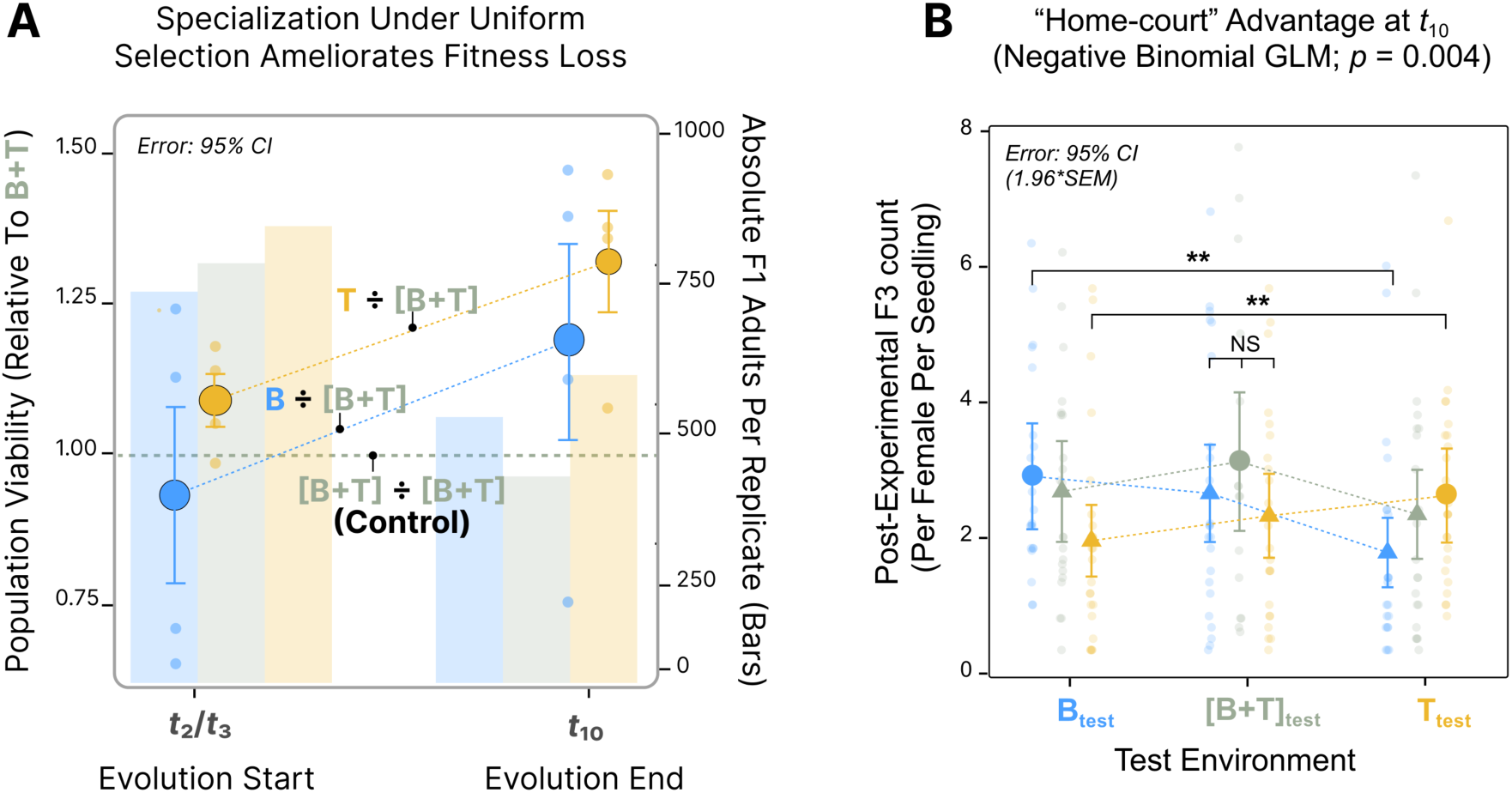
Experimental flies evolved reciprocal fitness trade-offs between hosts. (**A**) Single-host selection generated a relative viability gain compared with two-host selection, despite an overall fitness decline. Population viability, measured as total F1 adults per replicate (right axis, bars), is expressed as a ratio relative to the **B+T** reference (left axis, points) at the start of evolution (*t*_2_/*t*_3_) and at *t*_10_, for **B** ÷ [**B+T**] in blue, **T** ÷ [**B+T**] in yellow, and the [**B+T**] ÷ [**B+T**] internal control as the olive horizontal line. Large points are treatment means; error bars are 95% confidence intervals. (**B**) A reciprocal performance assay revealed host-antagonistic fitness trade-offs. Post-experimental F3 adult production per F2 female per seedling at *t*_10_ across three test environments (**B**_test_, [**B+T**]_test_, **T**_test_) for three evolution treatments: **B**-evolved in blue, [**B+T**]-evolved in green, and **T**-evolved in yellow. Small points are per-plant values, large points are treatment means with 95% confidence intervals, and connecting lines show reaction norms. Brackets show contrasts from the negative binomial generalized linear model (\*\**p* < 0.01; NS, not significant).

A viability gain under single-host selection is consistent with adaptation to that host. On its own, however, it does not show that alleles favored on one host are disfavored on the other. We therefore ran a reciprocal performance assay on the progeny of *t*_10_ flies to test for an evolved “home-court” advantage. To minimize maternal effects, post-evolution females were first allowed to oviposit on both host species. Groups of 30 F2 females then oviposited onto six *B. vulgaris* seedlings (**B**_test_), six *T. glabra* seedlings (**T**_test_), or a 3+3 mixture ([**B+T**]_test_). We refer to the environments of the evolution experiment as evolution environments and those of the assay as test environments. F3 adult production was 23% higher when the evolution environment matched the test environment (**Fig. 2B**; negative binomial GLM, *p* = 0.0257, **Table S5**). The advantage was strongest between the two single-host environments, where the reciprocal difference was 2.2-fold (*p* = 0.004). Flies evolved under two-host selection may nonetheless have performed better on average across all three test environments than single-host populations (*p* = 0.05), while matching them in their own home environment (*p* = 0.93). We found that this assay was compatible with measurements of population productivity in the experimental cages. The assay’s home-environment contrast (**Fig. 2B**; 1.09×, 95% bootstrap CI 0.79-1.63×) spans the 1.25× advantage of single-host over two-host population cages at *t*_10_ (**Fig. 2A**), an effect the per-plant assay (**Fig. 2B**) was underpowered to resolve (Fig. S4). Using the empirical selection intensity between **B**_test_ and **T**_test_ (*s* = 0.24), a balanced equilibrium was supported in ∼70% of bootstrap replicates under the Levene model, and both invasion conditions were satisfied: **B** genotypes could invade a **T**_test_ environment and **T** genotypes a **B**_test_ environment (both *p* < 0.001, **Fig. S5**).

These reaction norms describe a fitness set in the sense of Levins (*43*, *44*). Fitting the fitness set returned a convex hull (α = 7.61), the geometry under which a population exposed to both environments is not driven to either specialist optimum. Fitness nonetheless remained strongly contingent on the test environment (Fig. 2B), as did developmental timing (**Fig. S6**) (*40*, *45*). To express the assay in population-genetic terms, we refit the negative binomial model treating each evolution treatment as a genotype, with **B**-evolved as *AA*, **T**-evolved as *aa*, and [**B+T**]-evolved as *Aa* at *p* ≈ 0.5. This gave per-host selection coefficients of *s*_B_ = 0.39 and *s*_T_ = 0.33, and partial dominance of each locally favored allele in its own environment, with *h*_B_ = 0.78 and *h*_T_ = 0.67. Because these dominance coefficients sum to more than one, the heterozygote lies closer to the favored homozygote in each environment than an additive model predicts (*46*, *47*). Averaged across the two hosts at equal exposure, heterozygote fitness (0.865) exceeded that of both homozygotes (0.806 and 0.787). Substituting the per-host selection coefficients into Levene’s additive protection inequality gave a predicted equilibrium of *q*\*_phenotype_ = 0.55 for the **B**-favored allele under symmetric exposure, a quantity we return to below.

### Antagonistic selection maintained polymorphism in a single linkage block

The performance assay demonstrated antagonism at the level of the whole organism, suggesting an evolutionary response to selection. To identify regions of the genome underlying this selection, we next searched for genomic regions in which alleles rose in **B** and fell in **T**. No single test is reliable for this in a pooled time series, so we required agreement among several tests (*48–50*); that is, we applied a conservative test that required an antagonistically-evolving region must survive several different sources of noise. We integrated five tests in 200-kb non-overlapping windows across the four chromosome-scale scaffolds (110, 126, 439, 597; **Fig. 3A**). The tests were based on the most-negative tail of the per-window covariance of allele-frequency change between **B** and **T** (**Fig. S7**); the *z*-scaled temporal *F*_ST_ slope between **B** and **T** across *t*_1_–*t*_10_; permutation *F*_ST_ at *t*_10_ across all 70 four-versus-four relabellings of the eight **B** and **T** pools; the per-SNP generalized linear mixed model (GLMM) generation × treatment interaction likelihood-ratio test (LRT), aggregated to a per-window mean; and the BayPass *C*_2_ treatment-contrast statistic among the twelve *t*_10_ pools (**B**, **T** and **B+T**) (*51*, *52*). After density-stratified calibration of each test, one window reached the maximum vote of four of five: scaffold 439, 2.80–3.00 Mb (**Fig. 3A**, top). The window fell inside a ∼1-Mb founder linkage disequilibrium block (scaffold 439:2.64–3.61 Mb, **Figs. S8, S9**) (*53*), which is itself nested within a scaffold-439 parallel-selection hotspot on the X chromosome identified independently from the full time series (**Fig. S9, S10**) (*54*). The peak signal block contains orthologs of the *Drosophila melanogaster* Achaete-Scute Complex genes *ac*, *sc* and *l(1)sc*, which specify peripheral nervous system sensilla; the BMP antagonist *short gastrulation*; and the cuticle-pigment gene *yellow*, a well-known hotspot for *cis*-regulatory evolution (*55–60*). The gustatory receptor gene *Gr5a* and the cytochrome P450 genes *Cyp4g1* and *Cyp318a1* were identified outside of the signal block but inside the larger LD block (*61*, *62*). *Cyp4g1* is an insect-specific, oenocyte-expressed enzyme catalyzing the terminal oxidative decarbonylation step of cuticular hydrocarbon biosynthesis and is associated with desiccation tolerance and the penetration of contact insecticides (*63*, *64*). *Cyp318a1* exhibits site-specific positive selection in *Drosophila* (*65*). The signal block is dominated by non-coding variation. Of the top 5% of LRT SNPs, 76.5% are intergenic and only 3.7% are coding, against 15.7% expected from the 1.05-Mb flanking background, a 4.2-fold depletion of coding sites. Of the few coding variants, 81.7% are synonymous and none are nonsense.

**Figure 3.**
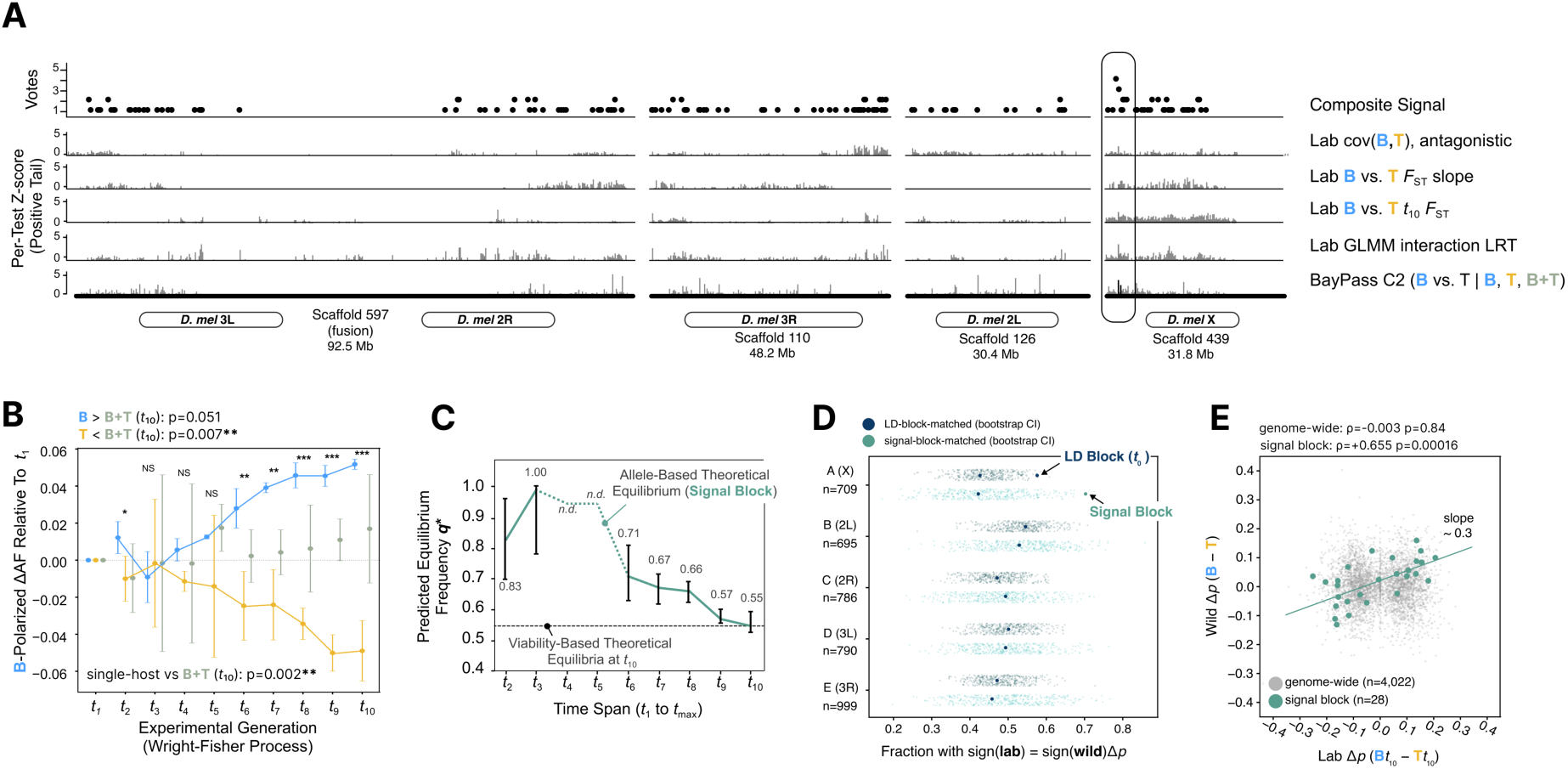
Antagonistic selection maintained polymorphism in a single linkage block on the X chromosome of *Scaptomyza flava*. **(A)** Multi-test vote scan across four chromosome-scale scaffolds spanning five Muller elements, labeled by *D. melanogaster* Muller-element homology. Top track, composite vote count per 200-kb window; lower tracks, positive-tail z-score per test. Signal block marked (439:2.80–3.00 Mb). **(B) B**-polarized ΔAF at the signal block relative to *t*_1_ for **B** (blue), **T** (yellow) and **B+T** (green); n = 4 cages each, **B**-favored allele assigned by leave-one-replicate-out cross-validation. Per-generation significance above; endpoint contrasts at left and bottom. **(C) *q\****_genomic_ across cumulative *t*_1_→*t*_max_ windows; median of 339 top-LRT SNPs in the LD block, ***q\**** = *s***_B_**/(*s***_B_** + |*s***_T_**|). Error bars, bootstrap intervals; n.d., median *s***_T_** non-negative. Dashed line, ***q\****_phenotype_ = 0.55 (Fig. 2B). **(D)** Per-Muller fraction of background SNPs with sign(lab Δp) = sign(wild Δp). Small points, founder-MAF-matched bootstraps matched to the LD block (navy, n = 67) or signal block (green, n = 28); filled circles, observed Muller A values; chance expectation, 0.50. **(E)** Wild Δp (**B** − **T**) against laboratory Δp (**B** − **T** at *t*_10_) for signal-block SNPs (green, n = 28; Spearman ρ = 0.655) and genome-wide background (grey, n = 4,022; ρ = −0.003). Green line, least-squares fit (slope ≈ 0.3).

Having identified a region in the *S. flava* genome that evolved in an antagonistic fashion between **B** and **T**, we next examined the trajectories of the alleles contained within that region. For each SNP in the signal block, we polarized on its **B**-favored allele under leave-one-replicate-out cross-validation, then tracked the polarized allele-frequency change relative to *t*_1_ across *t*_1_–*t*_10_ (**Fig. 3B**). The polarized allele rose monotonically in **B** to ∼0.05 by *t*_10_, fell symmetrically in **T** to ∼−0.05, and stayed near zero in **B+T** with wider per-replicate variance. Per-generation contrasts between the single-host treatments and **B+T** emerged at *t*_2_ and grew monotonically through *t*_10_. At *t*_10_, **B** > [**B+T]** (one-sided *p* = 0.051), **T** < [**B+T]** (one-sided *p* = 0.007), and the frequencies between both single-host environments diverged from **B+T** (*p* = 0.002). A density- and depth-matched genome-wide background frequency fluctuated around zero in all three treatments at every generation. In summary, each single-host population behaved like the homozygote favored on its own host, while the two-host populations behaved like the heterozygote and remained at intermediate frequency.

The performance assay and the allele-frequency data supported the hypothesis that antagonistic selection can maintain fitness-associated genetic diversity. We next asked to what extent the selection differentials between host species uncovered from both datasets supported the maintenance of genetic diversity under Levene’s theoretical model. We fit per-SNP logit slopes for **B**, **T** and **B+T** using cumulative *t*_1_→*t*_max_ windows for *t*_max_ from 2 to 10, took the median across the 339 top-LRT SNPs in the LD block, and substituted the median *s*_B_ and *s*_T_ into the additive equilibrium *q*\*_genomic_ = *s*_B_/(*s*_B_ + |*s*_T_|). At small *t*_max_ the estimate was dominated by the large early changes from the founder bottleneck, giving *q*\*_genomic_ = 0.83 at *t*_max_ = 2, and the **T** slope was not yet negative, leaving the equilibrium undefined at *t*_max_ = 4 and 5. As generations accumulated, the **T** slope became reliably negative, the **B** slope stabilized, and the estimate converged: 0.71 at *t*_max_ = 6, then 0.67, 0.66, 0.57, and 0.55 at *t*_10_ (**Fig. 3C**). The *t*_10_ value matches *q*\*_phenotype_ = 0.55 derived from the reciprocal performance assay (**Fig. 2B**). An independent set of selection coefficients estimated from within the same LD block using a temporal least-squares estimator (*66*) gave 0.66, within 0.11 of the phenotypic prediction. Antagonistic selection at a single locus is therefore sufficient to account for the performance trade-off, and the equilibrium geometry is mirrored between genotype and phenotype.

We next asked to what extent these experimentally-evolved selection differentials could predict selection differentials in wild populations exposed to the same host species. For each background SNP we recorded whether sign(lab Δ*p*) = sign(wild Δ*p*). Lab Δ*p* was the difference between the **B** and **T** allele-frequency change at *t*_10_, averaged across the four laboratory replicates, and wild Δ*p* was the difference in allele frequency between flies reared from **B** and from **T** hosts, averaged across wild *S. flava* pools collected after one generation of natural selection during immature development in **B** or **T** hosts at the field sites (**Fig. S11, Table S6**). The chance expectation is 0.50. Genome-wide, concordance matched chance on all Muller element homologs of *D. melanogaster* (**Fig. 3D**): 0.46 ± 0.02 on A (*n* = 709), 0.53 ± 0.02 on B (*n* = 695), 0.48 ± 0.02 on C (*n* = 786), 0.49 ± 0.02 on D (*n* = 790), and 0.48 ± 0.02 on E (*n* = 999). There was no genome-wide correlation between laboratory antagonism and wild between-host differentiation (Spearman ρ = −0.003, *p* = 0.84, *n* = 4,022). The scaffold 439 LD block (Muller A), however, showed a concordance of 0.58 ± 0.06 (*n* = 67 SNPs), and within it the signal block showed a concordance of 0.71 ± 0.09 (*n* = 28 SNPs). Both values exceeded the 95% upper bound of 1,000 size-matched and founder-MAF-matched bootstrap samples drawn from the same Muller element homolog, including A, whose background median was 0.43. Laboratory and wild allele-frequency differences were strongly correlated at the signal block (ρ = 0.655, *p* = 1.6 × 10^−4^), with about one third of the laboratory between-treatment change recovered as wild between-host differentiation. A single generation of selection in the wild was therefore sufficient to shift alleles within the signal block in the same direction as ten generations of laboratory evolution, but the magnitude of the shift was weaker after one generation of wild selection compared to 10 generations of lab selection (**Fig. 3E**).

### A two-host environment preserved more rare variants than single-host environments

Having localized antagonistic selection to one block and confirmed its direction in the wild, we turned to its genome-wide consequences. The model predicts that a two-host environment removes variation less efficiently than a single-host environment, because variation that reduces performance on one host is returned to the population each generation by the alternative host. To test this prediction, we tracked allele-frequency trajectories of all twelve experimental cages through ten generations using measurements of θ_π_, Watterson’s θ_W_ and Tajima’s *D* for the wild samples, the laboratory founder colony and the twelve *t*_10_ cages. The founder colony was represented by four independent 80-fly subsamples, and 128 individually sequenced founders showed no residual host-of-origin structure (**Fig. S12**). All analyses were restricted to the same 812,670 biallelic SNPs ascertained jointly across the **B**, **T** and **B+T** time series. Projecting the per-cage *t*_1_–*t*_10_ trajectories onto the principal components of allele-frequency space showed that **B** and **T** cages evolved along a shared uniform-selection axis, while the four **B+T** cages diverged onto a distinct axis (**Fig. 4A**). The *t*_10_ centroid of the **B+T** cages lay *d* = 66.2 units from the midpoint of the **B** and **T** *t*_10_ centroids in PC1–PC10 space, at an angle of 121° from where **B+T** would fall if it were simply intermediate between **B** and **T** (permutation *p* = 0.055). Evolution under two hosts therefore followed a trajectory not shared with either single-host regime.

**Figure 4.**
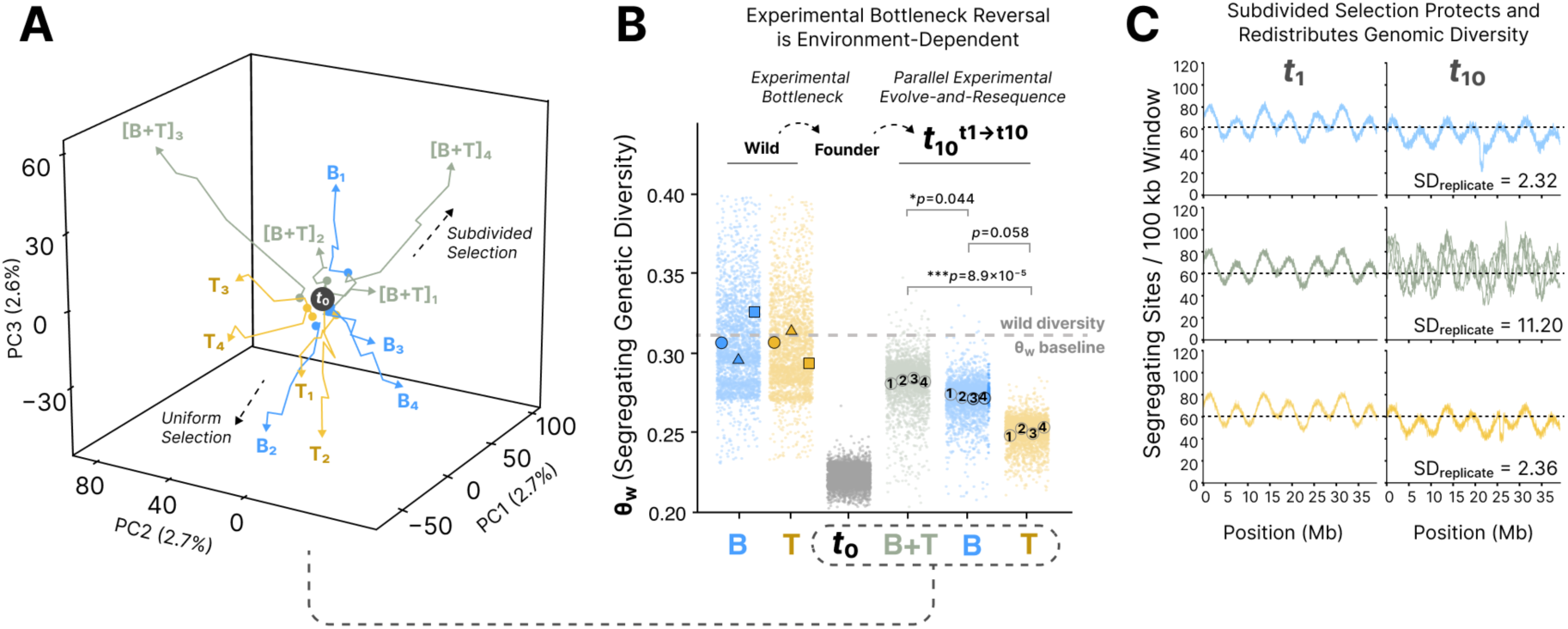
A two-host (Levene-type) environment preserved genomic diversity compared to single-host selection. (**A**) **B+T** diverged from **B** and **T** along a non-intermediate axis of genomic variation. Three-dimensional projection of per-cage allele-frequency trajectories (*t*_1_ to *t*_10_) onto the first three axes of the joint allele-frequency PCA (812,670 SNPs); arrowheads mark *t*_10_ endpoints. (**B**) **B+T** recovered rare-variant genetic diversity more strongly than **B** or **T** after the experimental bottleneck. Environment-dependent bottleneck reversal in Watterson’s θ_W_ across wild, founder and *t*_10_ samples, weighted by the *t*_1_ to *t*_10_ time-series slope in non-overlapping windows. Large points are per-pool means with replicate number shown; the dashed line is the wild diversity baseline. Significance is from a linear mixed-effects model with treatment × generation and per-replicate random intercepts and slopes. (**C**) **B+T** retained rare-variant sites idiosyncratically among evolution replicates. Per-100-kb segregating-site density along chromosome-scale scaffold chr_ScDA7r2_110 (48.2 Mb) at *t*_1_ (left column) and *t*_10_ (right column) for **B** (top, blue), **B+T** (middle, olive) and **T** (bottom, yellow); each panel overlays the four replicate cages of that treatment, and SD_replicate_ is the mean across windows of the between-replicate standard deviation in segregating-site count.

We focused on θ_W_, Watterson’s estimator, as the primary diversity statistic. θ_W_ is proportional to the number of segregating sites and is therefore dominated by rare alleles, which are the alleles most easily lost in a founding bottleneck and most readily returned to detectable frequency by experimental selection. θ_π_, by contrast, is weighted toward intermediate-frequency alleles, which should survive a moderate bottleneck. Establishing the founder colony produced the expected bottleneck signature (**Fig. 4B)**. θ_W_ fell by 26%, from 0.300 to 0.224 (Wilcoxon paired *p* < 10^−30^, **Fig. S13B**), and Tajima’s *D* flipped in sign, from −0.86 ± 0.12 in wild populations to 0.83 ± 0.08 in the founders (*p* < 10^−30^, **Fig. S13C**). θ_π_ showed no comparable signature and instead rose 11% above the wild median (*p* < 10^−30^, **Fig. S13A**); the expected loss of heterozygosity under drift alone, at an effective size of a few hundred diploids over ten generations, was under 3% (**Fig. S14B**). Any later recovery of θ_W_ in an evolving cage, absent a change in population size, must therefore reflect either new mutation or the return of previously low-frequency alleles to higher frequency by positive selection. After ten generations, the single-host **B** and **T** environments retained the founder signature (**B**: θ_W_ = 0.246 ± 0.031, *D* = 0.32 ± 0.55; **T**: θ_W_ = 0.228 ± 0.009, *D* = 0.63 ± 0.13). The two-host environment partially reversed it, raising θ_W_ from 0.224 to 0.271 ± 0.029 and returning Tajima’s *D* to approximately zero (−0.11 ± 0.52). This left **B+T** closer to the wild mean of ∼0.31 than to the founder mean of ∼0.22. New mutation cannot account for the recovery because at the *D. melanogaster* mutation rate of μ = 2.8 × 10^−9^ per site per generation (*67*), ten generations would contribute ∼5 × 10^−5^ to θ_W_, which is three orders of magnitude below the observed Δθ_W_ of ∼0.047. The rare variants that re-emerged in **B+T** were therefore pre-existing founder alleles driven upward by positive selection. In the linear mixed-effects model of θ_W_ trajectories (Methods), **B+T** exceeded both **T** (estimated Δ = 0.029, *p* = 8.9 × 10⁻⁵) and **B** (Δ = 0.015, *p* = 0.044), while **B** and **T** differed only weakly (Δ = 0.014, *p* = 0.058). The reversal was consistent across cages: per-replicate θ_W_ slopes over *t*_1_–*t*_9_ were positive in all four **B+T** cages, at +0.002 to +0.004 per generation (**Fig. S15**).

Between-replicate divergence, measured as the normalized variance of allele frequency among replicates within a treatment, increased linearly with time in all three regimes but at different rates. After correcting for technical noise, **B+T** diverged at the lowest apparent effective size (*N*_e_ ≈ 219), followed by **B** (≈ 242) and then **T** (≈ 347), against a census size of 500 (**Fig. S14C, Table S7**); two independent estimators gave effective sizes of similar magnitude (N_e_ ≈ 200–350) but not the same ranking (**B+T** < **B** < **T**, **Figs. S16, S17**). We interpret the excess variance in **B+T** as a consequence of opposing selection pressures given that the two host species add a subdivided-selection variance component that is absent from single-host regimes, leading to inflation of variance-based estimates of effective size. To separate drift from positive selection, we quantified genome-wide linked-selection intensity using a temporal-covariance framework (*68*), with per-treatment covariance matrices computed after correcting for pool-size overdispersion (**Figs. S18–S20**). The fraction of allele-frequency change attributable to selection, *G*, computed from off-diagonal temporal covariances at *k* ≥ 2, was 0.06 in **B+T** with all four evolutionary replicates positive (one-sided permutation *p* = 0.045, **Fig. S21**). The within-interval convergence correlation recovered similar patterns (**Fig. S22**), and simulations confirmed that the design had limited power to resolve a convergence correlation of this size (**Fig. S23**). **T** showed the same direction (*G* = 0.04, three of four replicates positive) but did not reach significance because of one negative outlier, and **B** showed no net signal (*G* = −0.03, replicates mixed in sign). The two-host regime therefore combined elevated stochastic variance with parallel genome-wide signatures of linked selection across replicates. It also differed in which sites were retained. The average between-replicate standard deviation of per-window segregating-site density at *t*_10_ was roughly five-fold higher in **B+T** than in either single-host treatment (11.20 in **B+T** versus 2.32 in **B** and 2.36 in **T**; Mann–Whitney *U*, both *p* < 10^−10^; **Fig. 4C**). Where each single-host regime converged on a shared subset of surviving rare variants, the four **B+T** replicates retained idiosyncratic subsets scattered across the genome.

These genome-wide patterns followed from the locus-level result. Linkage disequilibrium in this population extended over roughly a megabase. Given that the experiment was run for ten generations, we expected widespread hitchhiking of linked genetic variation. In a single-host environment the response to selection efficiently removed both genome-wide genetic variation as selective sweeps led to the loss of linked genetic variation. In the two-host environment, opposing selection each generation resulted in incomplete selective sweeps of varying direction that enabled the retention of more genetic variation.

## Discussion

Although environmental heterogeneity has been proposed for more than seventy years as a mechanism by which balancing selection can maintain genetic diversity in nature, few experimental tests of the theory proposed as a mechanism have been conducted. In Levene’s classic model, individuals experience selection in discrete ecological niches before survivors rejoin a common mating population, allowing alternative alleles to persist when fitness changes sufficiently among niches (protected genetic polymorphism) (*10*). Subsequent theory showed that this depends on the life cycle of the species under selection, the relative contribution of different habitats to reproduction, migration, dominance, linkage, and the spatial scale of density regulation (*8*, *30*, *32–34*, *69*). Environmental heterogeneity alone therefore does not imply that balancing selection will operate. A central empirical challenge has been to demonstrate that alternative environments impose opposing selective pressures and that the resulting allele-frequency dynamics can protect genetic polymorphism for fitness. This distinction is particularly important at the genomic level, where demography, linked selection, population structure, and other selective processes can produce elevated diversity or intermediate-frequency alleles without generating protected polymorphisms *per se* (*8*, *69*, *70*).

The prediction that heterogenous environments preserve more genetic variation than homogenous environments has substantial empirical precedent. In one of the earliest direct tests, Powell (1971) (*37*) evolved *D. willistoni* populations under homogenous versus heterogeneous combinations of food and temperature, finding greater allozyme variation in the heterogeneous environments. Related experiments subsequently showed similar effects of environmental and competitor heterogeneity (*71*, *72*). A later experiment in *Escherichia coli* manipulated selection regimes and found that spatial heterogeneity maintained polymorphism under soft but not hard selection (*73*). However, the lack of genomic resources impeded the discernment of whether the implicated loci were experiencing selection. More recently, evolve-and-resequence experiments using genome-wide datasets have shown that adaptation from standing variation is often highly polygenic, involving different allele subsets in replicate populations exposed to the same selection regime (*22*, *23*, *74*, *75*). Experimental and field studies have further found that allele frequencies can track temporally heterogeneous environments over remarkably short ecological timescales, including seasonal oscillations (*24–26*, *76*). Genome-wide differentiation also distinguishes replicated cadmium- and salt-enriched selective regimes (*27*) in *D. melanogaster*. Experimental evolution in *Brassica rapa* plants has demonstrated that herbivory modifies evolutionary responses to bumble bee pollination, highlighting how interacting selective agents shape adaptation (*77*, *78*). Herbivores that have been evolved experimentally include mites maintained on alternating host plants, resulting in the evolution of broader niche breadth compared to single-host lines (*79*, *80*). Those designs alternated hosts between generations, whereas ours subdivided each generation between hosts and returned all survivors to one mating pool––the spatial soft-selection configuration for which protected polymorphism was originally derived (*10*). Combining the Levene-type configuration with a replicated genome-wide time series here, enabled us to quantify how alternative natural host species shape fitness trade-offs and the retention of standing genetic variation.

Plant-herbivore interactions have long been considered a particularly promising context for testing evolutionary theory. Individual herbivorous insects often complete development on a single plant while populations encounter genetically and chemically heterogeneous host plants. These attributes of the life cycle generate a coarse-grained selective landscape–whereby individuals experience a single selective regime, but populations experience many––envisioned by early models of spatially-varying selection (*10*, *81*, *82*). Host plants can therefore act as selective agents that drive both adaptation between herbivore species and within herbivore species. When host-associated mating or restricted migration allows divergence to accumulate, divergent selection can generate local adaptation, host races, and ultimately reproductive isolation, as documented in pea aphids, *Rhagoletis*, and *Timema* (*83–88*). With sufficient mixing of herbivores among hosts, however, the same environmental heterogeneity could instead maintain variation within a population, as we show. Although the genomics of host-associated divergence has been relatively well characterized, within approximately panmictic populations, the amount of genetic variation maintained and the specific selective dynamics required for its maintenance by host heterogeneity has remained largely unknown (*82*, *89*).

This study addressed the problem of how host heterogeneity can maintain genetic diversity within populations when an individual experiences a single environment (coarse-grained selection) but the population experiences multiple environments. In this case, *S. flava* immatures (eggs, larvae and puparia) each experienced host plant-driven (bottom-up) viability selection within alternative host species, but the survivors were returned as adults each generation to a common mating population to create the next generation. We found that fitness trade-offs evolved within ten generations of experimental selection on each alternative host species. Fitness trade-offs have been central to theories of ecological specialization (*90*, *91*) because antagonistic pleiotropy provides a relatively simple explanation for why adaptation to one resource should constrain performance on another. Yet, empirical evidence from herbivorous insects have generated mixed results. Negative genetic correlations in performance among hosts are often weak or undetectable, but positive correlations caused by variation in vigor are common (*92*, *93*). Studies that have leveraged reciprocal transplant experiments have frequently found evidence for local adaptation, but the corresponding fitness tradeoffs between environments are substantially weaker (*94*), likely owing to complex genetic architectures shaped by the interplay of behavior, ecological fitting, and multitrophic interactions (*95*). By conducting a laboratory evolve-and-resequence experiment, coupled with tests for local adaptation, we were able to detect a surprising 2.2-fold fitness tradeoff driven by selection from the single host plant species alone, while simultaneously finding that populations evolved on more complex host environments (two host species) maintained relatively even performance across both one-host and two-host treatments.

Comparable effects of host heterogeneity have emerged across remarkably different antagonistic interactions: *Brachiola algerae* evolved on *Aedes aegypti* (*96*), *Serratia marcescens* evolved on *Caenorhabditis elegans* (*97*), bacteriophage evolved on *E. coli* (*98*), arboviruses evolved on alternating vertebrate and mosquito hosts (*99–101*), and *Aphidius ervi* parasitoid wasps on aphid hosts (*102*). Together with the broader observation that performance trade-offs arise more frequently during adaptation to homogeneous than heterogeneous environments (*103*), these studies establish that host diversity can alter the evolution of specialization. However, a major gap is in understanding the relationship between the biotic pressures that select for specialists and generalists and the consequences of those selective dynamics on standing genetic variation within sexual populations.

At the genomic level in our study, single-host and two-host selection produced different outcomes for the *S. flava* populations: selection on either host alone increased relative viability while eroding rare-variant diversity, whereas exposure to both hosts retained more of this variation. One potential explanation is that host heterogeneity weakened selection against variants whose deleterious effects were expressed on only one host. Theory predicts that conditional expression can increase standing variation by reducing the efficacy of purifying selection (*104*). Antagonistic fitness effects across hosts could also contribute to the retention of particular variants. These mechanisms have different implications: an allele may persist because selection against it is weaker or because opposing selective pressures actively protect it from loss. Establishing which mechanism predominates––and whether retained variants explain the lower relative viability of two-host populations––will demand linking individual variants to their fitness effects on each host, which will require the sequencing of individuals.

In conclusion, our results connect the evolution of host specialization with the fate of standing genetic variation within a sexual herbivorous insect population. Fitness trade-offs emerged rapidly under single-host selection, while populations exposed to both hosts retained more genetic variation and maintained relatively even performance across host environments. The diversity of biotic interaction among members of natural ecological communities has the potential to reshape genetic diversity for fitness within populations of their constituent species. Understanding how variation persists within herbivorous arthropods and other parasitic animals—which are estimated to comprise nearly half of all multicellular species of life—will require experimental approaches allowing for the prediction and measurement of the diversity of selective forces imposed by their hosts.

## Methods

### Study system and wild collections

*Scaptomyza flava* (Diptera: Drosophilidae) is an oligophagous leaf-mining fly whose immature stages (egg to larva to puparium) develop entirely within the living leaves of mustards (Brassicaceae and other Brassicales). In the fallow fields we surveyed in New England, *S. flava* attacked two sympatric hosts, *Barbarea vulgaris* and *Turritis glabra*. In the summer of 2014 we collected third-instar larvae mining leaves of naturally established *B. vulgaris* and *T. glabra* at three sites (AV, PS, RM) near Newington, New Hampshire, USA, bagged the parental leaves, and reared larvae to adult emergence. We retained 130 adults (58 from *B. vulgaris*, 72 from *T. glabra*) from the site yielding the most emergences to found the laboratory colony. At each of the three sites, emerged adults were pooled separately by host species (22-60 flies per pool) for pooled whole-genome sequencing, giving six wild libraries (AVB, AVT, PSB, PST, RMB, RMT). Seeds of both host plant species were also collected from the same wild populations and propagated in an indoor growth chamber to generate the bank of experimental plants used throughout. Host-plant occurrence records shown in Fig. 1A were downloaded from iNaturalist (https://www.inaturalist.org; accessed February 2026) for New Hampshire and Massachusetts (*B. vulgaris*, taxon 56126, n = 3,711 records; *T. glabra*, taxon 79460, n = 1,695). Records were clipped to state boundaries and binned on a 40 x 40 grid with each occupied cell classified as *B. vulgaris* only, *T. glabra* only, or sympatric when both species were recorded in the same cell. Local host abundance was measured in May 2015 by counting flowering stalks of both species along transects at two sites: 211 *B. vulgaris* and 215 *T. glabra* at the Pease (PS) founder site, and 99 *B. vulgaris* and 182 *T. glabra* at Rock Meadow, Massachusetts. Non-flowering rosettes were not counted because they could not be identified reliably among surrounding vegetation.

### Founder colony and experimental evolution

The 130 wild-derived adults were interbred and maintained for two generations, the first on *Arabidopsis thaliana* (Col-0) and the second on a mixture of *B. vulgaris* and *T. glabra*, and the outbred colony was expanded to ∼15,000 individuals before subdivision. In Fall 2014 the colony was split across nine mesh cages provisioned with experimental plants on a two-day oviposition schedule designed to remove bias from host preference: four cages received *B. vulgaris* on day 1 and *T. glabra* on day 2, four received the reverse order, and one received a mixture. Infested plants were then redistributed across ten rearing cages so that emerging adults were separated by host of origin while pooling offspring from pairs of the original nine cages (**Fig. S1**). Each of the ten founder cages was split into batches of 25 flies, and each experimental cage was founded with two batches from each founder cage, producing genetically balanced populations of 500 flies. This allocation was performed on two dates, founding six populations (B1, B2, T1, T2, M1, M2; M denotes the two-host **B+T** cages) first and six (B3, B4, T3, T4, M3, M4) approximately ten days later; ∼6,000 flies in total were used. Cages were maintained at census sizes of 1,000–2,000 flies with transfers every 18 days onto fresh plants. Single-host cages were provisioned exclusively with *B. vulgaris* or *T. glabra*. Two-host cages received equal numbers of both species, and larvae were managed so that approximately equal numbers of individuals were contributed by each host, satisfying the equal-contribution assumption of the Levene model. Populations were propagated for 10 generations in parallel under identical conditions.

### Plant growth and fly rearing

The host plants used for the single expansion generation before the start of the experiment were grown from purchased seed (*T. glabra*, Prairie Moon Nursery, Winona, MN; *B. vulgaris*, Garden Medicinals, Mineral, VA). All plants used during the evolution experiment were grown from seed collected from the wild source populations. Seeds were surface-sterilized with bleach and cold-stratified at 4 °C on MS medium before germination. Germinated seedlings were grown in Jiffy-7 peat pellets in 6-cm divided cells in climate-controlled rooms in the BioSciences West building at the University of Arizona under a 16 h/8 h light cycle at 21 °C/15 °C and approximately 50% relative humidity. Light was supplied by F40T12 Cool White Supreme fluorescent tubes (Philips Lighting, Somerset, NJ) at approximately 100 µmol m⁻² s⁻¹ measured at soil level. Plants were watered from the bottom to saturation three times per week with 15-16-17 fertilizer at 130 ppm nitrogen (1 mL granules per liter of tap water). Plants did not flower under these conditions and were used 6–8 weeks after germination. Flies were reared under the same conditions except at 22–23 °C day and 20 °C night. Adults were collected daily on emergence with an aspirator, anaesthetized with CO₂, and held in 50-mL vials provisioned with one cotton ball soaked in spring water and a second soaked in 10% honey solution. Once emergence was complete, flies were released into a mesh cage and aspirated in sets of 25 to synchronize mating and to reduce the post-eclosion die-offs seen when flies were left in vials. For each population, 20 sets (500 flies) were selected at random to propagate the next generation, which avoided selection on development time; remaining flies were preserved at −20 °C in 100% ethanol for sequencing. The 500 flies were divided between two 16 × 16 × 24 inch (41 × 41 × 61 cm) fine-mesh cages with a clear vinyl top panel to avoid crowding. Each oviposition cage was provisioned with 12 plants and three dishes holding cotton balls soaked in 10% honey solution. After five days, when approximately 70% of flies were feeding actively, these plants were discarded and replaced with 16 fresh plants, which were added in the afternoon and removed the following morning, giving 18 h of exposure. The infested plants were then transferred to cages without flies, placed in trays and watered as needed through larval development and pupation, and the cycle was repeated each generation.

### Host-plant defense chemistry

Eight *B. vulgaris* and eight *T. glabra* leaves, each from a separate representative experimental plant, were lyophilized, shipped to Copenhagen, re-lyophilized, weighed and extracted by three 1-min rounds of boiling MeOH–H_2_O (7:3). Crude extracts were loaded onto anion-exchange mini columns and desulfated on-column overnight, with benzylglucosinolate as external standard, following established protocols (*105*). Desulfoglucosinolates were compared and quantified by HPLC-UV and identities confirmed by HPLC-ion trap-MS/MS for *B. vulgaris* and by UHPLC-UV-QToF-MS/MS for *T. glabra*; identification of all major glucosinolates was backed up by parallel analysis of authentic, NMR-verified reference compounds isolated from other accessions of *B. vulgaris* (*105*, *106*) or *T. glabra* (*107*). Full profiles, chromatograms, and per-plant levels are in the Supplementary Text (Figs. S2, S3; Tables S1–S4).

### Population viability and realized heritability

Total numbers of flies surviving from egg to adult were counted per cage at generations 2, 3 and 10. Generation-10 counts were increased by 200 flies per cage, the number of adults removed from each cage for pool sequencing. Relative population viability was defined as the log-ratio of survivor counts in each single-host cage to the geometric mean of the **B+T** cages in the same generation. Generations 2 and 3 were grouped as the pre-selection state (logistically it was not possible to complete the phenotyping only in Generation 2) and generation 10 as the post-selection state. The selection differential was the deviation of each single-host population’s mean log-ratio at the pre-selection state from the grand mean across all populations, and the response was the change in mean log-ratio between states. A realized heritability-like response coefficient was estimated from a per-generation form of the breeder’s equation for 7 and 8 elapsed generations, giving a range. This statistic summarizes how strongly the population-level phenotype moved in the direction predicted by the selection differential.

### Reciprocal performance assay

Tenth-generation F1 adults from six experimental cages (two per treatment) were reared for one generation on *B. vulgaris*, *T. glabra*, or a 1:1 mixture, to minimize maternal environmental effects. Sixty flies (1:1 sex ratio) per rearing cage were introduced into phenotyping cages containing six plants (six *B. vulgaris*, six *T. glabra*, or three of each). Flies mated for five days, plants were removed, fresh plants were provided for 24 h of oviposition, and each plant was then bagged separately for larval development in a 36-cell tray with one plant per replicate × test-environment combination. Emerging flies were counted daily for 10 days (**Fig. S6**). Counts were normalized by the number of ovipositing females (*n* = 30) and modelled with a negative binomial GLM (glm.nb, MASS) including evolution treatment, test environment, a match indicator, and an offset for the number of plants. Hypotheses were tested as linear contrasts on model coefficients (emmeans, multcomp): home-court advantage (matched versus mismatched), specialist trade-off (the 2 × 2 single-host matrix), generalist jack-of-all-trades (one-sided interaction contrast of **B+T** against the specialists across all test environments), and generalist innovation (**B+T** versus specialists in [**B+T**]_test_). Modelled and empirical means are in **Table S5**.

### Levene model parameterization, bootstrap and invasion analysis

Fitness was parameterized with selection coefficients *s*_B_ and *s*_T_ under an additive model, and the equilibrium condition of the two-niche Levene model was solved numerically across a grid of niche contribution *c* and selection asymmetry. Empirical parameters were obtained by 1,000 non-parametric bootstrap iterations: in each iteration the assay data were resampled with replacement, refit with the negative binomial GLM, and fitnesses predicted for each genotype class in each test environment. The effective niche size was the proportion of total productivity contributed by the **B** environment and selection strength was the relative advantage of the favored homozygote over the disfavored homozygote in that niche. Stability was assessed by invasion analysis, computing both invasion indices per iteration and recording the fraction of iterations in which both exceeded one (**Fig. S5**). To express the assay in genotypic terms, the GLM was refit treating **B**-evolved as *AA*, **T**-evolved as *aa*, and [**B+T**]-evolved as *Aa* at *p* ≈ 0.5, yielding per-host *s* and *h* and the additive protection equilibrium *q*\*_phenotype_ = *s*_B_/(*s*_B_ + *s*_T_).

### Pooled and individual whole-genome sequencing

DNA was extracted and sequenced from experimental and wild populations in three efforts. In 2015 (University of Arizona), DNA from pools of 80 females (80 for founders; 22–60 per wild host-site sample) was prepared with the Kapa HyperPrep Kit and sequenced on an Illumina HiSeq 2500 with 100-bp paired-end reads; overlapping pairs were collapsed to ∼180-bp single-end equivalents. In 2016 (UC Berkeley), all 12 evolved populations at generation 10, six wild populations and pooled founders were prepared as indexed libraries in a multiplexed superpool and sequenced on a HiSeq 4000 (100-bp paired-end, 350–400-bp inserts), with 75% of effort on the evolved pools. In 2022 (UC Berkeley and New York University), 114 pools spanning all experimental and wild populations were extracted with Qiagen DNeasy kits (40 flies per batch, pooled 1:1 to 80 females), quantified by Qubit, prepared with SeqWell plexWell LP 384 in three independent libraries per pool, and sequenced on Illumina NovaSeq. Per-pool sequencing and mapping summaries are in **Table S8**. Separately, 128 founder flies and 64 T2G07 evolved flies were individually sequenced to ∼0.5× coverage for genotype-likelihood analyses.

### Read processing, mapping and variant calling

Reads were processed and aligned against the *S. flava* reference assembly (sfla_v2) with the grenepipe Snakemake pipeline (*108*). Adapters and low-quality bases were removed with fastp v0.20.1 (*109*) (paired-end adapter detection; poly-G trimming with minimum length 10; 5′ and 3′ sliding-window trimming of 4 bp at mean quality 25 and 20; minimum read length 60 bp; first 6 and 8 bases trimmed from reads 1 and 2). Reads were mapped with BWA-MEM v0.7.17 (*110*), replicate BAMs merged with samtools, alignments with mapping quality below 20 removed, duplicates marked and removed with Picard MarkDuplicates v2.27.5, and indels locally realigned with GATK v3 (*111*). Pooled allele frequencies were called with PoolSNP (*112*, *113*) from samtools mpileup summaries. Because depth, pooling structure and biological sampling differed across the study, separate PoolSNP panels were called for founder pools (max-cov 0.99, min-cov 100, min-count 50, min-freq 0.10, miss-frac 0), wild pools (0.98, 20, 5, 0.05, 0), the generation 1–9 time series (0.98, 30, 10, 0.01, 0), and the generation 10 pools (same nominal thresholds, run independently because replicate merging altered the depth distribution). Only biallelic SNPs were retained. Effective sample sizes (*N*_eff_) were estimated per pool and site from census pool sizes and PoolSNP output and used wherever an analysis required an effective number of chromosomes rather than raw read depth. Sites were then filtered to those polymorphic (0.01 < *f* < 0.99) in at least half of the sampled generations within a replicate and supported by at least 12 population × treatment replicates, and the panels were intersected to a single canonical coordinate set of 812,670 biallelic SNPs used for all comparative analyses. Software versions and parameters are listed in **Table S9**.

### Muller-element assignment

The *S. flava* v2 proteome (12,254 proteins from 12,365 gene records) was aligned against the *D. melanogaster* reference proteome (UniProt UP000000803) with DIAMOND blastp v2.1.10 (*114*) in --more-sensitive mode, retaining the top hit per query at *e* ≤ 10^−5^ (92% of queries hit). *D. melanogaster* chromosomes were converted to Muller elements (X→A, 2L→B, 2R→C, 3L→D, 3R→E, 4→F). Scaffolds were classified as confident (≥ 10 resolved genes and ≥ 70% top-Muller agreement) or ambiguous (≥ 5 and ≥ 50%). Scaffold 597 failed both thresholds because its resolved genes split 49%/47% between Muller C and D, consistent with a 2R + 3L fusion or an assembly mis-join (**Table S10**) (*115*).

### Founder population structure

Low-coverage reads from the 128 individually sequenced founders were mapped as above, and genotype likelihoods estimated with ANGSD v0.940 (*116*) (-GL 1, -SNP_pval 1e-6, minimum base and mapping quality 20, ≥ 64 individuals with data, minimum minor-allele frequency 0.01). Population structure was inferred with PCAngsd (*117*) (-e 10 -iter 500 -maf 0.05 -admix - admix_K) for *K* = 2 to 8, and the first admixture component compared between founders of *B* and *T* host origin (**Fig. S12**).

### Allele-frequency trajectory PCA

Per-sample allele frequencies were taken from the AD field of the joint VCF across the 22 pool-seq libraries spanning the time series. Sites were retained at per-sample depth between 10 and 500 in all samples and mean frequency in [0.01, 0.99]. Each SNP row was centered on its across-sample mean and scaled by √(*p*(1 − *p*)) (*118*), and the sample × SNP matrix decomposed with a full SVD, retaining ten components. Per-treatment *t*_10_ centroids were computed in PC1–PC10 space, and the displacement of the **B+T** centroid from the midpoint of the **B** and **T** centroids was measured as a Euclidean distance *d* and an angle θ from the *B*–*T* axis. Significance was assessed by 10^4^ treatment-label permutations, repartitioning the 12 *t*_10_ cages into three groups of four and recomputing *d*.

### Genetic diversity

θ_π_, Watterson’s θ_W_ and Tajima’s *D* were computed per pool in non-overlapping windows with grenedalf v0.6.2 (*119*), which applies pool-seq normalization, using per-sample filters of minimum allele count 2, minimum depth 4 and maximum depth 500, and pool sizes of 50 (wild), 80 (founder) and 160 (generation 10; two independent 80-fly pools per cage). Per-replicate trajectories across generations 1–9 were computed in 200-kb windows restricted to the 1,192 windows common to all twelve cages (**Fig. S15**); founder and generation-10 endpoints were excluded from the trajectory analysis because they were sequenced to different mean depth. Treatment contrasts used a linear mixed-effects model of window medians with treatment × generation as fixed effects and per-replicate random intercepts and slopes. Between-replicate idiosyncrasy was quantified as the per-window standard deviation of per-pool segregating-site counts across the four replicates of a treatment, computed in 100-kb windows on scaffold 110 (the largest chromosome-arm-scale scaffold that is neither the fused scaffold 597 nor the candidate scaffold 439) and compared across treatments by one-sided Mann–Whitney *U* tests.

### Technical noise, effective population size and temporal covariance

Pool-sequencing overdispersion was measured directly by comparing individual pseudo-pool allele frequencies from the individually sequenced founders against the founder pool-seq libraries, giving an effective pool size of 29 diploids (58 haploid chromosomes) against a nominal 80 (**Fig. S18**). Effective population size was estimated three ways: from the among-replicate normalized variance of allele frequency (*F*) regressed on generation, from within-replicate temporal estimators implemented in poolSeq (*120–122*), and from a method-of-moments aggregate; raw and noise-corrected values are reported in **Table S7** and **Figs. S14, S16, S17**. Genome-wide linked-selection intensity was estimated with the temporal-covariance framework of Buffalo and Coop (*68*), computing per-treatment temporal covariance matrices after applying the overdispersion correction and summarizing the fraction of allele-frequency change attributable to selection, *G*, from off-diagonal covariances at lag *k* ≥ 2; significance was assessed by one-sided permutation (**Figs. S19–S23**).

### Haplotype blocks and linkage disequilibrium

Haplotype blocks were detected from the pool-seq time series with haplovalidate (*54*). Candidate SNPs were identified with the drift- and pool-noise-corrected Cochran–Mantel–Haenszel test in ACER (*123*), calibrated by scanning *N*_e_ from 100 to 750 at fixed effective pool size 58; *N*_e_ = 250 produced the null *p*-value distribution closest to uniform and was used throughout. Blocks were clustered on the four largest scaffolds (204 Mb total) with minimum founder minor-allele frequency 0.05, minimum coverage 20 in all 28 per-treatment samples, and a cap of 15,000 candidates per scaffold, across three single-treatment and three pairwise runs (**Figs. S10, S24**). Linkage disequilibrium was measured independently from genotype likelihoods with ngsLD (*53*) on the 128 founders and 64 T2G07 individuals, restricted to SNPs passing the ANGSD filter in both cohorts (**Figs. S8, S9**).

### Per-SNP GLMM and parametric-bootstrap LRT

For each SNP, allele counts were reconstructed as successes = AF × *N*_eff_ and totals = *N*_eff_, with three pre-filters (every retained sample at total ≥ 15, ≥ 6 samples with successes > 0, ≥ 3 distinct success/total ratios). Null and full binomial GLMMs were fit in lme4 (*124*) with a logit link, weights equal to total depth, a random intercept by replicate cage, generation as a continuous covariate and treatment as a two-level factor; the full model added the generation × treatment interaction. Each likelihood-ratio statistic was assigned a *p*-value by 200 parametric-bootstrap simulations from the fitted null, retaining only SNPs for which more than half of simulations converged for both models. Of 813,431 SNPs tested, 812,590 (99.9%) returned a usable statistic.

### Multi-test vote scan and signal-block definition

Five tests were combined per 200-kb non-overlapping window across the four chromosome-scale scaffolds: (1) the most-negative tail of cov(Δ*p̅*_B_, Δ*p̅*_T_) across SNPs within a window (minimum 50 SNPs); (2) the *z*-standardized slope of grenedalf temporal *F*_ST_(**B**, **T**) across *t*_1_–*t*_10_, fit per window by a linear mixed-effects model with a per-pair random intercept; (3) the empirical *p*-value of the observed **B** versus **T** Hudson *F*_ST_ at *t*_10_ among all 70 four-versus-four re-labellings of the eight *t*_10_ pools, computed with pool-seq-corrected effective chromosome counts (*N*_eff_ = 58); (4) the per-window mean of the per-SNP generation × treatment LRT statistic; and (5) the per-window maximum of the per-SNP BayPass *C*_2_ contrast statistic, taken across all three treatment contrasts (**B** versus **T**, **B** versus **B+T**, and **T** versus **B+T**), among the twelve *t*_10_ pools, conditional on the among-pool covariance matrix estimated within the same run. This BayPass run used the twelve *t*_10_ pools, four per treatment, at a haploid pool size of 160 each, across 813,431 markers without thinning; core, covariate and contrast models were each fit under BayPass v3.1 default MCMC settings. Each test was thresholded at the genome-wide top 5% (or *p* ≤ 0.05 for the permutation *F*_ST_) and calibrated by 10^4^ permutations stratified on per-window SNP density. Haplovalidate block tracks are shown as orthogonal evidence but do not contribute to the vote. A single window, chr_ScDA7r2_439:2,800,000–3,000,000 on the X chromosome, reached the genome-wide maximum of 4/5; the remaining test, the temporal *F*_ST_ slope, placed it at the 84.5th percentile. This window is referred to as the signal block, and the surrounding ngsLD-defined founder block (scaffold 439:2,640,000–3,610,000) as the LD block. Gene content was annotated by extracting CDS records for scaffold 439:2.50–3.75 Mb, translating them, and assigning *D. melanogaster* orthologs by DIAMOND blastp top hit at *e* ≤ 10^−5^.

### Equilibrium frequency estimation from allele-frequency dynamics

Per-SNP logit slopes were fit for **B**, **T** and **B+T** over cumulative *t*_1_→*t*_max_ windows for *t*_max_ = 2 to 10. The median *s*_B_ and *s*_T_ across the 339 top-LRT SNPs in the LD block were substituted into the additive Levene equilibrium *q*\*_genomic_ = *s*_B_/(*s*_B_ + |*s*_T_|). Generations at which the median *s*_T_ was non-negative give a degenerate equilibrium and are reported as not determined. As an independent check, selection coefficients were estimated within the same block using a temporal least-squares estimator (*66*) and substituted into the same expression.

### Wild host-associated differentiation

Pairwise *F*_ST_ among the six wild pools was computed in 390-kb windows and summarized by pair category (**Fig. S11**). Because between-site structure was at least as large as within-site between-host structure, all per-SNP tests were conditioned on the genome-wide among-pool covariance matrix Ω. BayPass (*51*, *52*) was run on a thinned SNP set (every 16th site) with default MCMC settings (20 pilot runs of 500 iterations; 5,000 burn-in iterations; 1,000 post-burn-in samples thinned every 20 iterations) to estimate the 6 × 6 scaled Ω, and the fit was checked against the χ² expectation for the per-SNP differentiation statistic. Conditional on Ω, a binary host covariate (*B* = +1, *T* = −1) was regressed against the full 812,670 SNPs to obtain per-SNP standardized effects and Bayes factors, and a contrast model was fit treating host as a discrete grouping to obtain the *C*_2_ statistic for the wild host contrast (**Fig. S25**).

### Laboratory-to-wild sign concordance

Starting from the per-SNP GLMM table, three filters were applied: top 50% of the genome-wide generation × treatment LRT distribution; |*z*| ≥ 2 on both *s*_B_ and *s*_T_, where *z* is the per-replicate logit-slope estimate divided by its standard error; and founder minor-allele frequency in [0.05, 0.95]. This gave 4,050 background SNPs, of which 3,979 fell on Muller elements A–E (709 on A, 695 B, 786 C, 790 D, 999 E); the remainder were on Muller F or lacked confident Muller element assignment and were omitted from per-element comparisons. For each SNP, lab Δ*p* was the mean across the four replicate pairs of AF(*B*_r_, *t*_10_) − AF(*T*_r_, *t*_10_), and wild Δ*p* the mean across the three site pairs of AF(site *B*) − AF(site *T*); both are unpolarized alternate-allele codings. Concordance was the mean of the indicator 1[sign(lab Δ*p*) = sign(wild Δ*p*)], with ties excluded and binomial standard errors included. Null distributions were built per Muller element by 1,000 bootstrap iterations in which each candidate SNP was matched to a randomly drawn background SNP on the same element whose founder minor-allele frequency fell within ± 0.05. Lab and wild Δ*p* were additionally compared by Spearman correlation and least-squares regression, separately for the genome-wide background and for the signal block.

### Statistics and reproducibility

The experiment comprised four independent replicate cage populations per treatment, founded and propagated independently. No statistical method was used to predetermine sample size; no data were excluded except as specified by the filters above; cages were assigned to treatments by balanced allocation from the ten founder cages rather than by randomization, and investigators were not blinded. Tests are two-sided except for contrasts whose direction was predicted in advance, each identified as one-sided where it is reported: the population-viability ratios of each single-host treatment relative to **B+T**; the *t*_10_ allele-frequency contrasts of each single-host treatment against **B+T** at the signal block, where the Levene model predicts that the locally favored allele rises under single-host selection relative to two-host selection and where the favored allele was assigned by leave-one-replicate-out cross-validation; the generalist “jack-of-all-trades” contrast, which asks only whether **B+T** exceeds the specialists averaged across test environments; the permutation tests of *G*, which is bounded below by zero under the null hypothesis of no selection; and the comparison of between-replicate variability in segregating-site density, where subdivided selection is predicted to retain more idiosyncratic variation than uniform selection. Permutation and bootstrap procedures used 10^3^–10^4^ iterations per analysis, except the per-SNP parametric-bootstrap likelihood-ratio tests, which used 200 per SNP.

## Data availability

Raw sequencing reads for all pooled and individually sequenced libraries have been deposited in the NCBI Sequence Read Archive under BioProject PRJNA1535502. The *S. flava* sfla_v2 reference assembly is available from GenBank under accession GCA_030179655.1. Processed allele-frequency matrices, per-window diversity and *F*_ST_ tables, the per-SNP GLMM output, the phenotypic performance data, and the sfla_v2 genome annotation are archived at Zenodo (https://doi.org/10.5281/zenodo.22983766). Source data for all main-text figures are included in the Zenodo archive.

## Code availability

All analysis code, including the grenepipe configuration, variant-filtering scripts, and the R and Python scripts implementing the Levene bootstrap, temporal-covariance analysis, GLMM scan, vote scan, equilibrium estimation and lab-to-wild concordance test, is at https://github.com/DilerHaji/scaptomyza_evolve.

## Acknowledgements

We thank Julie Ray, Mitchell Feldmann, Alana Augur, Amelia White, Soraya Sandoval, and Elijah Sensibar for help with plant and fly rearing throughout the experiment. We are grateful to Julie Johnson for creating the fly and plant illustrations used in Figure 1. We thank members of the Whiteman laboratory, including Sarah Lai, Julianne Peláez, Nicolas Alexandre, Kirsten Verster, Hiromu Suzuki, Samridhi Chaturvedi, Moe Bakhtiari, Rebecca Tarnopol, Jessica Aguilar, and Jeff Groh, for discussion, feedback, and support throughout the project. We thank Priya Moorjani, Rebecca Tarvin, and Benjamin Blackman for serving on D.H.’s dissertation committee, reading the manuscript in full, and providing extensive feedback that improved the framing, interpretation, and presentation of this work. We also thank Peter Sudmant, Guy Sella, Chris Simon, Moi Exposito-Alonso, and the UC Berkeley Center for Theoretical and Evolutionary Genetics community for helpful discussions that improved the population-genomic framing and interpretation of the experimental evolution results. We thank the UC Berkeley Savio/High Performance Computing support team for computational infrastructure and technical support, and the NYU, UC Berkeley, and University of Arizona sequencing facilities for sequencing support. Anthropic’s Claude (Opus 5 model) was used during the preparation of this work to assist with: organizing and documenting analysis code and troubleshooting computational environments; verifying the consistency of figures, tables, and in-text citations against their sources and against the underlying data; managing reference and figure numbering; copy-editing; and literature search. All AI-assisted output was checked against the primary data and the cited literature by the authors, who reviewed and take full responsibility for the content of this work. This work was supported by funding from the National Science Foundation Doctoral Dissertation Improvement Grant DEB-1405966 to A.D.G. and N.K.W., the National Institutes of Health/National Institute of General Medical Sciences (award no. R35GM119816) to N.K.W., the National Science Foundation Graduate Research Fellowship Program to D.H., a Berkeley Fellowship to D.H., a UC Dissertation-Year Fellowship/Dissertation Completion Award to D.H., a Society for the Study of Evolution R. C. Lewontin Early Award to D.H., and financial support from the Department of Integrative Biology at the University of California, Berkeley to D.H.

## Author contributions

D.H., A.D.G. and N.K.W. designed the study. A.D.G. and N.K.W. established the field collections and the founder colony. A.D.G. and E.R.L. maintained the experimental evolution populations and performed the performance assays. N.A. performed the glucosinolate chemistry and interpreted the defense profiles. D.H. performed sequencing and analysis. J.F.A. and J.B. advised on experimental design and population-genomic inference. D.H. wrote the manuscript with input from all authors. N.K.W. and A.D.G. supervised the project and acquired funding. All authors read and approved the final manuscript.

## Additional information

Supplementary Information is available for this paper. Correspondence and requests for materials should be addressed to D.H., A.D.G. or N.K.W.

## Supplementary Information

Supplementary Information is provided as a single file. It contains the Supplementary Text, Supplementary Figures S1–S25, Supplementary Tables S1–S11 and the Supplementary Literature Cited, covering: experimental design and founder establishment (Fig. S1); host-plant defense chemistry (Figs. S2, S3; Tables S1–S4); reciprocal-assay power (Fig. S4; Table S5); developmental timing (Fig. S6); Levene stability sensitivity analysis (Fig. S5); founder population structure (Fig. S12); genetic diversity across wild, founder and evolved pools (Figs. S13, S15); pool-sequencing overdispersion and effective population size (Figs. S14, S16–S18; Tables S7, S11); temporal covariance and the *G* statistic (Figs. S19–S23); the genome-wide **B** versus **T** correlation (Fig. S7); haplotype blocks and linkage disequilibrium (Figs. S8–S10, S24); wild host-associated differentiation (Figs. S11, S25; Table S6); Muller-element assignment (Table S10); and sequencing and software summaries (Tables S8, S9).

