## Supplemental Materials for "Multiple host species maintain genetic diversity in herbivore populations"

This file contains Supplementary Figures S1–S25, Supplementary Tables S1–S11, the Supplementary Text, and the Supplementary Literature Cited.

### Supplementary Text

### Supplementary Figure Index

| Figure | Description | Located in |
|---|---|---|
| S1 | F3 founder population establishment | Experimental evolution design |
| S2 | HPLC-UV quantification and comparison of glucosinolates in *B. vulgaris* and *T. glabra* | Defense chemistry of *B. vulgaris* and *T. glabra* host plants |
| S3 | HPLC-ion trap-MS/MS and UHPLC-UV-QToF-MS/MS chromatograms for *T. glabra* and *B. vulgaris* | Defense chemistry of *B. vulgaris* and *T. glabra* host plants |
| S4 | Reciprocal assay power and per-plant adult production | Resolution of the two fitness measurements |
| S5 | Sensitivity analysis of Levene's stability conditions | Equilibrium dynamics from fitness differentials |
| S6 | Developmental timing across test environments | Test of reciprocal fitness trade-offs in post-evolution experimental flies |
| S7 | Genome-wide BvT correlation distribution | Genome-wide contribution of selection to allele frequency change |
| S8 | ngsLD r2 hotspot vs flanking | Linkage disequilibrium (LD) using individually-sequenced experimental flies |
| S9 | Genome-wide windowed LD: founders vs T2G07 | Linkage disequilibrium (LD) using individually-sequenced experimental flies |
| S10 | Haplotype-block sizes per treatment | Haplotype structure and scale of linked selection |
| S11 | Wild-pool pairwise FST heatmap (six pools) | Wild host-associated differentiation in *Scaptomyza flava* |
| S12 | PCAngsd population structure of 128 individually-sequenced founders | Population structure of experimental founder flies |
| S13 | Per-pool genetic diversity boxplots (wild / founder / G10) | Standing variation, colony bottleneck, and diversity rebound |
| S14 | Mean depth + F-regression noise correction for Ne | Drift variance and effective population size |
| S15 | Per-replicate diversity trajectories across G1-G9 | Standing variation, colony bottleneck, and diversity rebound |
| S16 | poolSeq Ne at three pool-size corrections | Within-replicate temporal estimates of Ne |
| S17 | Summary of three independent Ne estimators | Within-replicate temporal estimates of Ne |
| S18 | Pool sequencing overdispersion (per-fly vs pool-seq, between-pool variance) | Technical noise |
| S19 | Temporal-replicate covariance matrix per treatment | Genome-wide contribution of selection to allele frequency change |
| S20 | Bias-correction diagnostic plots for the temporal covariance | Genome-wide contribution of selection to allele frequency change |
| S21 | Cumulative G(t) statistic per treatment | Genome-wide contribution of selection to allele frequency change |
| S22 | Convergence correlation, Npool=80 vs Neff=29 | Genome-wide contribution of selection to allele frequency change |
| S23 | Power to detect convergence correlation > 0 | Genome-wide contribution of selection to allele frequency change |

| Figure | Description | Located in |
| --- | --- | --- |
| S24 | Block-trajectory AUC (non-equilibrium dynamics by t10) | Haplotype structure and scale of linked selection |
| S25 | Wild BayPass scan for host-associated differentiation | Wild host-associated differentiation in *Scaptomyza flava* |

### Sampling and experimental design

We designed the experimental evolution study to test the prediction from population genetic theory that environmental heterogeneity can maintain fitness-associated genetic variation within populations. Under the Levene model, polymorphism is maintained when alternative alleles are favored in different habitats, provided that mating occurs randomly across habitats and juvenile densities on any given habitat are approximately equal. Our study system, *Scaptomyza flava*, is an oligophagous leaf-mining drosophilid that develops on living mustard plants. In the fallow fields we sampled in New England, *S. flava* primarily attacked two sympatric host species, *Barbarea vulgaris* and *Turritis glabra*, which we showed differ in defense chemistry (see below).

#### Wild sampling

In the summer of 2014, wild larvae were collected from both *B. vulgaris* and *T. glabra* growing in Newington, New Hampshire, a site that yielded the largest number of emerging flies among surveyed localities. We collected 130 adults (58 adults from *B. vulgaris* and 72 adults from *T. glabra*) by rearing them from bagged, field-collected leaves. These flies were interbred in the laboratory to establish an outbred colony. To minimize inbreeding and preserve natural genetic diversity, this colony was maintained for one generation on *Arabidopsis thaliana* (Col-0) and then for one generation on a mixture of *B. vulgaris* and *T. glabra*. During this period, seeds of *B. vulgaris* and *T. glabra* were collected from the same wild populations and propagated in an indoor plant growth chamber to generate a bank of experimental plants. The outbred fly colony was expanded to ~15,000 individuals before subdivision into founder cages containing these experimental plants.

#### Experimental evolution design

In Fall 2014, the colony was split across nine mesh cages provisioned with experimental host plants over a two-day oviposition schedule designed to minimize bias from host preference. Four cages received *B. vulgaris* on the first day and *T. glabra* on the second, four cages received *T. glabra* followed by *B. vulgaris*, and one cage received a mixture of both host species. After a two-day oviposition period, plants containing larvae were redistributed across ten new cages so that emerging adults were separated by host of origin while retaining the oviposition rotation design. These ten founder cages comprised pooled offspring from pairs of the original nine cages (T1+T2, T3+T4, B1+B7, B2+B8, B3+B4, B5+B6, and so on), along with a mixture cage created by aspirating equal numbers of flies from all nine. Together these cages contained the full genetic base for experimental evolution with minimal evidence of population genetic structure due to the population expansion procedure outlined above (See "Population structure of experimental founder flies").

To initiate a set of replicate experimental populations, each of the ten founder cages was split into batches of 25 flies. Each new experimental cage was founded with two batches (50 flies) from each of the ten founder cages, producing genetically balanced founding populations of 500 flies per cage. This allocation was performed twice on separate dates, such that six populations (B1, B2, T1, T2, M1, M2; M denotes the two-host **B+T** cages) were founded first and six (B3, B4, T3, T4, M3, M4) were founded approximately ten days later. In total, ~6,000 flies drawn from more than 8,000 in the founder cages were used to establish the 12 experimental populations. Cages were maintained at census sizes of 1,000-2,000 flies to minimize drift, with transfers every 18 days onto fresh plants. In monoculture treatments, cages were provisioned exclusively with either *B. vulgaris* or *T. glabra*. In the mixed-host treatments, equal numbers of *B. vulgaris* and *T. glabra* plants were supplied, and larvae were managed such that approximately equal numbers of individuals contributed from each host, thereby maintaining the assumptions of the Levene model. Populations were propagated for 10 generations in parallel under identical environmental conditions.

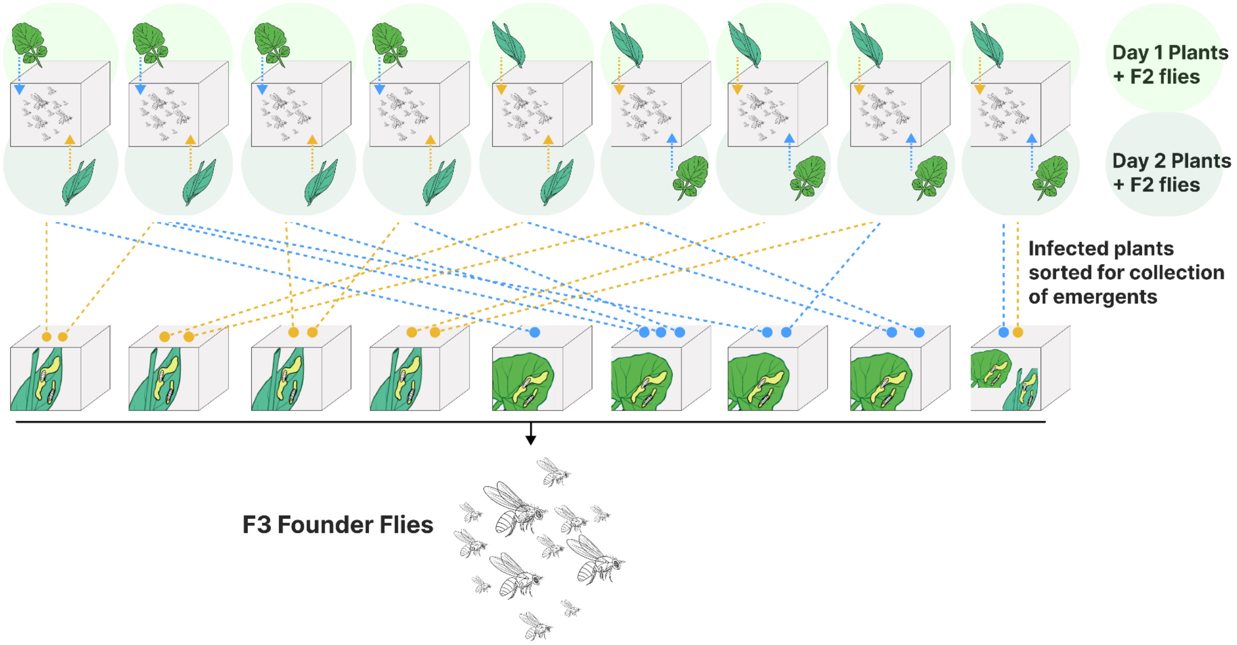

Figure S1: Detailed visualization of the establishment of the F3 founder population of *S. flava* for experimental evolution. To minimize bias from host preference and maximize retention of standing genetic diversity, the expanded outbred colony was distributed across nine oviposition cages (top row). These cages were provided with pots of *B. vulgaris* (round leaves, blue arrows) and *T. glabra* (long leaves, yellow arrows) rosettes on an alternating two-day schedule. Infected plants containing larvae were redistributed into ten rearing cages following oviposition (bottom row). We indicated this sorting process by the dotted lines, separating emerging flies by their host of origin while pooling offspring from the original cages. We collected adults emerging from these rearing cages to form the F3 founder flies, which served as the genetically balanced source population for the twelve experimental evolution replicates in this study.

### Defense chemistry of *B. vulgaris* and *T. glabra* host plants

Glucosinolates are major chemical defenses in crucifers such as *B. vulgaris* and *T. glabra*. Biochemically, glucosinolates are thioglucosides with an aglycone that contains a variable side chain derived from a precursor amino acid[1]. In this study we discuss only glucosinolates derived from the amino acids homo-phenylalanine (homoPhe), tryptophan (Trp), and chain-elongated methionine (n-homoMet). Upon physical tissue damage, such as insect chewing, the plant enzyme myrosinase comes into contact with the glucosinolates and catalyzes their hydrolysis to unstable aglucones. The aglucones can fragment into several compound types depending on the presence or absence of specifier proteins. In the absence of specifier proteins they fragment into isothiocyanates, also known as mustard oils. Mustard oils exert various antinutritional effects through reaction with thiols and other nucleophiles, and their consumption depletes cellular glutathione[2]. Some glucosinolates carry a hydroxy group at the $\beta$-position of the side chain. In that case the isothiocyanate that forms first undergoes a rapid intramolecular reaction to give a cyclic oxazolidine-2-thione. Oxazolidine-2-thiones also appear to have antinutritional effects on insects[3], although the mechanism is less well understood.

The leaf glucosinolate profiles of European populations of *B. vulgaris* are well characterized, and only glucosinolates derived from homoPhe and Trp are reliably reported from the species[4–6]. Two major chemotypes are known[4]. The main Central and Western European chemotype, the "G-type", is dominated by (*S*)-2-hydroxy-2-phenylethylGSL, whose trivial name is glucobarbarin (BAR). Here and in the following "GSL" in a systematic name means "glucosinolate". This G-chemotype has previously been reported from introduced North American populations[7]. A second chemotype, found mainly in Eastern Europe, is dominated by the (*R*) epimer, epiglucobarbarin (EBAR). Rarer, mutant-like forms have also been described, often dominated by phenethylGSL (PE), the biosynthetic precursor of the glucobarbarins. All chemotypes also contain Trp-derived glucosinolates, typically indol-3-ylmethylGSL (IM) and its 4-methoxy derivative, abbreviated 4moIM.

#### Identification and quantification of glucosinolates in experimental plants

For glucosinolate analysis, we individually lyophilized eight *B. vulgaris* leaves ("1B" to "8B") and eight *T. glabra* leaves ("1T" to "8T"), each from a separate representative experimental plant, in our U.S. laboratory and shipped them by airmail to a Danish laboratory. The samples were received in September 2015 and processed within the same month. We lyophilized the samples again before weighing and extraction to obtain the true dry weight. We then extracted glucosinolates by three brief rounds of hot extraction, each of 1 min, in boiling MeOH–$H_2O$ (7:3). We extracted the entire sample, between 60 and 120 mg for *B. vulgaris* and between 70 and 110 mg for *T. glabra*, which lies within the usual linear range of the method. The remaining procedure converted the extracted glucosinolates to their desulfated derivatives for HPLC identification and quantification. We loaded each crude extract onto an anion-exchange mini column prepared as described previously[8]. In parallel, we applied standards of 535 nmol benzylGSL (BZ; Merck, Darmstadt, Germany) to separate mini columns. We carried out the column wash and the enzymatic overnight on-column desulfation exactly as described previously[8]. Finally, we eluted the resulting desulfoglucosinolates with 5 × 1.00 mL $H_2O$ and used the eluate for HPLC analysis.

**Comparative HPLC-UV identification of glucosinolates in *B. vulgaris* and *T. glabra***

Instruments for HPLC with UV detection were exactly as described previously[6], except that the LC-ABZ column was replaced with a Luna Phenylhexyl column (250 mm × 4.6 mm, 5 µm; Phenomenex, Torrance, CA) to ensure baseline separation of the desulfo derivatives of PE and 4moIM. We injected 100 µL per sample and operated the diode array detector at 200-400 nm for peak identification, recording at 229 nm for peak integration with 8 nm band width. A data-example for each species is shown as Table S1. Authentic BAR, BZ, IM, 4moIM, 7mSOh, 8mSOo, Buen and other references were run in parallel. For modified Brown-Windsor abbreviations of glucosinolates not detected in *B. vulgaris* or *T. glabra*, see reference [5]. We analyzed all eight *B. vulgaris* desulfoglucosinolate samples by HPLC-UV. The HPLC-UV chromatograms of desulfoglucosinolates revealed that the profile of glucosinolates from *T. glabra* contrasted sharply with the profile from *B. vulgaris* (Figure S2; Tables S2, S3). In fact, the two species had only one minor glucosinolate in common (4moIM) (Table S4). For quantification from UV peak areas we applied well-established approximate relative response factors. For *B. vulgaris* these were 0.95 for the external standard BZ and for the homoPhe-derived PE, BAR and EBAR; 0.29 for IM; and 0.25 for 4moIM [4]. For *T. glabra* we used the usual default relative response factor of 1.00 for all aliphatic desulfoglucosinolate peaks[9], 0.25 for 4moIM, and 0.95 for the external standard BZ. Peak identification did not rely solely on the HPLC-UV data (retention times and diode array UV spectra) but was concluded based on additional experiments as described below.

**Table S1.** HPLC-UV peak table for (A) a representative *T. glabra* leaf desulfoglucosinolate sample and (B) a representative *B. vulgaris* leaf desulfoglucosinolate sample. Retention time (min), integrated UV peak area at 229 nm (instrument units) and area percent are given for every peak resolved in a single chromatogram. Annotations give the glucosinolate assigned to each peak; unannotated peaks were not identified as specific desulfoglucosinolates. Some unannotated peaks may be due to impurities or random baseline fluctuations, other minor peaks may correspond to minor peaks quantified from extracted ion chromatograms in UHPLC-UV-QToF-MS/MS. Area percentages are of total integrated peak area and are not corrected by relative response factors, so they differ from the response-factor-corrected levels in Table S3. Abbreviations are defined in Table S4.

| **Peak #** | **Retention Time** | **Area** | **Area Percent** | **Annotation** |
|---|---|---|---|---|
| A (*T. glabra* sample 1T) | | | | |
| 1 | 4.565 | 6,866 | 0.10% | |
| 2 | 4.960 | 10,423 | 0.15% | |
| 3 | 5.557 | 1,377 | 0.02% | |
| 4 | 5.995 | 43,587 | 0.64% | |
| 5 | 6.197 | 43,564 | 0.64% | |
| 6 | 6.400 | 55,310 | 0.81% | |
| 7 | 6.741 | 137,348 | 2.00% | |
| 8 | 7.115 | 154,445 | 2.25% | |
| 9 | 8.981 | 111,526 | 1.63% | |
| 10 | 9.824 | 322,903 | 4.71% | 3o-5mSOp |
| 11 | 11.701 | 313,192 | 4.57% | |
| 12 | 12.224 | 86,027 | 1.26% | |

| Peak # | Retention Time | Area | Area Percent | Annotation |
|---|---|---|---|---|
| 13 | 12.661 | 3,545,696 | 51.75% | 3o-6m$SO_2$h |
| 14 | 13.205 | 333,536 | 4.87% | 3h-6m$SO_2$h |
| 15 | 14.080 | 17,539 | 0.26% | |
| 16 | 15.029 | 116,849 | 1.71% | 3o-7mSOh |
| 17 | 15.669 | 7,370 | 0.11% | |
| 18 | 16.619 | 253,402 | 3.70% | 3o-7m$SO_2$h |
| 19 | 17.856 | 231,528 | 3.38% | 3h-7m$SO_2$h |
| 20 | 19.456 | 2,995 | 0.04% | |
| 21 | 20.000 | 23,411 | 0.34% | |
| 22 | 21.077 | 175,699 | 2.56% | 6mSOh |
| 23 | 21.600 | 30,725 | 0.45% | |
| 24 | 22.571 | 517,226 | 7.55% | 6mSO2h |
| 25 | 24.128 | 2,005 | 0.03% | |
| 26 | 27.723 | 36,862 | 0.54% | 7mSOh |
| 27 | 29.195 | 115,384 | 1.68% | 7m$SO_2$h |
| 28 | 29.845 | 523 | 0.01% | |
| 29 | 37.237 | 117,745 | 1.72% | 4moIM |
| 30 | 40.160 | 9,706 | 0.14% | |
| 31 | 43.456 | 1,567 | 0.02% | |
| 32 | 44.485 | 9,789 | 0.14% | 6mSh |
| 33 | 52.981 | 15,313 | 0.22% | |
| Total | | 6,851,438 | 100.00% | |
| B (*B. vulgaris* sample 2B) | | | | |
| 1 | 4.843 | 12,216 | 0.24% | |
| 2 | 4.971 | 12,662 | 0.25% | |
| 3 | 5.877 | 6,390 | 0.13% | |
| 4 | 6.176 | 20,167 | 0.40% | |
| 5 | 7.019 | 53,680 | 1.06% | |
| 6 | 10.987 | 25,555 | 0.50% | |
| 7 | 11.467 | 5,353 | 0.11% | |
| 8 | 15.221 | 2,275 | 0.04% | |
| 9 | 18.315 | 4,513 | 0.09% | |
| 10 | 22.805 | 58,520 | 1.15% | EBAR |
| 11 | 24.544 | 3,773,963 | 74.37% | BAR |
| 12 | 26.773 | 1,540 | 0.03% | |

| Peak # | Retention Time | Area | Area Percent | Annotation |
|---|---|---|---|---|
| 13 | 30.709 | 976,529 | 19.24% | IM |
| 14 | 36.235 | 10,422 | 0.21% | PE |
| 15 | 36.608 | 83,595 | 1.65% | 4moIM |
| 16 | 39.584 | 13,924 | 0.27% | |
| 17 | 52.448 | 13,192 | 0.26% | |
| **Total** | | **5,074,496** | **100.00%** | |

**Table S2.** Glucosinolate levels, absolute and relative, in *B. vulgaris* leaves from growth-chamber grown plants as used in the experiments, originating from a wild population from Newington, New Hampshire (N = 8 plants). All reported glucosinolates were conclusively identified at phytochemical identification confidence level C, based on matching retention time, *m*/*z* and MS2 against authentic standards analyzed in parallel. Abbreviations and numbers follow ref.[5] and are defined in Table S4. Absolute levels are µmol $g^{-1}$ dry weight. Glucosinolates searched for by HPLC-ion trap-MS but not found: 4hBAR, 4hEBAR, 4hIM, 1moIM, BZ, any hydroxyBZ, 4moBZ, Pren, Buen, Peen, 4hBuen, 1mEt, 1mPr, 2mPr, n-butylGSL, any isomeric hydroxybutylGSL, any isomeric pentylGSL or hydroxypentylGSL, 3mPe or any other isomeric hexylGSL, 2h3mPe, 3hmPe or any other isomeric hydroxyhexylGSL, 3mSp, 3mSOp, 4mSb, 4mSbuen, 4mSOb, 5mSp, 5mSOp, 6mSh, 6mSOh, 6mSO$_2$h, 7mSh, 7mSOh, 7mSO$_2$h, 8mSo, 8mSOo, 8mSO$_2$o, or any hydroxy or oxo derivatives detected in *T. glabra*.

| Glucosinolate | Absolute, mean | SD | Relative (%), mean | SD |
|---|---|---|---|---|
| **HomoPhe-derived** | | | | |
| PE (**105**) | 0.02 | 0.01 | 0.3 | 0.3 |
| BAR (**40S**) | 11.3 | 7.7 | 89.7 | 1.5 |
| EBAR (**40R**) | 0.18 | 0.12 | 1.4 | 0.2 |
| **Subtotal** | **11.5** | **7.8** | **91.3** | **1.4** |
| **Trp-derived** | | | | |
| IM (**43**) | 1.00 | 0.87 | 7.6 | 1.7 |
| 4moIM (**48**) | 0.09 | 0.06 | 1.1 | 1.0 |
| **Subtotal** | **1.09** | **0.87** | **8.7** | **1.4** |
| **Total** | **12.6** | **8.7** | **100** | **–** |

**Table S3.** Glucosinolate levels, absolute and relative, in *T. glabra* leaves from growth-chamber grown plants as used in the experiments, originating from a wild population from Newington, New Hampshire (N = 8 plants). Names marked "tent." are tentative and denote hypothetical glucosinolates predicted from MS2, retention time, and the structures of co-occurring known glucosinolates. Levels in parentheses were not backed by MS2 because peak intensity was too low but were supported by retention time and *m/z*; "tr." denotes a trace. Abbreviations follow ref.[5], supplemented with 3h for 3-hydroxy and 3o for 3-oxo, and are defined in Table S4. Glucosinolate numbers follow refs. [1] and [10]. Absolute levels are μmol $g^{-1}$ dry weight. The aliphatic subtotal (8.86) plus 4moIM (0.04) gives the total (8.90). Glucosinolates searched for by HPLC-ion trap-MS but not found: PE, BAR, EBAR, IM, 4hIM, 1moIM, BZ, any hydroxyBZ or methoxyBZ, 1mEt, 1mPr, 2mPr or any other isomeric butylGSL, any isomeric hydroxybutylGSL, any isomeric pentylGSL or hydroxypentylGSL, 3mPe or any other isomeric hexylGSL, 2h3mPe, 3hmPe or any other isomeric hydroxyhexylGSL, Pren, Buen, 2hBuen, Peen, 3mSp, 3mSOp, 4mSb, 4mSbuen, 4mSO$_2$b, 8mSo, 8mSOo, 2h8mSOo.

| Glucosinolate | Absolute, mean | SD | Relative (%), mean | SD |
|---|---|---|---|---|
| **DihomoMet- to hexahomoMet-derived** | | | | |
| 4mSOb (**64**) | tr. | – | – | – |
| 5mSp (**94**) | (0.0007) | 0.0006 | (0.01) | 0.01 |
| 5mSOp (**72**) | 0.04 | 0.01 | 0.50 | 0.16 |
| 5mSO$_2$p ([**81**]) | 0.005 | 0.007 | 0.06 | 0.08 |
| 6mSh (**88**) | 0.01 | 0.005 | 0.15 | 0.05 |
| 6mSOh (**67**) | 0.24 | 0.09 | 2.7 | 1.0 |
| 6mSO$_2$h (**78**) | 0.93 | 0.73 | 10.6 | 9.4 |
| 7mSh (**87**) | (0.003) | 0.004 | (0.04) | 0.05 |
| 7mSOh (**66**) | 0.07 | 0.02 | 0.86 | 0.38 |
| 7mSO$_2$h (**160**) | 0.22 | 0.13 | 2.5 | 1.7 |
| 8mSO$_2$o (**80**) | tr. | – | – | – |
| **3-Hydroxy derivatives** | | | | |
| 3h-6mSO$_2$h (tent., [**34**]) | 0.45 | 0.16 | 4.9 | 0.9 |
| 3h-7mSO$_2$h (**161**) | 0.37 | 0.10 | 4.1 | 0.7 |
| **3-Oxo derivatives** | | | | |
| 3o-5mSO$_2$p (tent.) | 0.22 | 0.12 | 2.7 | 1.0 |
| 3o-6mSOh (tent.) | 0.37 | 0.20 | 4.2 | 2.1 |
| 3o-6mSO$_2$h (**162**) | 5.30 | 1.57 | 59 | 10 |
| 3o-7mSOh (tent.) | 0.17 | 0.04 | 1.9 | 0.2 |
| 3o-7mSO$_2$h (tent.) | 0.46 | 0.14 | 5.2 | 1.4 |
| **Subtotal, aliphatic** | **8.9** | **1.9** | **99.6** | **0.1** |
| **Trp-derived** | | | | |
| 4moIM (**48**) | 0.04 | 0.02 | 0.39 | 0.12 |
| **Total** | **8.9** | **2.0** | **100** | **–** |

**Table S4.** Comparison of the glucosinolates detected in *B. vulgaris* and *T. glabra*, with systematic names, common names, abbreviations and global glucosinolate numbers. Abbreviations are modified Brown-Windsor abbreviations[5], in which sulfanyls are abbreviated S, sulfinyls SO and sulfonyls $SO_2$, supplemented here with the prefixes 3h for 3-hydroxy and 3o for 3-oxo.

| Abbreviation | Systematic name | Common name | GSL no. |
|---|---|---|---|
| **External standard** | | | |
| BZ | BenzylGSL | Glucotropaeolin | **11** |
| **In both host species** | | | |
| 4moIM | 4-Methoxyindol-3-ylmethylGSL | 4-Methoxy glucobrassicin | **48** |
| ***Barbarea vulgaris* only** | | | |
| BAR | (*S*)-2-HydroxyphenethylGSL | Glucobarbarin | **40S** |
| EBAR | (*R*)-2-HydroxyphenethylGSL | Epiglucobarbarin | **40R** |
| PE | PhenethylGSL | Gluconasturtiin | **105** |
| IM | Indol-3-ylmethylGSL | Glucobrassicin | **43** |
| ***Turritis glabra* only** | | | |
| 4mSOb | 4-(Methylsulfinyl)butylGSL | Glucoraphanin | **64** |
| 5mSp | 5-(Methylsulfanyl)pentylGSL | Glucoberteroin | **94** |
| 5mSOp | 5-(Methylsulfinyl)pentylGSL | Glucoalyssin | **72** |
| 5mSO$_2$p | 5-(Methylsulfonyl)pentylGSL | – | **[81]** |
| 6mSh | 6-(Methylsulfanyl)hexylGSL | Glucolesquerellin | **88** |
| 6mSOh | 6-(Methylsulfinyl)hexylGSL | Glucohesperin | **67** |
| 6mSO$_2$h | 6-(Methylsulfonyl)hexylGSL | – | **78** |
| 7mSh | 7-(Methylsulfanyl)heptylGSL | – | **87** |
| 7mSOh | 7-(Methylsulfinyl)heptylGSL | – | **66** |
| 7mSO$_2$h | 7-(Methylsulfonyl)heptylGSL | – | **160** |
| 8mSO$_2$o | 8-(Methylsulfonyl)octylGSL | – | **80** |
| 3h-6mSO$_2$h | 3-Hydroxy-6-(methylsulfonyl)hexylGSL | – | tent., **[34]** |
| 3h-7mSO$_2$h | 3-Hydroxy-7-(methylsulfonyl)heptylGSL | – | **161** |
| 3o-5mSO$_2$p | 5-(Methylsulfonyl)-3-oxopentylGSL | – | tent. |
| 3o-6mSOh | 6-(Methylsulfinyl)-3-oxohexylGSL | – | tent. |
| 3o-6mSO$_2$h | 6-(Methylsulfonyl)-3-oxohexylGSL | – | **162** |
| 3o-7mSOh | 7-(Methylsulfinyl)-3-oxoheptylGSL | – | tent. |
| 3o-7mSO$_2$h | 7-(Methylsulfonyl)-3-oxoheptylGSL | – | tent. |

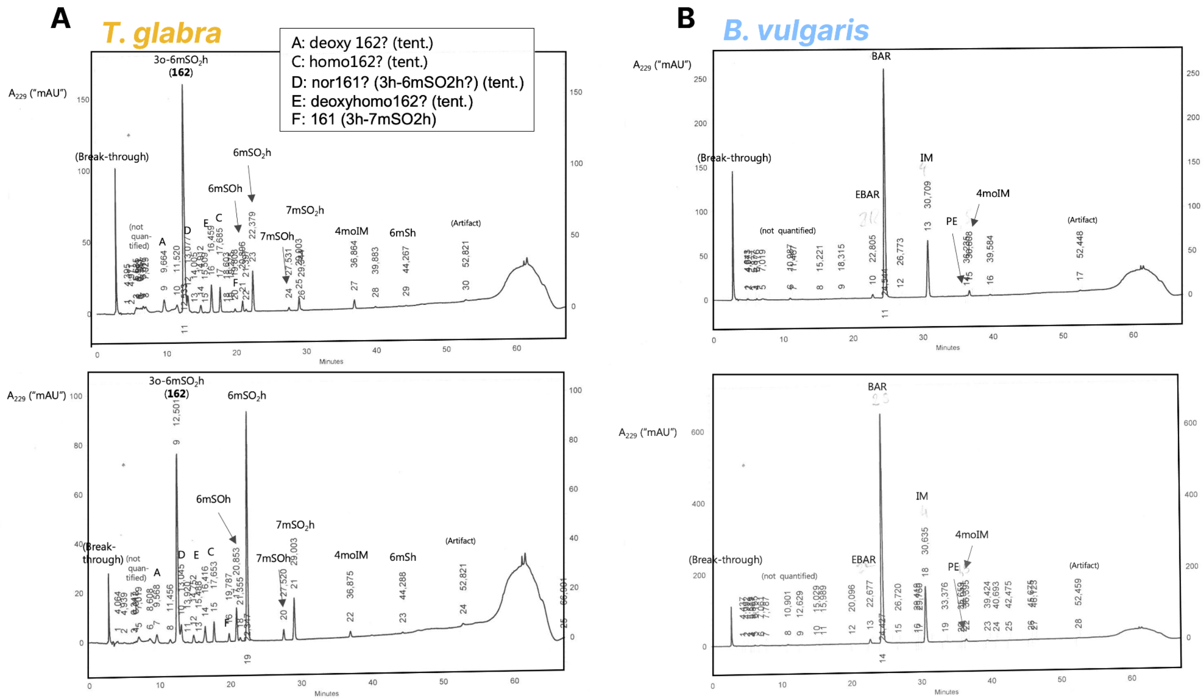

**Figure S2:** HPLC-UV (229 nm) separation, comparison, quantification and preliminary identification of glucosinolates in *T. glabra* (A) and *B. vulgaris* (B); the upper and lower traces within each panel are separate leaf samples (*T. glabra*: 3T, 4T, *B. vulgaris* 2B, 3B). The lower *T. glabra* chromatogram represents an individual with somewhat deviating profile, with 6mSO$_2$h as prominent as the 3-oxo derivative 3o-6mSO$_2$h (**162**). Lettered peaks A, C, D and E in panel A are tentative identifications: A, 3o-6mSOh ("deoxyhomo**162**"); C, 3o-7mSO$_2$h (homo**162**"); D, 3h-6mSO$_2$h ("nor**161**"); E, 3o-7mSOh ("deoxyhomo**162**"); codes in parentheses were used for the same peaks in a separate publication[10]. Peak F was conclusively identified as 3h-7mSO$_2$h (**161**). Abbreviations are defined in Table S4.

#### HPLC-ion trap-MS/MS confirmation of glucosinolates and NMR-identification of glucosinolates in a Danish *T. glabra* accession.

To further investigate the *T. glabra* samples and to confirm the preliminary identification of *B. vulgaris* peaks in HPLC-UV, based on retention time and diode array UV spectrum only, two representative *B. vulgaris* desulfoglucosinolate samples (2B, 6B) and all the *T. glabra* desulfoglucosinolate samples were analyzed in HPLC-ion trap-MS/MS in parallel with authentic references of the *B. vulgaris* peaks and various other references. The results confirmed peak identities of the *B. vulgaris* peaks by retention time, MS and MS2 (based on comparison with authentic references described previously[4]), and confirmed absence of BAR, EBAR and IM in *T. glabra*. (Fig. S3 A and B). However, the major peaks in the *T. glabra* samples were at that time unknown to science. For this reason, a Danish accession available in larger amounts and with a similar profile was investigated and the major, previously unknown glucosinolates identified, as published elsewhere.[10]

The systematic search for additional desulfoglucosinolates reported as "searched for, not found" in Tables S2 and S3 was carried out using HPLC-ion trap-MS/MS data files with the two selected *B. vulgaris* samples 2B and 6B, and with *T. glabra* samples 3T, which was typical for the species, and 4T, which had moderately higher 6m$SO_2$h and lower levels of the otherwise dominant 3o-6m$SO_2$h. Several other *T. glabra* samples including 1T were searched less systematically. These searches were backed by an authentic-standard retention-time and MS/MS library[4,6,11].

**UHPLC-UV-QToF-MS/MS identification of glucosinolates in *T. glabra*.**

In addition to the HPLC-UV and HPLC-ion trap-MS/MS analyses used for quantification and initial comparison of the species, we performed UHPLC-UV-QToF-MS/MS analysis of all eight *T. glabra* samples (Figure S3 C). We diluted samples three-fold with $H_2O$ before this more sensitive analysis and injected only 5 µl sample. The instrument for UHPLC-UV-QToF-MS/MS and its use was as described previously[11]. This analysis was performed because of the much higher mass accuracy and resolution compared to the ion trap experiment, and to allow direct comparison with Danish material used for NMR structure elucidation of equivalent peaks[10]. The *B. vulgaris* samples were not investigated with this instrument since the HPLC-ion trap-MS/MS analyses had already confirmed the identity of all glucosinolates in that species. We compared the samples with an extensive library of standards[6,11] and reference materials, including the Danish *T. glabra* samples and isolated and NMR-verified desulfoglucosinolates described elsewhere[10]. Hence. using the *T. glabra* UHPLC-QToF-UV-MS/MS data to confirm the HPLC-UV peaks allowed a higher-confidence determination of the metabolic differences distinguishing the two host species. The peaks in the *T. glabra* samples matched the peaks identified in

Danish *T. glabra* leaves, allowing us to identify the glucosinolates in U.S. *T. glabra* from this study (Table S3). For five minor *T. glabra* peaks at or below the UV limit of detection, or which could not be meaningfully associated with a specific minor peak in HPLC-UV chromatograms, we measured combined extracted ion chromatography peak intensity of $[M+H]^+$ and $[M+H\text{-anhydroglucose}]^+$ ions at stringent resolution (± 0.005 Da) relative to 6mSOh with the QToF MS detector and used this to estimate levels of 5mSp, 5mSOp, 5mSO$_2$p, 7mSh and 3o-5mSO$_2$p.  (Table S3). Four additional minor *T. glabra* peaks were only tentatively identified in the published structural elucidation[10], as homologs and expected sulfinyls of major peaks. We matched these (3o-6mSOh, 3o-7mSO$_2$h, 3h-6mSO$_2$h, 3o-7mSOh) to the corresponding HPLC-UV peaks in the U.S. material (Table S1; Fig. S2) by comparing relative elution times, diode-array UV spectra (3-oxo derivatives showed a distinct UV band[10]) and elution orders, and these matches were the basis of the reported levels (Table S3).

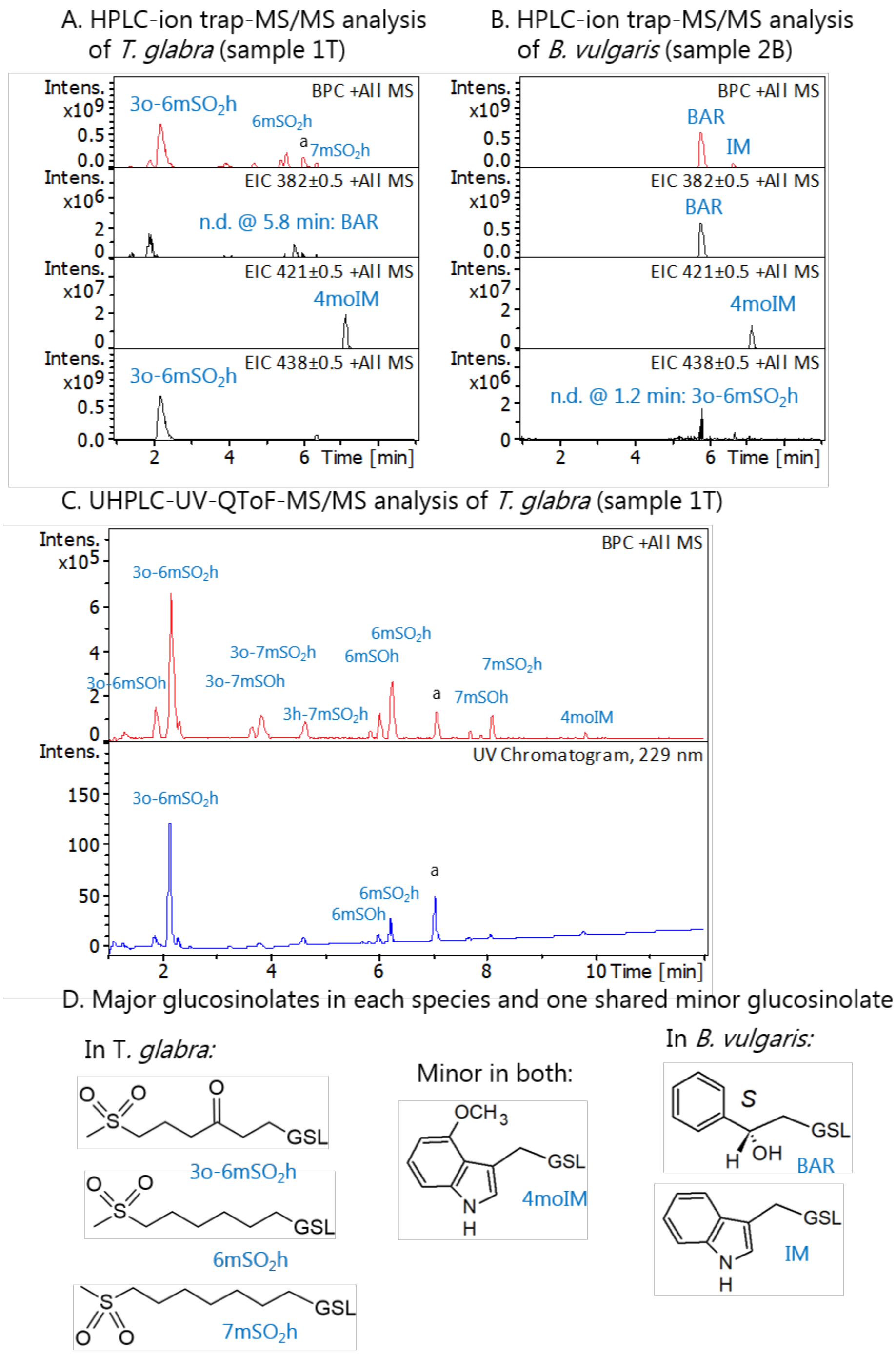

**Figure S3:** Overview of hyphenated liquid chromatography-mass spectrometry analyses for confirmation of glucosinolates in *T. glabra* and *B. vulgaris* leaves (A, B, C) and summary of glucosinolate contrasts and one glucosinolate similarity (D). A+B. Comparative HPLC-ion trap-MS/MS analysis of both species at conditions favoring sodium adducts in MS. The red upper traces depict base peak chromatograms that reveal all major peaks. The three black traces below depict extracted ion chromatograms targeted at $[M+Na]^+$ ions showing presence or absence of BAR, 4moIM and 3o-6mSO$_2$h in each species, confirming peak identities and the between-species contrasts revealed in HPLC-UV. C. Higher resolution UHPLC-UV-QToF-MS-MS analysis of *T. glabra*, used for confirming all major and minor peak identities. The red upper trace reveals the entire diversity of major and minor peaks, but major peaks are above the linear range and appear disproportionately small. All peak identities were backed up by extracted ion chromatograms[10] matched with reference materials at stringent mass conditions (± 0.005 Da) (not shown). The blue lower trace reveals the UV chromatogram (229 nm) detected in line with the mass detector, with all peaks in their linear range and hence with peak areas representative of relative levels in leaves. Peak "a" denotes one or more unidentified compounds whose MS/MS fragmentation is not typical of desulfoglucosinolates. Abbreviations are defined in Table S4.

Only one glucosinolate was common to the two host species (Table S4, Fig. S3). This was 4moIM, at 1.1% of total glucosinolates in *B. vulgaris* and 0.39% in *T. glabra*, and at these levels it is unlikely to be the only important antinutritional compound for a leaf-mining fly. The dominant glucosinolates of the two species differed both in chemical class and, more importantly, in the product they yield on myrosinase-catalyzed hydrolysis. In *B. vulgaris* the main glucosinolate was glucobarbarin (BAR) at 89.7 ± 1.5% of the total (Table S2). Hydrolysis of glucobarbarin is well described[3,8] and yields (*R*)-barbarin, a cyclic oxazolidine-2-thione, rather than an isothiocyanate. Only one very minor *B. vulgaris* glucosinolate, phenethylGSL (PE) at 0.3% of the total, is chemically able to produce a stable isothiocyanate. These plants therefore correspond to the "G-type" *B. vulgaris* chemotype of Western Europe. The glucosinolate profile of *T. glabra* was more complex and was built almost entirely on an aliphatic skeleton derived from chain-elongated methionine (Table S3). Its most abundant glucosinolate was the recently described 6-(methylsulfonyl)-3-oxohexylGSL (3o-6m$SO_2$h) at 59 ± 10% of the total, a compound so far unknown from any other plant species[10]. The second most abundant was its apparent biosynthetic precursor 6-(methylsulfonyl)hexylGSL (6m$SO_2$h), at a median of 7% of the total across samples, with a range of 5–9% in seven of the eight plants. In the remaining plant, sample 4T, 6m$SO_2$h reached 33.7% and nearly matched 3o-6m$SO_2$h at 35.0%, which raised its arithmetic mean to 10.6%. The remaining glucosinolates in *T. glabra* were lower and higher homologs, the expected 3-hydroxy biosynthetic intermediates, and sulfinyl analogs of the major sulfonyls. Mean total glucosinolate levels tended to be somewhat higher in *B. vulgaris* (12.6 ± 8.7 μmol $g^{-1}$ dry weight) than in *T. glabra* (8.9 ± 2.0 μmol $g^{-1}$ dry weight), but the difference was moderate relative to the roughly nine-fold variation among individual *B. vulgaris* plants. Because quantification from UV peak areas relies on approximate and sometimes assumed relative response factors, comparisons between glucosinolates of different chemical classes are approximate whenever their levels are similar, here as elsewhere in the glucosinolate literature.

These differences bear on the selective pressures the two plants impose. Some herbivorous insects neutralize glucosinolates by general mechanisms that act on any glucosinolate, such as hydrolysis of the sulfate ester bond[12], diversion of aglucone fragmentation[13], or glycosylation of the thioglucose residue[14]. Others, including several glucosinolate-adapted Diptera, tolerate the glucosinolate–myrosinase system through efficient detoxification of the isothiocyanate products[2,15], and this is a strategy used by *S. flava*[16]. With the exception of the minor 3-hydroxy derivatives, all of the aliphatic glucosinolates of *T. glabra* are expected to form isothiocyanates upon tissue disruption, so the products of *T. glabra* fall within a chemical class this species is known to encounter. What occurs chemically when *B. vulgaris* is consumed by *S. flava* is not known. The dominant glucosinolate in *T. glabra* yields an oxazolidine-2-thione rather than an isothiocyanate, and whether *S. flava* detoxifies that compound—and by what route—has not been tested. We therefore made no claim that either host is chemically more challenging. Our experiment shows that the two plants differ almost completely in glucosinolate composition and in the chemical class of the products released on tissue damage, so a larva developing inside a leaf of one species encounters a different chemical environment from one developing in the other.

Several caveats are needed when making this comparison. Isothiocyanates have not been confirmed as in vivo hydrolysis products of *T. glabra* by a reliable method. The one early report[17] is not considered dependable and in any case concerns material with a different glucosinolate profile, so the products described here are expected. The biological properties of the *T. glabra* isothiocyanates will also vary with chain length, with sulfinyl versus sulfonyl substitution, and with the hydroxy or oxo group at the 3-position, and because of the reactive oxo group, the dominant glucosinolate may behave unlike the rest. In a direct comparison, moreover, isothiocyanates and oxazolidine-2-thiones exerted approximately equal antinutritional effects on a generalist insect[3], so a difference in the chemical class of the hydrolysis product need not imply a difference in the magnitude of its effect but could well imply a difference in the detoxification enzymes required to cope with defenses in each species. Finally, *B. vulgaris* also contains saponins, typically at lower levels than its glucosinolates. These are detrimental to some but not all insect herbivores[18] and may contribute to the difference between the two hosts.

### Fitness of experimental evolved populations

#### Realized heritability of host-specific relative viability

In this section, we quantified the extent to which selection is able to increase the viability of fly populations on single host plant species, which we interpret as a signature of specialization. Because fly populations, which encounter multiple host plants in the wild, experience different selection pressures on different plant species, we reasoned that selection for any given trait favored on one host environment would compete against selection favoring other traits in a different host environment. Therefore, in order for selection to favor traits on any single host plant species over another host species when populations are exposed to both plant species concurrently, the relative viability benefit to populations evolving on single hosts must be larger than the relative viability benefit of fly populations evolving on multiple hosts. In our experiment, we show that this relative viability increases for both the **B** and **T** single-host environments over the course of our evolution experiment.

Here, we treat this experimentally-derived relative viability as a proxy for an unknown set of traits underlying specialization on a single host plant species at the population level. We sought to quantify the extent to which natural selection can act on this proxy to increase relative viability in either of the experimental host environments we used in this study, **B** and **T**. To do this, we employed a realized heritability approach via the breeder's equation.

First, we defined relative population viability as the ratio of survivors from each single-host environment to the geometric mean of the **B+T** environment in the equivalent generation. Viability counts were measured at generations 2, 3, and 10. Early generations (2+3) were grouped as the pre-selection population

($G_0$) and the 10th generation was designated as the post-selection population ($G_{10}$). To normalize across replicates and reduce the influence of extreme values, raw counts were log-transformed and expressed as log-ratios relative to the geometric mean of **B+T**:

$$\ln(\text{Ratio}) = \ln\left(\frac{v_{B|T}}{\underline{v}_{BT}}\right)$$

where $v_{B|T}$ is the viability count of the single-host environment (**B** or **T**) and $\underline{v}_{BT}$ is the geometric mean viability of the **B+T** environment. This log-ratio measures host-specific viability relative to the multiple host reference and was used as our operational phenotype for specialization.

Log-ratios were averaged across replicates within each generation group and environment to obtain the mean log-ratio for each single-host population at $G_0$ and $G_{10}$. We calculated the selection differential (S) for each single-host population as the deviation of its mean log-ratio at $G_0$ from the mean log-ratio across all populations at $G_0$,

$$S = \ln\underline{(\text{ratio}})_{B|T,G0} - \ln\underline{(\text{ratio}})_{B+T,G0}$$

The implied response to selection (R) was defined as the change in the mean log-ratio between $G_0$ and $G_{10}$ for each environment:

$$R = \ln\underline{(\text{ratio}})_{B|T,G10} - \ln\underline{(\text{ratio}})_{B|T,G0}$$

Finally, we estimated a realized heritability-like response coefficient ($h^2$) for relative viability using a modified breeder's equation, adjusted for the number of generations (G) elapsed between $G_0$ and $G_{10}$,

$$h^2 = \frac{R}{S \times G}$$

Estimates were calculated for $G = 7$ and $G = 8$ to provide a range of heritability values representing the interval between $G_0$ and $G_{10}$. We emphasize that this estimate does not represent a direct partitioning of phenotypic variance into additive genetic and environmental components. Rather, it summarizes how strongly the population-level relative viability phenotype changed in the direction predicted by the initial selection differential.

In the **T** environment, realized heritability estimates were high and positive, with $h^2$ ranging from 0.607 ($G = 8$) to 0.693 ($G = 7$). This indicates that up to 69% of the variance in relative viability (our proxy for traits underlying specialization in multiple-host environemnts) was explained by additive genetic effects, with the population responding to selection in the same direction as initially favored. For **B** environment, $h^2$ estimates ranged from -0.424 ($G = 8$) to -0.485 ($G = 7$), indicating up to 48%. This negative heritability reflects a scenario where the response to selection opposed the direction of the selection differential.

#### Test of reciprocal fitness trade-offs in post-evolution experimental flies

To test for evolved performance trade-offs, 10th-generation F1 adults from six experimental replicates (two from each of the three original treatments) were first reared for one generation on each host plant species (**B**, **T**, or a 1:1 mixture of both **B+T**) to minimize maternal environmental effects. From these rearing cages, 60 flies (1:1 male:female) were collected and introduced into phenotyping cages containing six plants: either six *B. vulgaris*, six *T. glabra*, or three of each species. Flies mated for five days, after which the plants were removed. Fresh plants were then provided for 24 hours of oviposition, after which each plant was separately bagged for larval development. Each bagged plant was placed in a 36-cell tray - with one plant for each replicate and test treatment combination - to minimize cage effects on larval development. The number of emerging flies was recorded daily for 10 days. We found a delay in development over this period when flies were reared on *T. glabra* compared to *B. vulgaris* and the 1:1 mixture of both plants (Figure S6), suggesting *T. glabra* is a more challenging host environment for *S. flava* than *B. vulgaris*.

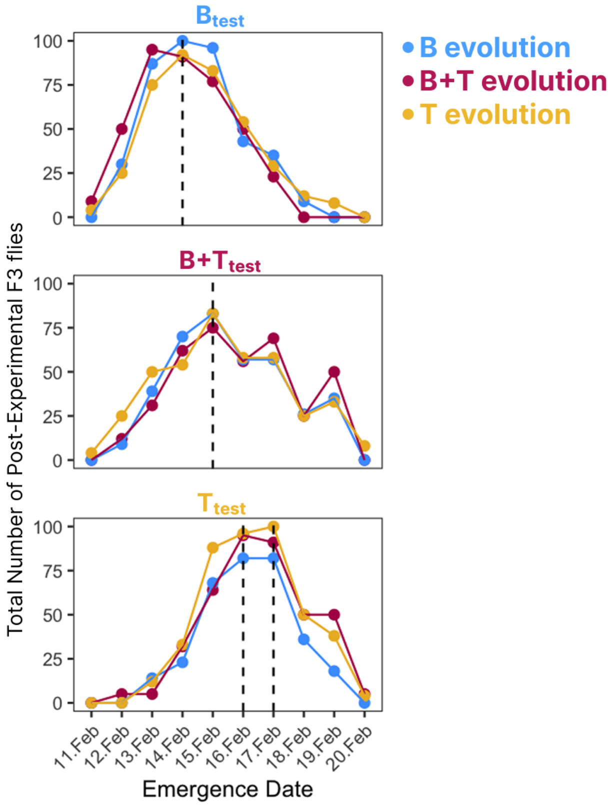

**Figure S6:** Developmental timing is determined primarily by the host plant environment rather than evolutionary history of the *S. flava* population. Total daily counts of emerging post-experimental F3 adults are shown for populations from the **B** evolution (blue), **B+T** evolution (magenta), and **T** evolution (yellow) environments. Panels indicate the specific test environment used for the performance assay ($B_{test}$, top; $B+T_{test}$, middle; $T_{test}$, bottom). Dashed vertical lines mark the date(s) of peak emergence. Consistent with *T. glabra* representing a more challenging host, development was delayed in the $T_{test}$ environment. Flies tested on *T. glabra* (bottom) peaked approximately 2-3 days later (Feb 16-17) than those tested on *B. vulgaris* (top; Feb 14), regardless of their evolutionary background. Flies tested on the multiple-host environment ($B+T_{test}$) exhibited intermediate emergence timing (Feb 15).

We normalized the total number of flies emerging over the 10-day period by the number of females originally used (n = 30) and the number of plants available to them for the 24-hour oviposition period (n = 6). These normalized counts were modeled using a negative binomial GLM approach:

$$y_i \sim \text{NegativeBinomial}(\lambda_i, \theta)$$
$$\log(\lambda_i) = \beta_0 + \beta_1 X_{i1} + \beta_2 X_{i2} + \beta_3 (X_{i3}) + \log(E_i)$$

where $X_{i1}$ is the evolution treatment, $X_{i2}$ is the test environment, $X_{i3}$ denotes whether the evolution environment and the test environment match, and $E_i$ is an offset for the number of host plants available to ovipositing females. The $\beta_3$ term captured interaction effects between evolution environment and the test environment (i.e., reciprocal fitness trade-off).

The model was fitted using the R function glm.nb from the MASS package. We assessed statistical significance for specific evolutionary hypotheses using linear contrasts derived from the model coefficients (via multicomp and emmeans). The first hypothesis we tested was the "Home-Court Advantage Hypothesis" (p = 0.0257) in which we tested the prediction that populations would perform significantly better in the test environment matching their evolution environment ("home") compared to an alternative test environment ("away"). This was calculated as a contrast comparing the average fitness of "matched" cases (e.g., **B** on $B_{test}$) versus mismatched cases. The second hypothesis was the "Specialist Trade-off Hypothesis" (p = 0.004) in which we tested the prediction that populations evolving specialization (**B** and **T**) would exhibit reciprocal fitness trade-offs between test environments. The third hypothesis was the "Generalist Jack-Of-All-Trades Hypothesis" (one-sided p = 0.05) in which we assessed the performance reaction norm of the **B+T** (generalist) populations relative to the specialists across all test environments using a one-sided interaction contrast. Finally, the fourth hypothesis was the "Generalist Innovation Hypothesis" (*p* = 0.93) in which we tested for the benefit of generalization compared to specialization in the **B+T** test environment.

Table S5. Reciprocal performance assay results, modeled (Negative Binomial GLM) versus empirical adult-production estimates per (evolution × test) environment combination. Values are means (95% confidence intervals in parentheses). Modeled values are predictions from the per-host genotypic-class fit; empirical values are raw F3 adult counts per ovipositing F2 female per seedling, averaged across cage replicates. Home-court combinations, where the evolution and test environments match, lie on the diagonal. The four single-host combinations (**B**-evolved × $B_{test}$, **B**-evolved × $T_{test}$, **T**-evolved × $B_{test}$, **T**-evolved × $T_{test}$) form the 2 × 2 single-host specialist trade-off matrix used to test the Specialist Trade-off Hypothesis.

| | | **Test environment** | | |
|---|---|---|---|---|
| **Evolution environment** | **Estimate** | **$B_{test}$** | **$[B+T]_{test}$** | **$T_{test}$** |
| **B** | modeled | 2.89 (2.21, 3.79) | 2.64 (2.01, 3.46) | 1.77 (1.32, 2.36) |
| | empirical | 2.89 (2.26, 3.52) | 2.65 (1.83, 3.47) | 1.78 (1.09, 2.48) |
| **B+T** | modeled | 2.67 (2.02, 3.52) | 3.11 (2.24, 4.32) | 2.33 (1.76, 3.09) |
| | empirical | 2.67 (1.98, 3.37) | 3.13 (1.90, 4.36) | 2.34 (1.53, 3.14) |
| **T** | modeled | 1.94 (1.48, 2.55) | 2.31 (1.76, 3.02) | 2.61 (2.00, 3.40) |
| | empirical | 1.94 (1.28, 2.60) | 2.31 (1.65, 2.96) | 2.61 (2.04, 3.18) |

#### Resolution of the two fitness measurements

The main text reports two measurements of fitness that use different data. Population viability (Fig. 2A) is the total number of adults emerging from a whole cage, and therefore integrates the output of the 500 parents seeded into that cage together with density-dependent survival of their larvae. The reciprocal performance assay (Fig. 2B) was the number of adults produced per plant by 30 F2 females at controlled density. We ran this analysis on populations 3 and 4 only. We found that single-host populations gained viability advantages relative to two-host populations in the experimental evolution cages, whereas the assay found no difference between them in their home environments ($p = 0.93$). We therefore asked whether these two results are statistically incompatible.

We placed both on a common scale as the ratio of specialist to two-host production in the home environment. For the assay we pooled the **B**-on-**B** and **T**-on-**T** plants ($n = 47$) and compared them with the **B+T**-on-**B+T** plants ($n = 17$), bootstrapping the ratio of means over 20,000 resamples of plants within cells. For the cages we took relative viability at $t_{10}$ on the scale of Fig. 2A. We then estimated the power of the assay by resampling the observed per-plant counts with a multiplicative home advantage applied to the specialist plants, testing each simulated dataset with a two-sided Welch $t$-test on log-transformed counts (4,000 simulations per effect size).

Per-plant production was highly variable and the coefficient of variation was 0.52 in both single-host cells and 0.71 in the two-host cell, which was also the smallest and the most dispersed ($n = 17$, range 2–36 adults per plant). Both specialist means fall inside the two-host distribution, at its 65th (**B** on **B**) and 59th (**T** on **T**) percentiles (Fig. S4A). The assay estimates a home advantage of 1.09×, with a 95% bootstrap confidence interval of 0.79–1.63×. The population cages place the specialists at 1.25× the two-host cages at $t_{10}$ (**B** 1.19×, **T** 1.32×), which falls inside that interval. The assay had 0.57 power to detect an effect of 1.25×, reaching 0.80 only near 1.4× (Fig. S4B); resolving a 1.25× difference at 80% power would have required roughly 30 plants per cell compared to the 17–24 available. The two measurements are therefore compatible. The reciprocal assay is adequately powered as a test of host-specific antagonism, where the reciprocal difference between the two single-host treatments is 2.2-fold, but it is not a sensitive test of the population-level response.

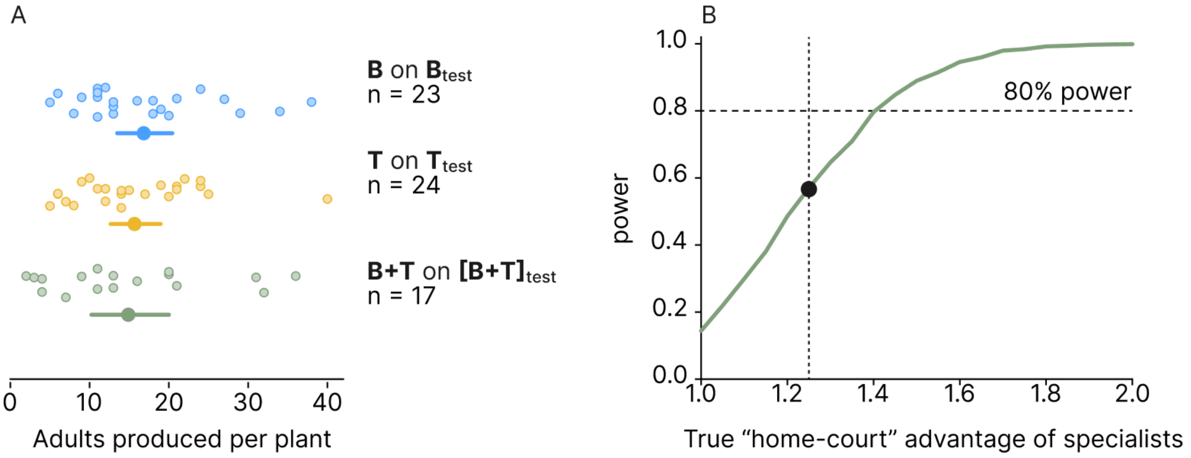

**Figure S4.** (**A**) Adults produced per plant in the three “home-court” environments; small points are individual plants, the larger point is the cell mean and the bar its 95% bootstrap confidence interval. Both specialist means fall within the two-host distribution. (**B**) Power of the assay to detect a true “home-court” advantage of specialists compared to flies evolved on two hosts, from resampling the observed per-plant counts. The dotted line marks the 1.25× advantage measured in the population cages, which the assay resolves with 0.57 power; the dashed line marks 80% power.

### Equilibrium dynamics from fitness differentials

#### <u>The Levene Model</u>

To analyze the conditions required for genetic equilibrium, we employed the theoretical model originally proposed by Levene (1953). This model assumes a single panmictic population where zygotes disperse randomly into multiple ecological niches. Selection occurs through differential mortality within each niche before the survivors emerge to mate randomly in a global pool.

Levene demonstrated that the change in allele frequency ($\Delta q$) per generation in a two-allele system across multiple niches is given by the weighted mean of the changes within each niche. For a two-niche system (Environment **B** and Environment **T**), the equilibrium condition ($\Delta q = 0$) is satisfied when:

$$\Delta q = \frac{c s_B p q}{1 - 2 s_B q} - \frac{(1-c) s_T p q}{1 - 2 s_T (1-q)} = 0$$

Where:

c is the proportion of the total survivors contributed by Niche **B** (effective niche size).

$s_B$ and $s_T$ are the selection coefficients in Niche **B** and Niche **T**, respectively.

#### Parameterization of the Levene Model

To visualize the theoretical boundaries of stability (the "heatmap" background in Fig. S5), we assumed an additive fitness model where the heterozygote phenotype is intermediate. We parameterized fitness using selection coefficients $s_B$ (favoring genotype AA in Niche B) and $s_T$ (favoring genotype aa in Niche T).

| Genotype | Phenotype | Fitness in Niche B (Levene's W) | Fitness in Niche T (Levene's V) |
|---|---|---|---|
| AA[**B**] | $w[B_{test}]$ | $W_{AA} = (1 + 2s_B)/(1 + s_B)$ | $V_{AA} = 1/(1 + s_T)$ |
| Aa[**B+T**] | $w[BT_{test}]$ | $W_{Aa} = 1$ | $V_{Aa} = 1$ |
| aa[**T**] | $w[T_{test}]$ | $W_{aa} = 1/(1 + s_B)$ | $V_{aa} = (1 + 2s_T)/(1 + s_T)$ |

We numerically solved Levene's equation for equilibrium ($\hat{q}$) across a grid of availability (c) and selection asymmetries ($k = s_T/s_B$). The regions presented as a mixture of yellow (B) and blue (T) in the figures represent the parameter space where a protected polymorphism is possible ($0 < \hat{q} < 1$).

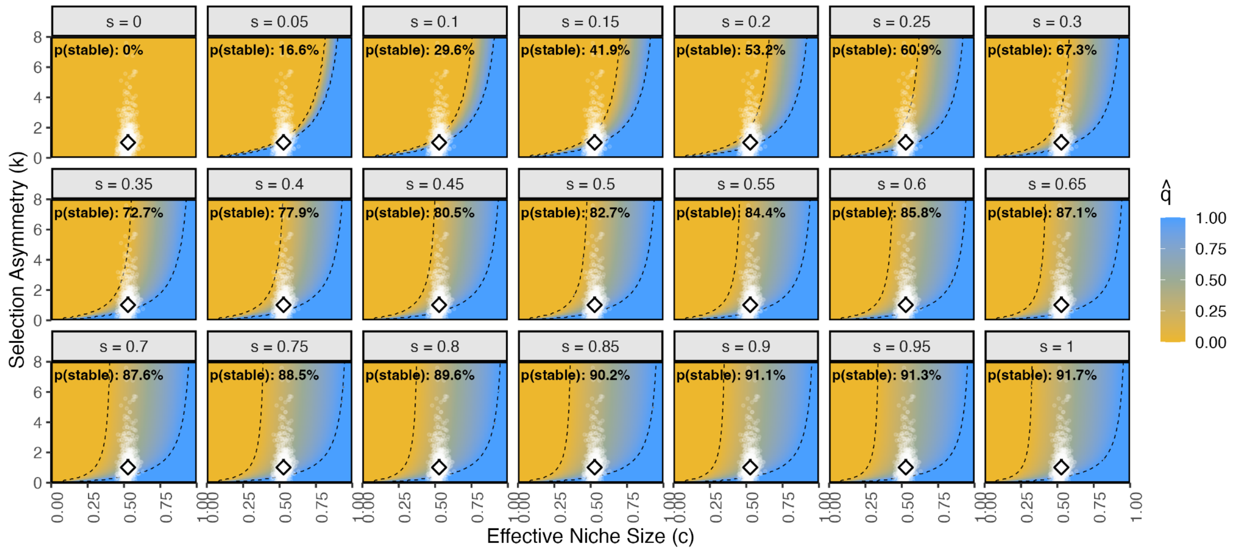

**Figure S5.** Sensitivity analysis of Levene's stability conditions to varying selection strengths. The theoretical parameter space for stable polymorphism (grey regions) is projected across increasing values of the selection coefficient (s), ranging from weak ($s = 0.05$) to strong ($s = 1.0$) selection. In each panel, the background heatmap represents the theoretical equilibrium frequency of the allele ($\hat{q}$) under Levene's model assuming additive fitness. Regions bounded by dashed lines indicate conditions where protected polymorphism is possible ($0 < \hat{q} < 1$).The overlaid cloud (white points) represents 1,000 non-parametric bootstrap estimates of the effective niche size (c) and selection asymmetry (k) derived from the empirical population data. The distribution of empirical estimates is identical in every panel. The central diamond and error bars represent the median and interquartile range of the empirical estimates.The annotation in the upper left ("p(stable)") indicates the probability that the populations satisfy the stability criteria for that specific theoretical selection strength (calculated as the proportion of bootstrap iterations falling within the stable region).

#### Empirical estimation via non-parametric bootstrap

We next tested whether the experimentally observed fitness trade-offs were sufficient to maintain polymorphism within this theoretical landscape. To do this, we used a non-parametric bootstrap approach and performed 1,000 bootstrap iterations. In each iteration, the original dataset was resampled with replacement and modeled using a negative binomial GLM as above. Absolute fitness values (w) were predicted for every genotype (AA, Aa, aa) in both test environments ($B_{test}$, $T_{test}$).

For each iteration, we calculated the effective niche size (c) as the proportion of total population productivity contributed by the evolution environment B:

$$c = \frac{\sum \widehat{w}_B}{\sum \widehat{w}_B + \sum \widehat{w}_T}$$

We calculated the selection asymmetry (k) using the ratio of selection coefficients. Selection strength (s) was calculated as the relative fitness advantage of the favored homozygote over the disfavored homozygote in that specific niche:

$$s_B = \frac{w_{B,AA} - w_{B,aa}}{w_{B,AA}}, \quad s_T = \frac{w_{T,aa} - w_{T,AA}}{w_{T,aa}}$$

$$k = \frac{s_T}{s_B}$$

**<u>Test of stability (invasion analysis)</u>**

To determine equilibrium stability, we assessed the ability of a rare allele to invade a population fixed for the common allele. Levene (1953) proved that a stable equilibrium exists if the weighted harmonic mean of the resident homozygote fitness (relative to the heterozygote) is less than one. This is equivalent to the condition that the rare heterozygote has a weighted fitness advantage > 1.

We calculated two Invasion Indices (I) for each bootstrap iteration, where the invasion of allele A (rare A into common aa) was:

$$I_A = \left[c\left(\frac{w_{B,Aa}}{w_{B,aa}}\right)\right] + \left[(1-c)\left(\frac{w_{T,Aa}}{w_{T,aa}}\right)\right] > 1$$

The invasion of a (rare a into common AA) was:

$$I_a = \left[c\left(\frac{w_{B,Aa}}{w_{B,AA}}\right)\right] + \left[(1-c)\left(\frac{w_{T,Aa}}{w_{T,AA}}\right)\right] > 1$$

The system is in a state of protected polymorphism only if both conditions ($I_A > 1$ and $I_a > 1$) are met simultaneously.

The probability of stability ($P_{Stable}$) was calculated as the proportion of bootstrap iterations in which both invasion criteria were satisfied. For visualization, the empirical results are presented as the median value of the bootstrap distribution, with error bars representing the interquartile range (25th-75th percentiles) for variable c and k.

#### *Drosophila* Muller-element assignment for the *S. flava* v2 assembly

The *S. flava* v2 assembly included 12,254 protein-coding genes distributed across 162 gene-bearing scaffolds. We assigned scaffolds to *D. melanogaster* Muller elements to enable downstream comparison between autosomal and sex-chromosome variants. We extracted the *S. flava* v2 proteome by parsing the v2 GFF for all coding-sequence (CDS) records. Sequences shorter than 30 amino acids or with more than 5% internal stop codons were discarded. Of 12,365 input gene records, 12,254 (99.1%) produced a usable protein sequence for downstream analysis. We aligned the resulting proteome against the *D. melanogaster* reference proteome (UniProt UP000000803, 13,826 sequences) using diamond blastp v2.1.10 in --more-sensitive mode, retaining the top alignment per query at e-value ≤ $10^{-5}$. 11,271 of 12,254 *S. flava* queries (92%) produced a hit. This allowed us to construct a *D. melanogaster* chromosome lookup table that we used to map *S. flava* genes to *D. melanogaster* chromosomes. We converted chromosomes to Muller elements using the standard correspondence (X→A, 2L→B, 2R→C, 3L→D, 3R→E, 4→F). For each *S.*

*flava* scaffold we counted the Muller assignments of its proteins and classified the scaffold as confident (≥ 10 Muller-resolved genes and ≥ 70% top-Muller agreement), ambiguous (≥ 5 genes and ≥ 50%), or unclassified otherwise. Of 162 scaffolds with at least one annotated gene, 5 were classified as confident and 8 as ambiguous. The remaining 149 scaffolds were short (mostly under 50 kb) with too few genes for a reliable determination of the muller element. The five confident assignments span four of the six Muller elements (A, B, E, F), with the largest scaffold in the assembly (scaffold 597) unclassified. Inspection of scaffold 597 revealed a previously unknown chromosomal fusion. Its 4,679 annotated genes (3,281 of which resolved to a *D. melanogaster* chromosome) showed a bimodal Muller distribution where 1,616 genes (49%) mapping to Muller C and 1,530 (47%) to Muller D, with all other elements below 2%. The scaffold 597 pattern is consistent with a 2R + 3L fusion. Muller-element fusions are recurring in Drosophilidae karyotype evolution, and *Scaptomyza* + Hawaiian *Drosophila* species (which form a monophyletic clade) carry many lineage-specific Muller-element rearrangements [19,20].

**Table S10.** Per-scaffold majority-vote Muller-element assignments for the four chromosome-arm-scale candidate scaffolds in the antagonism analyses. Top-Muller agreement is the percentage of Muller-resolved genes on each scaffold whose *D. melanogaster* orthologs map to the majority Muller element. Confident classification requires ≥ 10 resolved genes and ≥ 70% top-Muller agreement. Scaffold 597 fails the threshold because it included a 2R + 3L (Muller C + D) fusion discovered here.

| **Scaffold** | **Top Muller (chrom)** | **Resolved genes (n)** | **Top-Muller agreement** | **Classification** |
|---|---|---|---|---|
| chr_ScDA7r2_439_HRSCAF_779 | A (X) | 1,295 | 93% | Confident |
| chr_ScDA7r2_126_HRSCAF_325 | B (2L) | 1,578 | 92% | Confident |
| chr_ScDA7r2_110_HRSCAF_295 | E (3R) | 2,070 | 93% | Confident |
| chr_ScDA7r2_2_HRSCAF_23 | F (4) | 78 | 92% | Confident |
| chr_ScDA7r2_597_HRSCAF_953 | C (2R) + D (3L) | 3,281 | 49% / 47% | Unclassified (fusion) |

### Population genomic sequencing

#### Pooled sequencing of experimental and wild populations

Genomic DNA was extracted and sequenced from both experimental and wild populations of *S. flava* in multiple sequencing efforts over a ten year span, from 2015-2025.

In 2015 at the University of Arizona, DNA was extracted from wild and experimental pools of 80 female flies using the Kapa HyperPrep Kit (KK8503). Founder samples consisted of two pools of 80 females, evolved populations were represented by one pool of 80 females per replicate at generation 10, and wild populations were sampled from three fallow field sites in New Hampshire, with 22-60 flies per host plant. Libraries were sequenced on an Illumina HiSeq 2500 platform at the University of Arizona across three lanes, with ½ lane allocated per library. Sequencing was carried out with 100 bp paired-end reads and the actual distribution averaged ~180 bp, creating a 20 bp overlap between read pairs. Reads were collapsed into ~180 bp single-end equivalents for downstream mapping.

In 2016, sequencing was repeated at the University of California, Berkeley for the experimental pools. All 12 evolved populations at generation 10, the six wild populations, and pooled founder samples were prepared as indexed libraries and combined into a multiplexed "superpool". Seventy-five percent of the sequencing effort was devoted equally to the 12 evolved populations, and 25% to the six wild populations. Sequencing was performed on an Illumina HiSeq 4000 across four lanes with 100 bp paired-end reads and an average insert size of 350-400 bp.

In 2022, an expanded sampling of 114 DNA pools representing all experimental and wild populations was conducted at the University of California, Berkeley and New York University. DNA was extracted using Qiagen DNeasy Blood and Tissue Kits, with 40 flies processed per batch and extracts pooled 1:1 to generate final samples representing 80 females. DNA concentration was measured with a Qubit High Sensitivity dsDNA assay and purity confirmed using a NanoDrop spectrophotometer. Libraries were generated using SeqWell plexWell™ LP 384, a multiplexed workflow, with three independent libraries prepared per pool to maximize coverage. Sequencing was performed on Illumina NovaSeq with 250 bp paired-end reads, producing high-coverage datasets suitable for allele frequency estimation and haplotype analysis.

Table S8. Per-pool sequencing and mapping quality summary for the *S. flava* experimental evolution time series and wild samples. The founder pools(F1G00) represent the four 80-fly subsamples (F1-F4) of the expanded outbred founder colony. The "pools" column gives the count of pools and the number of sequencing batches per pool. Serial-evolution pools are labeled by treatment cage (B1-B4 = *B. vulgaris*, M1-M4 = **B+T** environment, T1-T4 = *T. glabra*) and generation (G01-G10). Generations G03-G05 for cages B3, B4, M3, M4, T3, T4 are absent from the time series. Mean coverage values reflect post-deduplication depth. Wild pools (AVB, AVT, PSB, PST, RMB, RMT) are 22-60 fly pools collected at three New Hampshire localities (AV, PS, RM near Newington) in summer 2014, sequenced across two batches (2015 UA HiSeq 2500, 2016 UCB HiSeq 4000). QC values were derived from the post-realignment BAM files. Duplication rate is left blank for wild pools because duplicates were removed before realignment. Insert size for wild pools is reported as the mid-point of the 350-400 bp range.

| **Sample** | **Origin** | **Gen** | **Host** | **Pools (batches/pool)** | **MapQ** | **Dup (%)** | **Insert (bp)** | **Coverage** | **Mapped (%)** |
|---|---|---|---|---|---|---|---|---|---|
| AVB | Wild | | *Barbarea vulgaris* | 1 (3) | 54 | <5 | ~375 | 40x | 100 |
| AVT | Wild | | *Turritis glabra* | 1 (3) | 54 | <5 | ~375 | 39x | 100 |
| PSB | Wild | | *Barbarea vulgaris* | 1 (3) | 53 | <5 | ~375 | 42x | 100 |
| PST | Wild | | *Turritis glabra* | 1 (3) | 54 | <5 | ~375 | 36x | 100 |
| RMB | Wild | | *Barbarea vulgaris* | 1 (3) | 55 | <5 | ~375 | 32x | 100 |
| RMT | Wild | | *Turritis glabra* | 1 (3) | 53 | <5 | ~375 | 43x | 100 |
| F1G00 | Founder | 0 | Mixture | 4 (8,8,8,8) | | | | | |

| **Sample** | **Origin** | **Gen** | **Host** | **Pools (batches/pool)** | **Ma pQ** | **Dup (%)** | **Insert (bp)** | **Coverage** | **Mapped (%)** |
|---|---|---|---|---|---|---|---|---|---|
| B1G01 | Serial | 1 | *Barbarea vulgaris* | 1 (3) | 47 | 33 | 234 | 39x | 100 |
| B1G02 | Serial | 2 | *Barbarea vulgaris* | 1 (3) | 47 | 33 | 240 | 42x | 100 |
| B1G03 | Serial | 3 | *Barbarea vulgaris* | 1 (4) | 46 | 31 | 293 | 36x | 100 |
| B1G04 | Serial | 4 | *Barbarea vulgaris* | 1 (4) | 48 | 33 | 300 | 41x | 100 |
| B1G05 | Serial | 5 | *Barbarea vulgaris* | 1 (3) | 47 | 37 | 300 | 50x | 100 |
| B1G06 | Serial | 6 | *Barbarea vulgaris* | 1 (3) | 47 | 36 | 230 | 47x | 100 |
| B1G07 | Serial | 7 | *Barbarea vulgaris* | 1 (3) | 47 | 34 | 241 | 42x | 100 |
| B1G08 | Serial | 8 | *Barbarea vulgaris* | 1 (3) | 47 | 33 | 234 | 41x | 100 |
| B1G09 | Serial | 9 | *Barbarea vulgaris* | 1 (4) | 46 | 31 | 293 | 38x | 100 |
| B1G10 | Serial | 10 | *Barbarea vulgaris* | 1 (4) | 47 | 27 | 285 | 28x | 100 |
| B2G01 | Serial | 1 | *Barbarea vulgaris* | 1 (3) | 47 | 33 | 231 | 40x | 100 |
| B2G02 | Serial | 2 | *Barbarea vulgaris* | 1 (3) | 48 | 37 | 232 | 50x | 100 |
| B2G03 | Serial | 3 | *Barbarea vulgaris* | 1 (3) | 47 | 30 | 235 | 34x | 100 |
| B2G04 | Serial | 4 | *Barbarea vulgaris* | 1 (3) | 47 | 29 | 234 | 32x | 100 |
| B2G05 | Serial | 5 | *Barbarea vulgaris* | 1 (4) | 46 | 37 | 303 | 53x | 100 |
| B2G06 | Serial | 6 | *Barbarea vulgaris* | 1 (3) | 47 | 32 | 234 | 38x | 100 |
| B2G07 | Serial | 7 | *Barbarea vulgaris* | 1 (3) | 47 | 28 | 247 | 29x | 100 |
| B2G08 | Serial | 8 | *Barbarea vulgaris* | 1 (3) | 48 | 29 | 249 | 32x | 100 |
| B2G09 | Serial | 9 | *Barbarea vulgaris* | 1 (3) | 48 | 29 | 244 | 30x | 100 |
| B2G10 | Serial | 10 | *Barbarea vulgaris* | 2 (4,4) | 47 | 40 | 295 | 64x | 100 |
| B3G01 | Serial | 1 | *Barbarea vulgaris* | 1 (3) | 48 | 33 | 246 | 43x | 100 |

| Sample | Origin | Gen | Host | Pools (batches/pool) | Ma pQ | Dup (%) | Insert (bp) | Coverage | Mapped (%) |
|---|---|---|---|---|---|---|---|---|---|
| B3G02 | Serial | 2 | *Barbarea vulgaris* | 1 (3) | 47 | 31 | 242 | 37x | 100 |
| B3G06 | Serial | 6 | *Barbarea vulgaris* | 1 (3) | 47 | 30 | 243 | 35x | 100 |
| B3G07 | Serial | 7 | *Barbarea vulgaris* | 1 (3) | 47 | 36 | 232 | 47x | 100 |
| B3G08 | Serial | 8 | *Barbarea vulgaris* | 1 (3) | 47 | 29 | 248 | 33x | 100 |
| B3G09 | Serial | 9 | *Barbarea vulgaris* | 1 (3) | 47 | 36 | 225 | 45x | 100 |
| B3G10 | Serial | 10 | *Barbarea vulgaris* | 2 (3,3) | 48 | 44 | 235 | 76x | 100 |
| B4G01 | Serial | 1 | *Barbarea vulgaris* | 1 (3) | 47 | 37 | 246 | 51x | 100 |
| B4G02 | Serial | 2 | *Barbarea vulgaris* | 1 (3) | 48 | 35 | 234 | 46x | 100 |
| B4G06 | Serial | 6 | *Barbarea vulgaris* | 1 (3) | 48 | 30 | 226 | 33x | 100 |
| B4G07 | Serial | 7 | *Barbarea vulgaris* | 1 (4) | 46 | 35 | 301 | 47x | 100 |
| B4G08 | Serial | 8 | *Barbarea vulgaris* | 1 (3) | 47 | 28 | 247 | 31x | 100 |
| B4G09 | Serial | 9 | *Barbarea vulgaris* | 1 (3) | 48 | 28 | 239 | 30x | 100 |
| B4G10 | Serial | 10 | *Barbarea vulgaris* | 2 (3,3) | 48 | 40 | 250 | 61x | 100 |
| M1G01 | Serial | 1 | Subdivided | 1 (3) | 48 | 30 | 234 | 35x | 100 |
| M1G02 | Serial | 2 | Subdivided | 1 (3) | 47 | 40 | 233 | 60x | 100 |
| M1G03 | Serial | 3 | Subdivided | 1 (3) | 47 | 30 | 239 | 35x | 100 |
| M1G04 | Serial | 4 | Subdivided | 1 (3) | 47 | 31 | 232 | 34x | 100 |
| M1G05 | Serial | 5 | Subdivided | 1 (4) | 48 | 37 | 294 | 50x | 100 |
| M1G06 | Serial | 6 | Subdivided | 1 (3) | 47 | 28 | 235 | 29x | 100 |
| M1G07 | Serial | 7 | Subdivided | 1 (3) | 47 | 32 | 242 | 40x | 100 |
| M1G08 | Serial | 8 | Subdivided | 1 (3) | 47 | 30 | 245 | 34x | 100 |
| M1G09 | Serial | 9 | Subdivided | 1 (3) | 48 | 31 | 249 | 36x | 100 |
| M1G10 | Serial | 10 | Subdivided | 2 (3,3) | 47 | 41 | 235 | 65x | 100 |
| M2G01 | Serial | 1 | Subdivided | 1 (3) | 47 | 33 | 251 | 41x | 100 |
| M2G02 | Serial | 2 | Subdivided | 1 (3) | 47 | 38 | 254 | 53x | 100 |
| M2G03 | Serial | 3 | Subdivided | 1 (4) | 47 | 34 | 298 | 44x | 100 |

| Sample | Origin | Gen | Host | Pools (batches/pool) | Ma pQ | Dup (%) | Insert (bp) | Coverage | Mapped (%) |
|---|---|---|---|---|---|---|---|---|---|
| M2G04 | Serial | 4 | Subdivided | 1 (3) | 47 | 28 | 252 | 31x | 100 |
| M2G05 | Serial | 5 | Subdivided | 1 (3) | 48 | 35 | 248 | 46x | 100 |
| M2G06 | Serial | 6 | Subdivided | 1 (3) | 47 | 31 | 250 | 36x | 100 |
| M2G07 | Serial | 7 | Subdivided | 1 (3) | 48 | 25 | 252 | 24x | 100 |
| M2G08 | Serial | 8 | Subdivided | 1 (4) | 46 | 29 | 303 | 33x | 100 |
| M2G09 | Serial | 9 | Subdivided | 1 (4) | 46 | 32 | 299 | 40x | 100 |
| M2G10 | Serial | 10 | Subdivided | 1 (3) | 47 | 27 | 240 | 28x | 100 |
| M3G01 | Serial | 1 | Subdivided | 1 (3) | 47 | 35 | 245 | 46x | 100 |
| M3G02 | Serial | 2 | Subdivided | 1 (3) | 47 | 36 | 234 | 46x | 100 |
| M3G06 | Serial | 6 | Subdivided | 1 (3) | 47 | 33 | 229 | 40x | 100 |
| M3G07 | Serial | 7 | Subdivided | 1 (3) | 48 | 29 | 246 | 32x | 100 |
| M3G08 | Serial | 8 | Subdivided | 1 (3) | 48 | 30 | 249 | 34x | 100 |
| M3G09 | Serial | 9 | Subdivided | 1 (4) | 46 | 28 | 284 | 31x | 100 |
| M3G10 | Serial | 10 | Subdivided | 1 (4) | 47 | 26 | 286 | 27x | 100 |
| M4G01 | Serial | 1 | Subdivided | 1 (3) | 48 | 33 | 242 | 42x | 100 |
| M4G02 | Serial | 2 | Subdivided | 1 (4) | 46 | 33 | 296 | 42x | 100 |
| M4G06 | Serial | 6 | Subdivided | 1 (3) | 47 | 37 | 230 | 50x | 100 |
| M4G07 | Serial | 7 | Subdivided | 1 (4) | 46 | 33 | 299 | 42x | 100 |
| M4G08 | Serial | 8 | Subdivided | 1 (3) | 47 | 25 | 237 | 25x | 100 |
| M4G09 | Serial | 9 | Subdivided | 1 (3) | 47 | 26 | 244 | 26x | 100 |
| M4G10 | Serial | 10 | Subdivided | 1 (3) | 48 | 32 | 233 | 38x | 100 |
| T1G01 | Serial | 1 | *Turritis glabra* | 1 (3) | 47 | 31 | 247 | 37x | 100 |
| T1G02 | Serial | 2 | *Turritis glabra* | 1 (3) | 48 | 36 | 231 | 47x | 100 |
| T1G03 | Serial | 3 | *Turritis glabra* | 1 (3) | 47 | 36 | 237 | 48x | 100 |
| T1G04 | Serial | 4 | *Turritis glabra* | 1 (3) | 48 | 33 | 252 | 40x | 100 |
| T1G05 | Serial | 5 | *Turritis glabra* | 1 (3) | 47 | 37 | 235 | 49x | 100 |
| T1G06 | Serial | 6 | *Turritis glabra* | 1 (3) | 47 | 39 | 233 | 55x | 100 |
| T1G07 | Serial | 7 | *Turritis glabra* | 1 (3) | 47 | 38 | 232 | 53x | 100 |
| T1G08 | Serial | 8 | *Turritis glabra* | 1 (3) | 48 | 35 | 247 | 48x | 100 |

| Sample | Origin | Gen | Host | Pools (batches/pool) | MapQ | Dup (%) | Insert (bp) | Coverage | Mapped (%) |
|---|---|---|---|---|---|---|---|---|---|
| T1G09 | Serial | 9 | *Turritis glabra* | 1 (3) | 47 | 34 | 251 | 44x | 100 |
| T1G10 | Serial | 10 | *Turritis glabra* | 2 (3,3) | 47 | 47 | 238 | 88x | 100 |
| T2G01 | Serial | 1 | *Turritis glabra* | 1 (3) | 47 | 31 | 238 | 35x | 100 |
| T2G02 | Serial | 2 | *Turritis glabra* | 1 (3) | 48 | 33 | 261 | 40x | 100 |
| T2G03 | Serial | 3 | *Turritis glabra* | 1 (3) | 47 | 33 | 233 | 40x | 100 |
| T2G04 | Serial | 4 | *Turritis glabra* | 1 (3) | 48 | 37 | 232 | 50x | 100 |
| T2G05 | Serial | 5 | *Turritis glabra* | 1 (3) | 48 | 33 | 250 | 40x | 100 |
| T2G06 | Serial | 6 | *Turritis glabra* | 1 (3) | 47 | 27 | 247 | 29x | 100 |
| T2G07 | Serial | 7 | *Turritis glabra* | 1 (3) | 48 | 29 | 235 | 32x | 100 |
| T2G08 | Serial | 8 | *Turritis glabra* | 1 (3) | 47 | 35 | 234 | 44x | 100 |
| T2G09 | Serial | 9 | *Turritis glabra* | 1 (3) | 47 | 28 | 247 | 30x | 100 |
| T2G10 | Serial | 10 | *Turritis glabra* | 2 (3,3) | 47 | 41 | 240 | 64x | 100 |
| T3G01 | Serial | 1 | *Turritis glabra* | 1 (3) | 47 | 28 | 236 | 32x | 100 |
| T3G02 | Serial | 2 | *Turritis glabra* | 1 (3) | 47 | 24 | 233 | 23x | 100 |
| T3G06 | Serial | 6 | *Turritis glabra* | 1 (3) | 47 | 33 | 246 | 41x | 100 |
| T3G07 | Serial | 7 | *Turritis glabra* | 1 (3) | 48 | 28 | 247 | 30x | 100 |
| T3G08 | Serial | 8 | *Turritis glabra* | 1 (3) | 47 | 26 | 230 | 26x | 100 |
| T3G09 | Serial | 9 | *Turritis glabra* | 1 (3) | 48 | 37 | 233 | 51x | 100 |
| T3G10 | Serial | 10 | *Turritis glabra* | 2 (3,4) | 47 | 45 | 266 | 82x | 100 |
| T4G01 | Serial | 1 | *Turritis glabra* | 1 (3) | 47 | 32 | 252 | 39x | 100 |
| T4G02 | Serial | 2 | *Turritis glabra* | 1 (3) | 48 | 30 | 235 | 34x | 100 |

| Sample | Origin | Gen | Host | Pools (batches/pool) | Ma pQ | Dup (%) | Insert (bp) | Coverage | Mapped (%) |
|---|---|---|---|---|---|---|---|---|---|
| T4G06 | Serial | 6 | *Turritis glabra* | 1 (3) | 47 | 33 | 251 | 40x | 100 |
| T4G07 | Serial | 7 | *Turritis glabra* | 1 (3) | 47 | 32 | 252 | 37x | 100 |
| T4G08 | Serial | 8 | *Turritis glabra* | 1 (3) | 47 | 34 | 234 | 42x | 100 |
| T4G09 | Serial | 9 | *Turritis glabra* | 1 (3) | 47 | 34 | 250 | 43x | 100 |
| T4G10 | Serial | 10 | *Turritis glabra* | 1 (6) | 47 | 40 | 244 | 60x | 100 |

#### Sequence processing, mapping, and variant calling

Raw paired-end sequencing reads were processed and then aligned against the *S. flava* reference genome (sfla_v2) using the grenepipe Snakemake pipeline [21]. Adapter trimming and quality filtering were performed with fastp v0.20.1 [22] using the following parameters: detection of adapter sequences in paired-end mode, poly-G trimming (--trim_poly_g with minimum length = 10), 5’ and 3’ quality trimming (--cut_front, --cut_tail) with sliding windows of 4 bp and mean quality thresholds of 25 (5’) and 20 (3’), respectively, and a minimum read length of 60 bp. The first 6 and 8 bases of read 1 and read 2, respectively, were trimmed. Trimmed reads were mapped to the reference genome using BWA-MEM v0.7.17 [23] with default parameters. For samples with multiple sequencing runs, replicate BAMs were merged with samtools merge [23]. To minimize mapping artifacts, reads with a mapping quality below 20 were filtered using samtools view -q 20. Duplicate reads were identified and removed using Picard MarkDuplicates v2.27.5 [24]. Local realignment around indels was performed with GATK v3 RealignerTargetCreator and IndelRealigner [25].

Pooled allele frequencies were estimated from pooled whole-genome sequencing data using PoolSNP [26] applied to samtools mpileup summaries. Because sequencing depth, pooling structure, and biological sampling differed across the study, we generated multiple PoolSNP variant panels. Specifically, variants in founder pools were called separately to define high-confidence standing variation in the founding population. Variants in wild-collected pools were called separately to capture natural population variation. Variants in the experimental evolution time-series pools from generations 1-9 were called as a single panel because these libraries were sequenced to comparable depth and variants for the experimental generation 10 pools were called separately because replicate-merging increased coverage and altered the depth distribution relative to generations 1-9. For each panel, PoolSNP thresholds (max-cov, min-cov, min-count, min-freq, and miss-frac) were tuned to the empirical coverage distribution of that panel to control depth-related artifacts while retaining sensitivity appropriate to the sampling design.

Founder pools were called using a stricter minimum coverage and allele count (max-cov = 0.99; min-cov = 100; min-count = 50; min-freq = 0.10; miss-frac = 0), reflecting higher coverage and the goal of defining a conservative starting variant set. Time-series pools sequenced at similar depths (generations 1-9) were called with max-cov = 0.98, min-cov = 30, min-count = 10, min-freq = 0.01, miss-frac = 0. The generation-10 panel (including merged pools) used the same calling thresholds as the time series but was executed as an independent panel to avoid depth heterogeneity inflating false positives or distorting allele-frequency uncertainty. Wild and wild-merged pools were called with max-cov = 0.98, min-cov = 20, min-count = 5, min-freq = 0.05, miss-frac = 0, reflecting lower depth and the need to avoid spurious low-frequency calls.

For each pool set definition (e.g., founder pools, intermediate timepoint pools, wild pools, and merged final generation pools), per-chromosome mpileup files were generated from the final BAMs alignments with samtools mpileup using base- and mapping-quality filters (-Q and -q) and read-group handling as specified in the workflow configuration. Mpileups were produced separately for each genomic entry, enabling parallel variant calling and downstream filtering. Where pools were constructed from merged BAM alignments (e.g., pooled across replicate samples within a treatment x generation combination), mpileups were generated from the merged BAM set. Otherwise, mpileups were generated from the per-sample BAMs contributing to each pool. Per-entry mpileups were concatenated when needed to create unified mpileup inputs for a given panel. Variant discovery was performed with PoolSNP in pooled-discrete mode, executed independently for each genomic entry.

We retained only biallelic SNPs by removing any records with multiple alternate alleles prior to computing any pooled-sequencing corrections. Because read depth is an imperfect proxy for the number of independent chromosomes sampled in pooled sequencing (due to unequal individual contributions and technical resampling), analyses that required explicit correction of sampling variance used pooled-sequencing bias corrections based on effective sample sizes. For each pool and site, we estimated effective sample sizes (Neff) using known census pool sizes and PoolSNP output, producing per-pool correction files. These were combined into a single long-format correction table with sample metadata, which was used downstream both for variant filtering and, where required, for correcting read-depth fields in VCFs used by methods that assume an effective number of chromosomes rather than raw read depth.

Because many downstream analyses required a harmonized set of site coordinates, we implemented a two-stage filtering protocol: (1) define analysis-specific "final variant" position lists using allele-frequency behavior within each variant panel described above, then (2) subset the full VCFs to exactly those positions that form the intersection of all variant panels. The variant filtering steps, as described above, were performed within each panel separately so that the retained sites were supported by consistent evidence under a single sampling/coverage regime (founders, wild populations, and experimental time series).

We filtered PoolSNP-called variants to retain SNPs that were consistently polymorphic across time and replicated populations. First, for each SNP we removed observations whose allele frequency fell outside the polymorphic range of 0.01-0.99 (i.e., we retained rows with $0.01 < f < 0.99$). For each evolutionary replicate (defined as a unique pop × treatment combination), we then calculated the fraction of sampled generations in which that SNP remained within this frequency range and expressed this as the percentage of generations polymorphic. A replicate was considered to support a SNP if this percentage exceeded a user-defined threshold (e.g., ≥50% of generations). Finally, we retained only SNPs supported by at least a minimum number of replicates (e.g., ≥12 pop × treatment replicates), yielding a set of variants that were both polymorphic over time and reproducible across replicate populations.

Per-entry filtered variant tables were then aggregated into a single "final variants" list for each panel, which served as the canonical coordinate set for generating downstream statistical analyses. In parallel, we also generated an uncorrected-but-identically-filtered VCF version for statistical software that corrects for pooled-sequencing sampling variance internally (e.g., $F_{ST}$ estimates below).

Table S9. Software, versions, and parameters used in the experimental evolution pipeline. Versions reflect the environment used for all analyses reported in this study; references give the canonical citation. Pool-seq calling and correction parameters are detailed in the Methods text under "Sequence processing, mapping, and variant calling".

| Tool | Version | Use | Reference |
|---|---|---|---|
| fastp | 0.20.1 | Adapter trimming, quality and poly-G filtering of raw paired-end reads. | Chen et al. 2018 |
| BWA-MEM | 0.7.17 | Short-read mapping to the *S. flava* sfla_v2 reference genome. | Li & Durbin 2010 |
| samtools | 1.x | BAM sorting, merging, indexing, depth and flagstat-based QC. | Li 2011 |
| Picard MarkDuplicates | 2.27.5 | PCR-duplicate identification and removal. | broadinstitute.github.io/picard/ |
| GATK RealignerTargetCreator / IndelRealigner | 3.x | Local realignment of reads around indels prior to variant calling. | McKenna et al. 2010 |
| PoolSNP | latest | Pooled-sequencing biallelic SNP calling with per-panel parameter tuning (see Methods). | Kapun et al. 2021 |
| grenedalf | 0.6.2 | Per-pool π, Watterson's θ, Tajima's D in non-overlapping windows; per-pair $F_{ST}$. | Czech & Exposito-Alonso 2024 |
| ANGSD | 0.940 | Genotype-likelihood inference from low-coverage individual sequencing of the 128 founders. | Korneliussen et al. 2014 |
| PCAngsd | latest | Population-structure inference (PCA + admixture) from genotype likelihoods. | Meisner & Albrechtsen 2018 |

| Tool | Version | Use | Reference |
|---|---|---|---|
| ngsLD | latest | Linkage disequilibrium $r^2$ from genotype likelihoods (founder LD block + hotspot LD validation). | Fox et al. 2019 |
| BayPass | latest | Wild-pool genome-wide drift-covariance Ω, host-covariate $\beta_i$ / BF(dB), and C2 host contrast. | Gautier 2015 |
| haplovalidate | latest | Haplotype-block detection from temporal allele-frequency trajectories. | Otte & Schlötterer 2021 |
| cvtkpy | github.com/vsbuffalo/cvtk | Temporal-replicate covariance, G statistic, convergence correlation and their bias corrections. | Buffalo & Coop 2020 |
| poolSeq (R) | latest | Within-replicate $N_e$ estimation from pool-seq time series (Plan I). | Taus et al. 2017 |
| ACER (R) | latest | Cochran-Mantel-Haenszel test for time-series allele-frequency change. | Spitzer et al. 2020 |
| lme4 (R) | latest | Linear and generalised linear mixed models (realised heritability; per-SNP GLMM with parametric-bootstrap LRT). | Bates et al. 2015 |
| diamond blastp | 2.1.10 | *S. flava* proteome vs *D. melanogaster* reference proteome alignment for Muller-element assignment. | Buchfink et al. 2021 |
| *S. flava* reference (sfla_v2) | — | Reference genome assembly used for read mapping and variant calling. | Published |
| *D. melanogaster* reference proteome | UniProt UP000000803 | *D. melanogaster* proteome target for Muller-element ortholog assignment. | UniProt 2023 |
| Drosophila_melanogaster.gene_info (NCBI) | — | Dmel gene to chromosome lookup for translating diamond hits to Muller A-F. | NCBI Gene |

### Genomic starting conditions

#### Population structure of experimental founder flies

The founder colony used for the experimental evolution time series was established by combining wild-caught individuals from *B. vulgaris* (host of **B**) and *T. glabra* (host of **T**) hosts and expanding the mixed pool for two generations prior to the start of the experiment. Residual host-source stratification in the colony would confound inference of drift and selection across subsequent generations. To address this, we individually sequenced 128 founder flies and characterized ancestry structure in the starting population. We then mapped low-coverage whole-genome reads (~0.5x) to the *S. flava* reference assembly and processed results using duplicate-marking and recalibration steps (see methods above). Given our low-coverage, we estimated genotype likelihoods with ANGSD [27] (v0.940) using the SAMtools likelihood model (-GL 1), retaining sites with -SNP_pval 1e-6, minimum base and mapping qualities of 20, a minimum of 64 individuals with data (half the cohort) and a minimum minor-allele frequency of 0.01. We inferred population structure from the resulting BEAGLE file using PCAngsd [28] and computed a sample-by-sample covariance matrix and iteratively estimated admixture proportions with an ancestry-component number K from 2 to 8 (-e 10 -iter 500 -maf 0.05 -admix -admix-K). To test whether and to what extent founders sourced from the **B** host and **T** host expansion plants in the laboratory retained residual population structure driven by the host plant, we used K = 2 and compared the first admixture component between the two groups.

We found no residual **B** or **T** host-plant associated residual population structure, suggesting that two generations of admixture during colony expansion in the laboratory was sufficient to generate a relatively well-recombined genetic background. On the principal-component axes inferred from the genotype-likelihood covariance matrix, samples formed a single cloud with no separation between **B**, **T**, or flies raised in cages with both **B** host and **T** host plants (**B+T**) (Figure S12B). A maximum of 3.16% of total variance could be explained by PC1, with PC2 and PC3 at 2.79% and 2.64%, suggesting weak population structure. In addition, admixture proportions were uniform across individuals at every value of K (Figure S12A). At K = 2 the mean proportion of the first ancestry component was 0.43 in **B** flies and 0.50 in **T** flies, with no significant difference (Welch's t = -1.24, P = 0.22). We found that the variation explained by principal components was stable across most consecutive K transitions (max per-component correlations ≥ 0.95 for 28 of 35 transitions), with minor reshuffling at K = 2 to 3 (second component max r = 0.71), which was expected as new components were introduced to a weakly structured dataset. PCAngsd reported non-convergence at every K tested, given the very low total eigenvalue mass on the leading PCs and the lack of an alternative signal. We interpreted non-convergence here as indicating the absence of strong population structure. These results suggest that founders represented a single well-mixed ancestral experimental pool, justifying our approach of considering the starting frequencies of each experimental replicate as drawn from the same common prior.

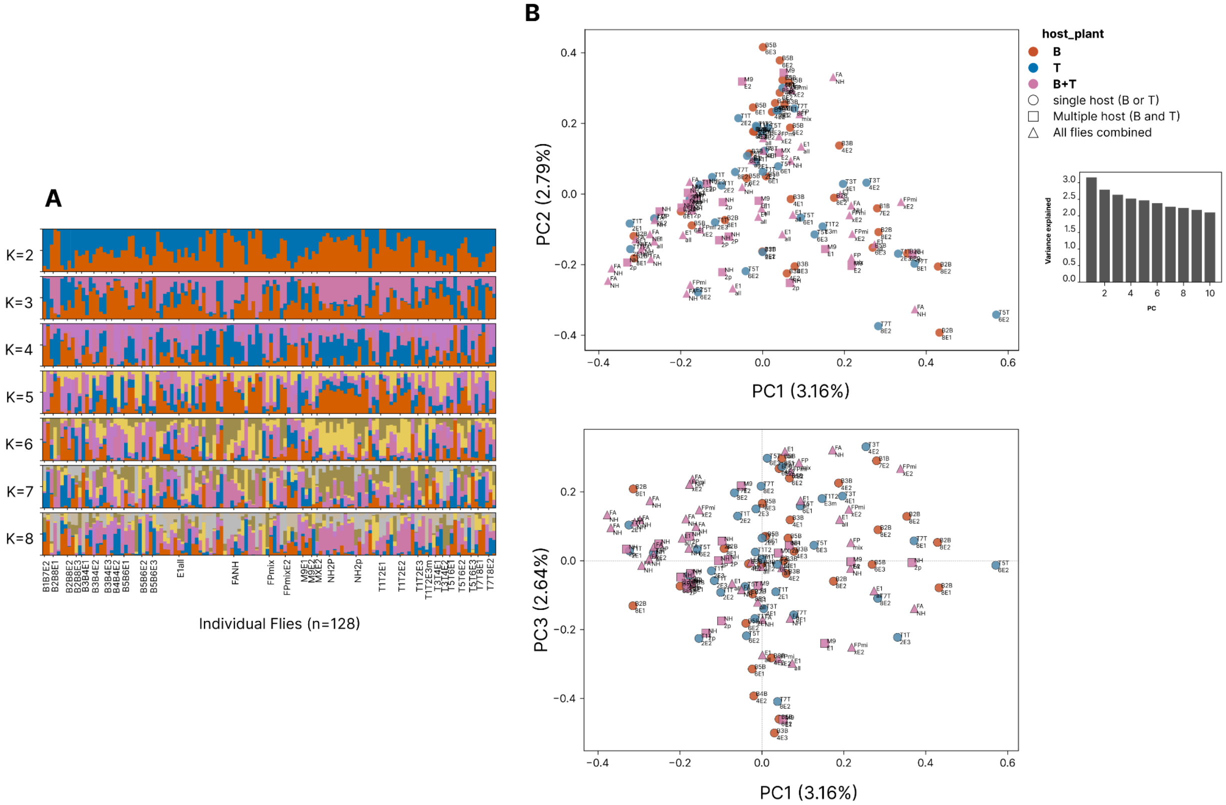

Figure S12: Population structure of the 128 individually-sequenced *S. flava* founders. (A) Stacked-bar admixture proportions for K = 2 through K = 8. Components were ordered by PCAngsd and colored consistently across K panels. (B) Principal components PC1 vs. PC2 (top) and PC1 vs. PC3 (bottom) from the PCAngsd genotype-likelihood covariance matrix. Variance explained is shown per axis. Points are colored by inferred host source (host **B** = *B. vulgaris*, host **T** = *T. glabra*, **B+T** = multiple host plant cages) and marker shape encodes collection pool type. The inset bar panel in the upper PCA panel shows the proportion of variance explained by the first 10 PCs. No separation between host-source groups is apparent.

#### Allele-frequency-trajectory PCA across the experimental time series

Having confirmed that the founder population formed a single well-mixed source pool with no detectable host-of-origin structure, we projected the cage-level evolutionary trajectories into a low-dimensional summary of overall genomic divergence to address how the three selection regimes drove genomic change across our replicate experimental populations. We performed principal-component analysis on the per-sample allele-frequency matrix across the 22 pool-seq libraries that span the experimental time series. These included the four founder pools (F1-F4 at $t_0$) plus the 12 $t_{10}$ cages (B1-B4, T1-T4, M1-M4), including an extended set including all available intermediate-generation pools ($t_1$, $t_2$, $t_6$, $t_7$, $t_8$, $t_9$ for all 12 cages, and $t_3$-$t_5$ for B1, B2, T1, T2, note that cages 3 and 4 were not sequenced at t3-t5 due to limited sample availability). Per-sample allele frequencies were extracted directly from the per-sample AD field of the joint VCF as alt / (ref + alt). We retained sites with per-sample read depth between 10 and 500 in all kept samples and a mean allele frequency in [0.01, 0.99]. For each retained SNP, we centred the per-sample row by the SNP-specific mean allele frequency across samples and scaled by the binomial standard deviation $\sqrt{(p_i(1 - p_i))}$, the standard for population-genetic allele-frequency PCA [29]. The resulting (samples x SNPs) matrix was decomposed using sklearn.decomposition.PCA with svd_solver = "full" and the first ten principal components were retained for downstream analysis. Variance explained by the top three axes was 2.7%, 2.7%, and 2.6% respectively, consistent with weak overall structure and the role of per-cage genetic drift and weak per-treatment differentiation on overall variance. Per-cage trajectories were drawn as 3D figures where per-generation positions were connected in order of generation using coordinated from PC1-PC3, with arrowheads at the $t_{10}$ endpoints.

To test whether **B+T** cages at $t_{10}$ diverged from a midpoint between **B** and **T** (the prediction under arithmetic-mean fitness across the two host plants), we computed for each treatment the centroid of its four $t_{10}$ cages. We used PC1-PC10 coordinates and then measured the Euclidean distance of the **B+T** centroid from the midpoint of the **B** and **T** centroids using the formula $d = c_{B+T} - ½(c_B + c_T)$. We additionally computed the angle θ between the displacement vector and the line connecting $c_B$ and $c_T$, where $\theta = 0°$ corresponded to the **B+T** centroid sitting on the **B** vs. **T** axis (the arithmetic-mean prediction). The observed values were d = 66.2 units and θ = 121°, indicating that the **B+T** cages at $t_{10}$ occupied a region of PC space that was neither between **B** and **T** on a linear interpolation nor consistent with the arithmetic-mean expectation. Instead, **B+T** represented a distinct evolutionary trajectory under a multiple-host subdivided selection environment.

To assess statistical significance, we generated a permutation null by treatment-label shuffling where each of $10^4$ permutations included the PC scores from the 12 $t_{10}$ cages randomly partitioned into three groups of four. The displacement statistic d was recomputed for the resampled **B+T** group against the resampled **B** and **T** group centroids. The empirical p-value was the fraction of permutations with d matching or exceeding the observed value ($p = 0.055$).

#### Standing variation, colony bottleneck, and diversity rebound

We tracked how genetic diversity evolved across three stages of the experiment that included the six wild pools, four founder pools (F1–F4, 80 flies each) drawn from the expanded founding colony, and twelve G10 pools (B1–B4, T1–T4, M1–M4, each combining two independent 80-fly pools, 160 flies) sampled from the twelve experimental cages. Each was previously seeded with 500 flies from the same founding colony and evolved for ten generations under one of three selection regimes (**B**, **T**, and the multiple-host Levene treatment **B+T**). All 22 pools were aligned to the same *S. flava* reference (sfla_v2.fa) and restricted to the same 812,670 biallelic SNPs.

We estimated three per-pool genome-wide diversity statistics, including π ($\theta_\pi$), Watterson's $\theta_W$, and Tajima's D, in 390-kb non-overlapping windows using grenedalf diversity (v0.6.2), which implements pool-seq normalisation for Tajima's D. We applied per-sample filters (minimum allele count 2, minimum read depth 4, maximum read depth 500) and averaged each statistic over valid SNPs within each window. Pool sizes (50 for wild, 80 for founder, and 160 for G10 pools) were used for variance normalizations so that D values could be made comparable across heterogeneous pool sizes. We first estimated how much standing variation the wild *S. flava* source population carried. Per-pool genome-wide medians across the six wild pools spanned $\theta_\pi$ ~ 0.24 to ~0.25, $\theta_W$ ~0.29 to 0.33, and Tajima's D ~ -0.7 to -1.3 (Figure S13). All six pools sat below D = 0 on average, which is consistent with the signature of an excess of low-frequency variants and a recent population expansion or weakly purifying selection. We used paired **B**-vs-**T** Wilcoxon tests within each site to detect significant shifts in all three metrics at every site ($p < 10^{-9}$), but the direction varied. PS and AV had slightly higher θ in **T**-host pools, while RM had higher $\theta_W$ in the **B**-host pool and more negative D. These within-site effects were small (effect < 0.03, $|\Delta D| < 0.3$) relative to the wild-vs-founder contrast we describe below.

We next estimated how much of the wild standing variation was retained in the founding colony from which all twelve experimental populations were drawn. The wild-vs-founder contrast isolates the combined effect of colony establishment and two generations of expansion (Figure 1). Three patterns distinguish the founder pools from the wild pools. First, $\theta_\pi$ shifted up (wild ~0.246, founder ~0.270) and the four founder pools collapsed onto an almost identical distribution (per-pool median SD = 2E-4, vs 7E-3 across the six wild pools), confirming both that F1–F4 sampled the same underlying allele frequency spectrum and that the mean per-site heterozygosity was mostly preserved. Second, $\theta_W$ dropped (wild ~0.300, founder ~0.223, ~26% reduction). Third, Tajima's D flipped sign, from -0.86 ± 0.12 across the wild pools (excess of low-frequency variants, consistent with population expansion) to +0.83 ± 0.08 across F1–F4 (deficit of low-frequency variants relative to common ones). The asymmetric loss of low-frequency variation captured by $\theta_W$ and the sign flip in Tajima's D reflected the same event from two different perspectives and confirmed the bottleneck. We concluded that the founding of the colony removed a substantial fraction of rare wild alleles while leaving intermediate-frequency variation largely intact.

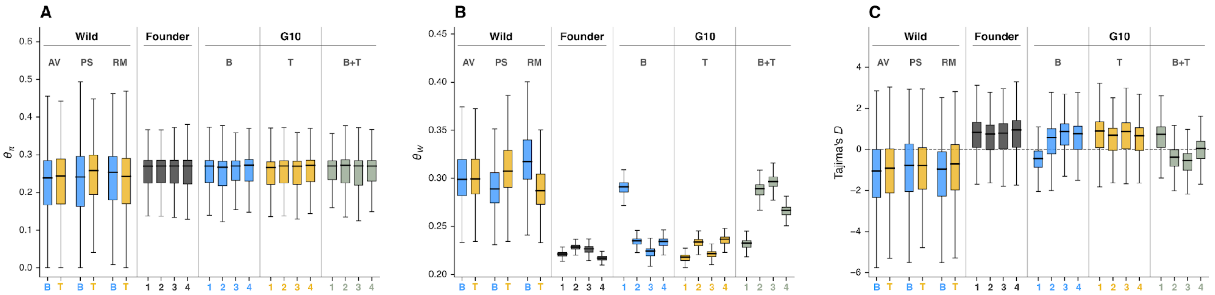

**Figure S13.** Per-pool genetic diversity across the wild, founder, experimental G10 samples of *S. flava*. Per-window estimates of $\theta_\pi$ (A), Watterson's $\theta_W$ (B), and Tajima's D (C) for all 22 pool-seq libraries, shown as boxplots over the 934 non-overlapping 390-kb genomic windows. Within each panel, pools are grouped left-to-right as: six wild pools at three New England localities (AV, PS, RM with paired **B**-host or **T**-host samples at each site), four founder pools (F1–F4, independent 80-fly subsamples of the expanded founder colony), and the twelve G10 pools grouped by selection regime (G10 **B**, G10 **T**, G10 **B+T**; four replicate cages each). Pool-size corrections (haploid n = 50 for wild, 80 for founder, 160 for G10) were applied via the pool-seq normalisation for all three statistics. **B**-host pools in blue, **T**-host pools in yellow, founder pools in dark grey, G10 **B+T** (Levene multiple-host) pools in light green. Dashed horizontal line marks Tajima's D = 0.

We then estimated how genome-wide diversity evolved over ten generations of experimental evolution under the three selection regimes. $\theta_\pi$ was effectively unchanged from the founder value across all three treatments (**B**, **T**, and **B+T** all ~0.270), indicating that per-site heterozygosity was similarly preserved through ten generations regardless of the particular selection pressures imposed. $\theta_W$ and Tajima's D diverged along the single-host vs multiple-host axis. The **B** or **T** treatments retained the founder signature in both metrics (**B**: $\theta_W$ = 0.246 ± 0.031, D = +0.32 ± 0.55; **T**: $\theta_W$ = 0.228 ± 0.009, D = +0.63 ± 0.13). This resulted in the retention of the rare-variant deficit that would characterize a recent bottleneck. The Levene multiple-host treatment (**B+T**), in contrast, partially reversed the bottleneck signature where $\theta_W$ climbed to 0.271 ± 0.029, closer to the wild median of 0.30 than to the founder value of 0.22. Tajima's D in **B+T** flipped back to -0.11 ± 0.52. The elevated between-replicate variance in **B+T** reflected cage-level heterogeneity. Three of the four **B+T** replicates (M2, M3, M4) showed this rebound while M1 retained the founder state. A parallel single-cage pattern appears in the **B** treatment (B1, with $\theta_W$= 0.289 and D = -0.48, an endpoint outlier where the $t_1$-$t_9$ slope is flat). This per-cage heterogeneity was examined in the next paragraph. At the treatment level, the multiple-host Levene environment was therefore the only environment that produced a measurable shift in the rare-variant fraction over ten generations, which supports the theoretical prediction that spatially varying selection maintains more intermediate-frequency variation than directional selection on a single host.

To distinguish a treatment-wide rebound in genetic diversity from per-cage anomalies and to disentangle the rebound signal from the depth-dependent rare-variant ascertainment that affects the G10 endpoint sequencing, we examined how diversity evolved during the experiment rather than only at its endpoints. Sequencing depth differed between the intermediate generations (G1–G9) and the founder/G10 endpoint libraries, so $\theta_W$ at the endpoints was partially inflated and could not be directly compared to intermediate generations. Nonetheless, across G1–G9, the per-replicate $\theta_W$ slopes within the **B+T** treatment were positive across all four cages (range +0.0016 to +0.004 per generation), suggesting that the diversity rebound under multiple-host selection is treatment-wide. The **B** treatment showed a much wider spread of per-replicate slopes (-0.001 to +0.004/gen). The **T** replicates remained flat throughout, consistent with no rebound under the **T** treatment. Tajima's D showed the same pattern from the opposite sign where there were negative slopes in **B+T** (and B1), flat in **T** and B2-B4 (Figure S15).

We then cross-referenced the per-replicate $\theta_W$ slopes with the temporal-covariance $G_{k\geq 2}$ statistic from the convergent-selection scan. We found that the within-treatment rebound outliers (B1, M2, M3, M4) overlapped with replicates showing positive temporal-selection signal in $G_{k\geq 2}$ (Spearman $\rho = +0.60$, $p = 0.039$ across the 12 G10 cages). B1 and M3 carried both the highest $G_{k\geq 2}$ values and the steepest $\theta_W$ slopes. The exception was M2, with an elevated $\theta_W$ slope but a $G_{k\geq 2}$ value that was negative through G08 and only weakly positive at G09 (Fig. S21), and an extreme outlier signature on PC4 of the genome-wide ordination of raw allele frequencies, driven by a single late allele frequency change instead of a consistent temporal response. These patterns suggest that the multiple-host Levene environment generally produced a consistent treatment-wide reduction of the bottleneck signature, which is again (as above) consistent with the theoretical maintenance of intermediate-frequency variation under spatially varying selection. The parallel rebound of diversity observed in B1 reflected cage-specific selection or drift on a single-host background but was identifiable when intermediate-generation trajectories were examined per-replicate.

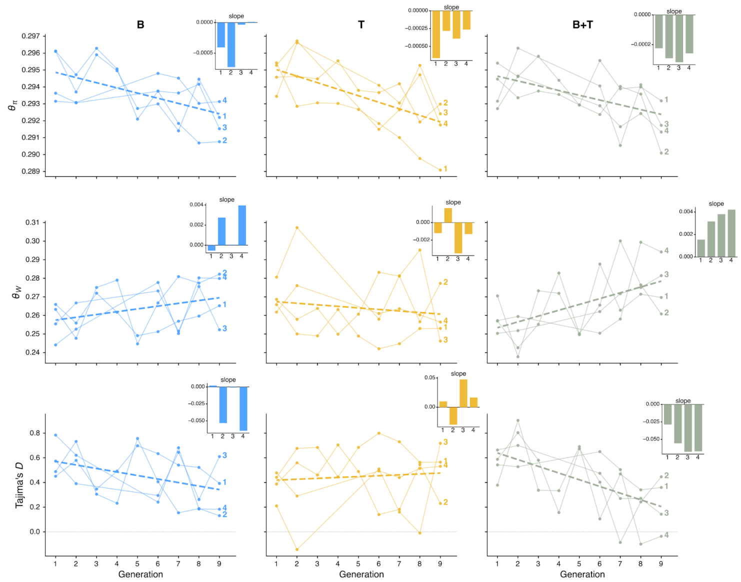

**Figure S15.** Per-replicate diversity trajectories of *S. flava* across intermediate generations G1–G9. Per-pool window medians of $\theta_\pi$ (top row), Watterson's $\theta_W$ (middle row), and Tajima's D (bottom row) for each of the twelve experimental cages, sampled at generations G1–G9. Columns separate the three selection regimes: **B**, **T**, and **B+T**. Each panel shows the four replicate cages of that treatment as lines. The numeric label (1–4) at the right edge of each trajectory identifies the replicate. Diversity statistics were estimated in 200-kb non-overlapping windows with pool sequencing correction. Window restricted to the 1,192 windows common to all twelve cages. The dashed line within each panel represents linear regression combining medians from all replicates. The inset bar chart in the upper right of each panel represents the per-replicate regression slope (one bar per cage). The dotted horizontal line in the bottom row represents Tajima's D = 0. Founder (G0) and G10 endpoints were excluded from this figure because they were sequenced to different average depth than the intermediate generations.

In summary, the wild *S. flava* source population carried high genome-wide standing variation with negative Tajima's D across all six wild pools, a signature of recent population expansion. Second, the founding of the laboratory colony imposed major bottlenecking on this wild genetic variation. Heterozygosity was largely preserved ($\theta_\pi$), but we found a 26% reduction in $\theta_W$, and a sign flip of Tajima's D. Third, after ten generations of selection from the founder population, only the multiple-host Levene treatment (**B+T**) consistently reversed the bottleneck signature, showing positive $\theta_W$ slopes and negative Tajima's D slopes across all four **B+T** replicates. The single-host **B** and **T** treatments preserved the founder state, with one cage-specific exception (B1) whose per-replicate trajectory and elevated $G_{k\geq2}$ temporal-covariance score suggested a strong cage-level effect. In addition we found some agreement of the $G_{k\geq2}$ temporal-covariance statistic with diversity shift metrics. These results are consistent with the theoretical prediction (Levene Model) that subdivided spatially varying selection across hosts maintains intermediate-frequency variation that directional selection purges on a single host.

#### Idiosyncratic rare-variant retention signal

The cage-level rare-variant retention pattern we reported in the main text is that **B+T** cages diverged into idiosyncratic subsets of retained rare variants while single-host **B** and **T** cages converged on similar variant subsets across replicates. We quantified this at finer genomic resolution (100-kb) by using non-overlapping 100-kb windows on scaffold chr_ScDA7r2_110 (48.2 Mb, Scaffold 110). We counted, per window and per pool, the number of SNPs that were polymorphic in that pool (the per-pool segregating-site count). The 100-kb resolution was chosen for the per-replicate variance test because the smaller window size resolved spatial heterogeneity at chromosome-scale scaffolds without degrading per-window estimation precision. Scaffold 110 was used as the focal scaffold because it was the largest chromosome-arm-scale scaffold not affected by the scaffold 597 chromosomal fusion and not the focal candidate scaffold of Section 2 (scaffold 439), therefore representing a non-candidate genome-wide background region. The same analysis run on every chromosome-scale scaffold reproduced the same pattern.

For each treatment $T \in \{\mathbf{B}, \mathbf{T}, \mathbf{B{+}T}\}$ and each window w, we computed the standard deviation of the per-replicate per-window segregating-site count across the four replicate cages of that treatment at $t_{10}$:

$SD_{replicate,T,w} = SD_{r \in \{1,2,3,4\}}(n_{segregating,T,r,w})$. The mean across windows on scaffold 110 of these per-window between-replicate SDs was 2.32 (**B**), 11.20 (**B+T**), and 2.36 (**T**), corresponding to approximately 5-fold higher between-replicate cage-level variance in segregating-site density under multiple-host selection than under either single-host environment. We then tested whether the per-window SD distributions differed across treatments via two pairwise one-sided Mann–Whitney U tests on the per-window SD distributions: **B+T** versus **B**, and **B+T** versus **T**. The test statistic is the rank-sum of **B+T** SD values among the pooled (**B+T**, **B**) or (**B+T**, **T**) windows. The p-values were computed under the asymptotic Mann-Whitney null using scipy.stats.mannwhitneyu with alternative = "greater", since the prediction is that **B+T** SD exceeds the single-host SD. Both contrasts gave $p < 10^{-10}$ at this scaffold.

The biological interpretation of this 5-fold variance excess is described in the main text. Each **B+T** cage retained a slightly different subset of rare variants (different cages drove different rare-variant subsets up in frequency under coarse-grained subdivided selection) while **B** and **T** cages purged similar rare-variant subsets across replicates under directional selection on a single host. This pattern is consistent with a process in which each **B+T** cage's evolutionary trajectory at any given chromosomal interval was dominated by stochastic outcomes of competition between the two host-favored allele subsets across the ten generations of selection, while at single-host loci all four replicate cages converged on the same allele-frequency response.

### Noise floor and effective population size

#### Variance decomposition of technical and biological allele frequency change

Allele frequency change in a pool-sequencing time series reflects three distinct sources of variation. The first is technical noise from the finite sampling of pooled individuals and from library preparation and sequencing, which sets minimum thresholds for detecting allele frequency change due to biological factors. The second is drift variance, governed by the effective population size $N_e$ of each replicate cage. The third is selection variance, the shared directional signal detectable when independent replicate populations evolve in parallel. We separated these three sources by first measuring the technical noise directly from data, then using it to correct the between-replicate and within-replicate estimators of $N_e$, and finally determine whether and to what extent a residual signal of selection remains above the drift baseline.

#### Technical noise

Under ideal conditions, pooled sequencing of $N_{pool}$ diploid individuals at read depth d has a per-site variance,

$$\mathrm{Var}(\hat{p} \mid p) = p(1-p) \cdot \left[\frac{1}{2N_{pool}} + \frac{1}{d}\right]$$

However, unequal DNA contribution across individuals inflated the pool-sampling component above this binomial expectation. We characterized the overdispersion with an effective pool size $N_{eff}$, which replaces $N_{pool}$ in the formula above and which we estimated from three independent analyses.

We first estimated overdispersion of allele frequencies due to pooled sampling and pooled sequencing by considering the four founder pools (F1-F4 at G00) as independent draws of 80 flies each from the same large admixed colony (see above for founder admixture analysis), with no drift between them. Between-founder-pool variance at a given site therefore reflected only pool sampling plus sequencing variance. We compared the observed between-pool variance at each of ~776,000 SNPs to the expectation from binomial pooling with $N_{pool} = 80$:

$$\varphi = \frac{\text{observed between} - \text{pool variance}}{p(1-p)/(2N_{pool})}$$

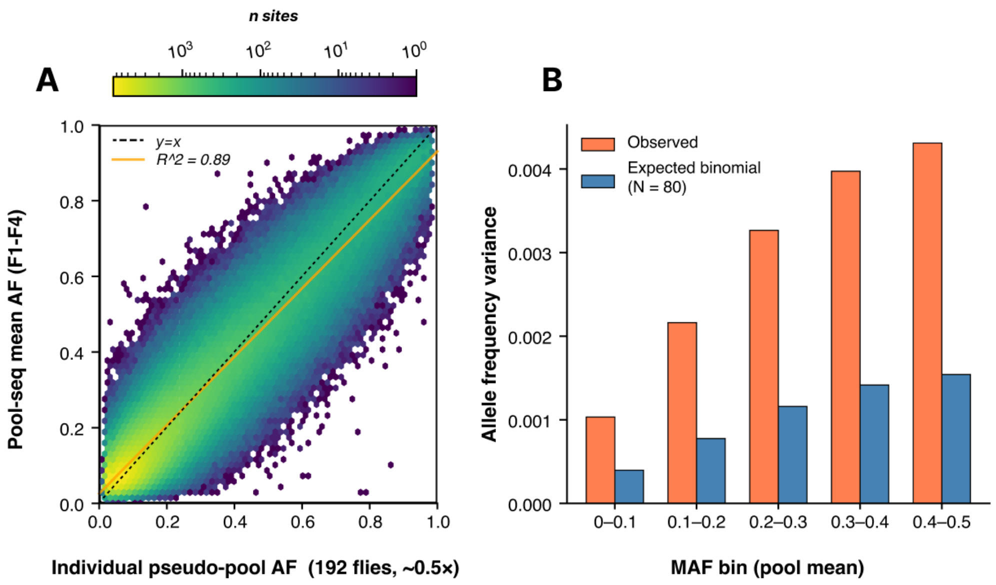

Figure S18. Pool sequencing overdispersion. (A) Individual pseudo-pool allele frequency (128 founder flies sequenced at ~0.5x each, aggregated with ANGSD) versus pool-seq mean allele frequency across the four founder pools (F1-F4) ($R^2$ = 0.89, slope = 0.90). Dashed black line: $y = x$. (B) Observed between-founder-pool allele frequency variance (orange) versus expected binomial variance at the pool size $N_{pool}$ = 80 (blue), stratified by pool-mean minor allele frequency. The consistent ~2.8x inflation across MAF bins gives a genome-wide overdispersion factor φ = 2.78, corresponding to N = 80/~29 diploids.

We then employed a second estimate using the k = 1 off-diagonal of the temporal-replicate covariance matrix (Buffalo and Coop 2019, 2020, see section below). Adjacent intervals $\Delta p_{t\to t+1}$ and $\Delta p_{t+1\to t+2}$ within a single replicate share the measurement at generation t +1, so their covariance equals the negative of the sampling variance at that shared time point:

$$\mathrm{Cov}(\Delta p_{t\to t+1}, \Delta p_{t+1\to t+2}) = -\sigma^2_{sampling}$$

We found that the k = 1 covariance is 1.72x more negative than expected from binomial pooling at $N_{pool}$ = 80 and the measured per-sample read depths, giving $N_{eff}$ ~24 diploids, slightly lower than the founder estimates. This is consistent with evolved-sample libraries having slightly more overdispersion than the founder libraries.

These two approaches converge on $N_{eff}$ ~24-32 diploids. We used this information to inform downstream corrections and note that these values are also supported by the intercept of the noise-corrected F regression conducted in the next section. A side-by-side comparison of the $N_e$ estimates from all methods appears at the end of this section.

#### Drift variance and effective population size

We next estimated the effective population size ($N_e$) of each treatment from the rate of divergence among replicate cages over time. Under genetic drift, Wright's F statistic between the k=4 replicates grows linearly with generation at a rate 1/2($N_e$), so that regressing F against generation t and taking the slope recovers effective population size [30,31]. We computed F at each generation using the ratio-of-averages estimator,

$$F(t) = \frac{\text{mean}_{\text{sites}}[\text{Var}(p_{1..k}(t))]}{\text{mean}_{\text{sites}}\left[\underline{p}(t)(1-\underline{p}(t))\right]}$$

with the unbiased $(k-1)^{-1}$ between-replicate variance. We produced point estimates of $N_e$ by fitting the regression to the six generations with complete 4-replicate sampling (G01, G02, G06, G07, G08, G09), which produced estimates of 205, 420, and 165 for treatments **B**, **T**, and **B+T** respectively. These values suggest that **T**'s effective size is approximately 2x that of **B** and **B+T**. However, because pool-sequencing sampling noise contributed in a depth-dependent way to the observed between-replicate variance, we conducted this analysis using a pool-sequencing aware correction. For a replicate r at a site with true population frequency $p_r$, the observed frequency $\hat{p}_r$ is influenced by the pooling of $N_{pool}$ diploids ($2N_{pool}$ gametes), with $d_r$ reads drawn from that pool. Two independent pool-seq diagnostics in this dataset, including overdispersion of the four founder pools at G00 ($\varphi = 2.78$, $N_{eff} = 29$) and the k = 1 diagonal of the temporal-replicate covariance matrix ($N_{eff}$ = 24, see next section below), indicated that the effective contributing pool size per library is about 29 diploids. With this effective pool size, the per-replicate sampling variance of $\hat{p}_r$ is

$$\mathrm{Var}(\hat{p}_r \mid p_r) \;\approx\; p_r(1 - p_r) \cdot \left[\frac{1}{2N_{eff}} + \frac{1}{d_r}\right]$$

Because the k replicates were sampled independently, this variance adds to the biological between-replicate variance. The expected F under sampling noise with no drift is then,

$$F_{noise}(t) \;\approx\; \frac{1}{2N_{eff}} + \mathrm{mean}_{r,\,sites}\left[\frac{1}{d}\right](t)$$

where $\mathrm{mean}_{r,\,sites}[1/d](t)$ represents the mean of the reciprocal depth $1/d_{r,i}$ taken across the four replicates r and across sites i at generation t, with the per-site contribution weighted by $\underline{p}(1 - \underline{p})$ to match the ratio-of-averages form of F. The $1/(2N_{eff})$ term is constant across generations and shifts only the regression intercept. The $1/d$ term tracked per-generation depth and introduced a bias on the slope whenever depth was not constant over time.

We found that the mean per-pool sequencing depth varied across the six 4-replicate generations and the direction of variation differed among treatments as follows (Fig. S14):

**Table S11.** Per-treatment mean sequencing depth across the six 4-replicate generations (G01, G02, G06, G07, G08, G09) used in the F-regression $N_e$ and temporal covariance analyses. Depth in **T** increased ~19% across this window while depth in **B** and **B+T** decreased ~16-22 %, generating a depth confound in the raw F-regression that motivated the per-generation noise correction (Methods).

| **Generation** | **B** | **T** | **B+T** |
|---|---|---|---|
| G01 | 87 | 72 | 85 |
| G02 | 88 | 73 | 103 |
| G06 | 75 | 84 | 76 |
| G07 | 84 | 76 | 70 |
| G08 | 68 | 78 | 62 |
| G09 | 73 | 86 | 66 |

Depth in **T** increased from G01 to G09 (+19%), depth in **B** decreased (-16%), while depth in **B+T** decreased (-22%). The effect of these patterns is that $F_{noise}(t)$ shrunk over time in **T** and grew over time in **B** and **B+T**. Considering the original F regression, this biases **T**'s slope flatter (inflating $N_e$) and **B**'s/**B+T**'s steeper (deflating $N_e$).

We recomputed F using a single common site filter (minimum depth ≥ 10 in all 4 replicates at every 4-replicate generation of every treatment, and minor allele frequency ≥ 0.05 based on the pooled founder mean, 762,812 sites passing) and subtracted the per-generation, per-site weighted expected sampling contribution:

$$F_{bio}(t) \;=\; F(t) \;-\; \frac{\mathrm{mean}_{\mathrm{sites}}\left[\,\underline{p}(1-\underline{p})\left(\frac{1}{2N_{eff}} + \mathrm{mean}_{r}\left[\frac{1}{d}\right]\right)\right]}{\mathrm{mean}_{\mathrm{sites}}\left[\,\underline{p}(1-\underline{p})\,\right]}$$

and re-regressed $F_{bio}(t)$ on t. We found that after correction, the regression intercepts for **B** and **B+T** were about zero (-0.0009 and -0.0016 respectively), and for **T** it was small but nonzero (0.0030) (Fig. S14). Because the four founder pools were sampled independently from a single large, admixed colony, the expected biological between-replicate F at G00 was approximately 0, and the observed raw intercepts of 0.028-0.036 were almost entirely sampling noise. $R^2$ of the linear regression improved from 0.936/0.894/0.935 to 0.956/0.966/0.958 for **B**/**T**/**B+T** after correction, suggesting that removing this sampling component of F improved estimates of drift variance over time. The **T** effective size shrank from a 2x inflation relative to **B**/**B+T** to about 1.5x.

**Table S7.** Noise-corrected effective population size ($N_e$) estimated from between-replicate F-regression, before and after subtracting the per-generation expected pool-sequencing sampling noise. Bracketed values are 95 % block-bootstrap confidence intervals. After correction, **T**'s apparent ~2x inflation in $N_e$ relative to **B** and **B+T** shrinks to ~1.5x.

| Treatment | Raw $N_e$ [95% CI] | Corrected $N_e$ [95% CI] |
|---|---|---|
| **B** | 205 [202, 207] | 242 [238, 245] |
| **T** | 420 [412, 432] | 347 [341, 355] |
| **B+T** | 165 [164, 167] | 219 [216, 221] |

We conducted block-bootstrapping to generate confidence intervals using 100 kb linkage windows but did not account for the finite number of biological replicates (k = 4) or the finite number of founder pools. Therefore this approach represents a lower bound on the true uncertainty.

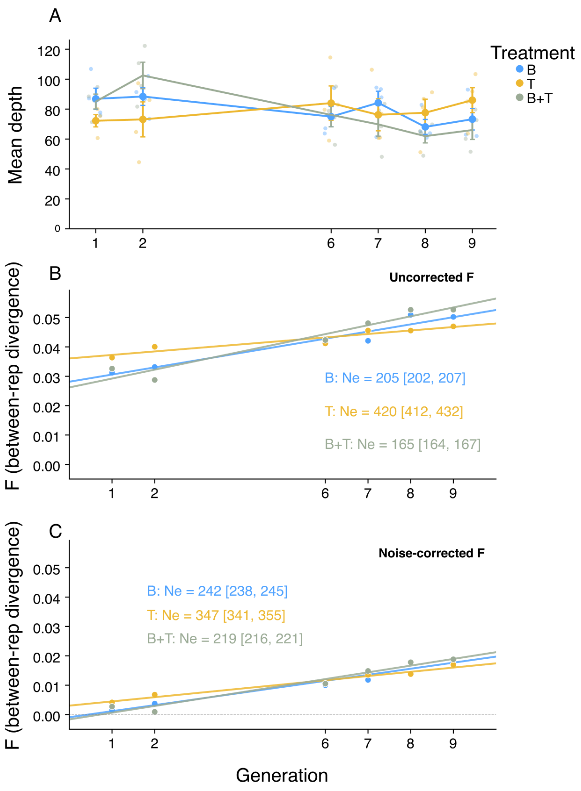

**Figure S14.** (A) Mean sequencing depth per pool for each treatment across the six generations with complete 4-replicate sampling. Points show replicate-level depths. Error bars show ±1 SEM across the four replicates. **T** depth increases over time while **B** and **B+T** decrease, generating a depth confound in the raw F regression. (B) Raw between-replicate F versus generation with linear regression fits. $N_e$ labels give point estimates and 95% block-bootstrap confidence intervals. (C) Same data after subtracting the per-generation expected pool sequencing sampling noise. After correction, intercepts are near zero for **B** and **B+T**, $R^2$ improved for all three treatments, and the treatment differences in $N_e$ partially decreased.

#### Within-replicate temporal estimates of $N_e$

As an independent estimate of $N_e$, we applied the poolSeq estimator [32]. We used 1000-SNP windows, G01 to G09 (t = 8 generations) and calculated the median $N_e$ across windows per replicate and then the mean across replicates. Under the $N_{pool}$ = 80 pool size, this method reports that all three treatments have $N_e$ ~100, about half of that reported by the F-regression. However, the effective pool size determined from founder overdispersion and temporal covariance was $N_{eff}$ ~29. With $N_{pool}$ = 29, poolSeq $N_e$ approximately doubled for all treatments (Fig. S16):

| Pool size | B | T | B+T |
|---|---|---|---|
| 80 | 100 | 100 | 102 |
| 29 (effective) | 208 | 211 | 226 |
| 24 (cvtkpy k=1) | 312 | 318 | 329 |

These results were robust across estimator variants (Plan I vs Plan II) and time intervals (G01-G09, G01-G06, G02-G09). We found that (1) after pool-size correction, the poolSeq and F-regression estimates converged to the same order of magnitude $N_e$ ~200–350 and (2) poolSeq gives identical $N_e$ across all three treatments regardless of pool size, while F-regression showed a residual **T** elevation.

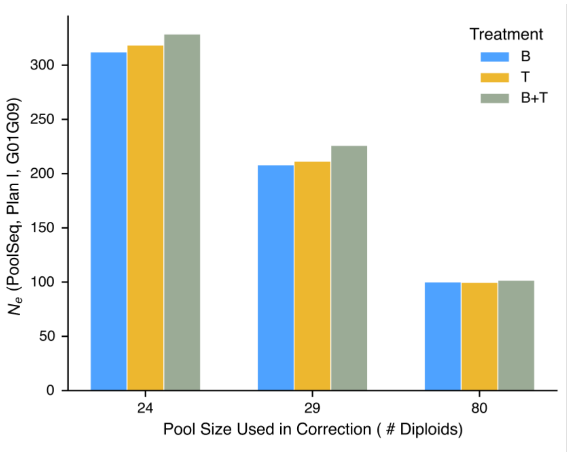

**Figure S16.** poolSeq $N_e$ (Plan I, G01-G09) at three pool-size corrections. With the true pool size included (80). With the founder-validated effective pool size (29), estimates rise to $N_e$ ~210-230, consistent with the F-regression. For downstream analyses we adopted $N_e$ ~250 as a working estimate for all three treatments, noting that the 95% block-bootstrap range spanned approximately 215-355 across treatments and methods. Treatment-specific signatures of a response to natural selection, if present, were better detected using the temporal covariance framework (G statistic and convergence correlations) applied in the following section.

Both the noise-corrected F-regression and the pool-size-corrected poolSeq estimator suggested an effective population size of $N_e$ ~200-350 per treatment. The two previously discordant estimates (F-regression $N_e$ =

165-420 vs poolSeq $N_e$ ~100) were reconciled by two corrections. First, subtracting the depth-dependent sampling noise from the F-regression, which removes a bias caused by non-constant sequencing depth across generations and second, using the empirically validated effective pool size ($N_{eff}$~29) in the poolSeq sampling correction. For downstream analyses we adopted $N_e$ ~250 as a working estimate for all three treatments, noting that the 95% block-bootstrap range spanned approximately 215-355 across treatments and methods.

As a third independent estimator, we computed the classical temporal method-of-moments drift estimator [30,33,34] following its recent application to temporal pool-sequenced data [35]. For each pair of samples j, k at site i within a single replicate, the standardized allele-frequency change is,

$$\hat{F}_{ijk} = \frac{(\hat{p}_{ij} - \hat{p}_{ik})^2}{\underline{p}_i(1 - \underline{p}_i)}, \qquad \underline{p}_i = (\hat{p}_{ij} + \hat{p}_{ik})/2,$$

and the sampling-corrected value is

$$\phi_{ijk} = \hat{F}_{ijk} - \frac{1}{2n_{ij}} - \frac{1}{2n_{ik}},$$

where $n_{ij}$ and $n_{ik}$ are the effective haploid sample sizes at site i at times j, k, computed from $N_{eff}$ = 29 and per-site read depth. The $N_e$ estimate is then,

$$\hat{N}_e = \frac{T}{2\underline{\phi}}$$

where $\underline{\phi}$ is the mean of $\phi_{ijk}$ across sites and T is the number of generations separating the two samples. Averaging $\phi$ across sites before inverting avoided inflation from division by near-zero values at individual sites. We aggregated across all pairs of consecutive available timepoints within each replicate, handling the G03-G05 gap in replicates 3 and 4 by setting T to the actual generation gap.

Applied to our data, the estimator yielded per-treatment harmonic-mean $\widehat{N}_e$ of 173 (**B**), 306 (**T**), and 183 (**B+T**). We report a 95% upper bounds as $\widehat{N}_e$ + 4.47SE across replicates, giving upper bounds of ~400-800 per treatment.

The convergence of all three estimators on $N_e$ below or near the census size suggests that the genome-wide drift is strong, wherein the genetic basis of host-plant tradeoffs is hypothesized to involve a small number of major-effect loci (e.g., those in or near chemosensory receptors, detoxification enzymes). If selection acted on a few loci, it may not have been able to overcome allele frequency variance generated by technical and drift noise at the whole genome scale of analysis. This left open the possibility that locus-specific signals of selection rather than overall variance magnitude was the primary signal of adaptation in this study.

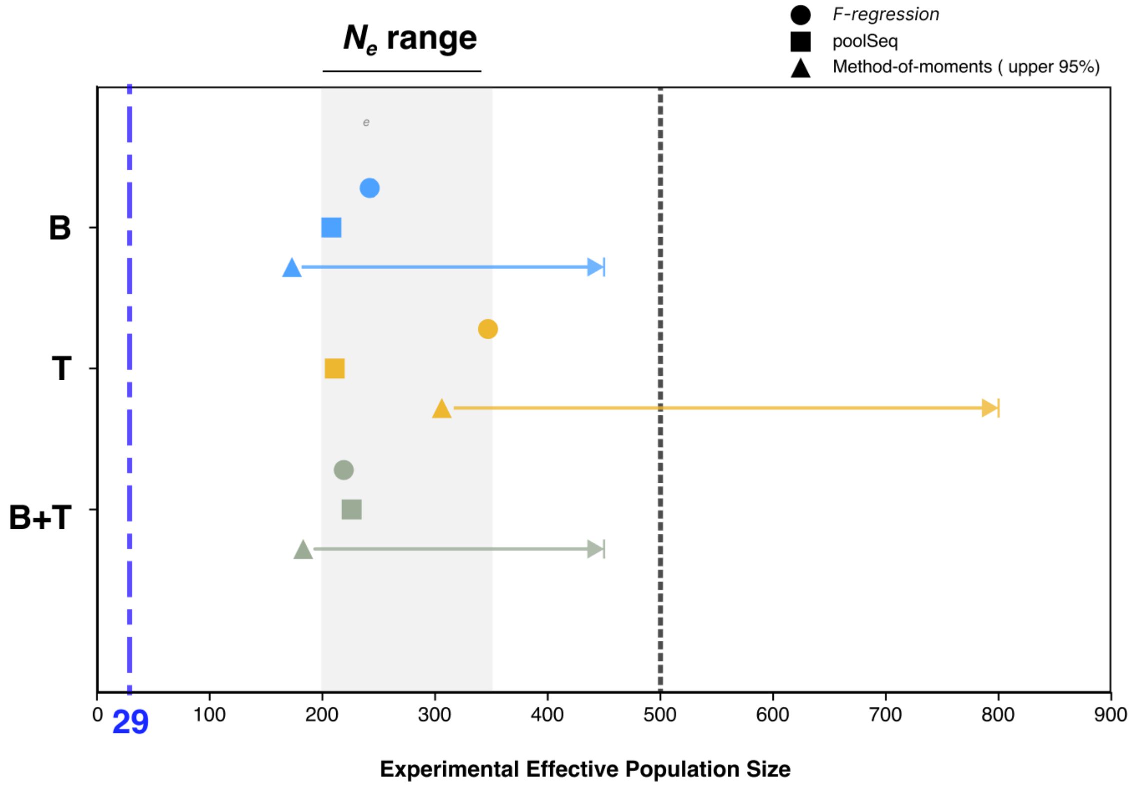

**Figure S17.** Summary of experimental effective population size estimates across three independent methods. Circles: noise-corrected between-replicate F-regression. Squares: within-replicate poolSeq windowed estimator (Plan I, $N_{eff}$ = 29). Triangles: Method-of-moments estimator with arrows extending to the 95% upper bound. Blue dashed line: effective pool size $N_{eff}$ = 29. Black dashed line: per-cage census size of reproductive populations used for infection of host plants (500 diploids). Gray band: Ne ~200-350 consensus range. All three estimators converged on $N_e$ below the per-cage census, supporting the a priori expectation that genome-wide allele-frequency variance was dominated by drift and technical noise when selection acted on a small number of loci.

### Genomic signatures of selection

#### Genome-wide contribution of selection to allele frequency change

Having established the noise floor and drift baseline in the preceding sections, we next addressed whether and to what extent linked selection is detectable above this baseline using the between-replicate covariance structure of the temporal covariance framework [36,37]. While the within-replicate temporal patterns (described above) characterized noise and drift, the between-replicate covariance captured whether independent replicate populations evolve in parallel, a signature of a parallel response to natural selection. This approach produced the statistic G, the fraction of total allele frequency change attributable to selection, and the convergence correlation, which measured the degree of parallel evolution across replicates.

We used the cvtkpy package (github.com/vsbuffalo/cvtk) to compute temporal variance-covariance matrices for each treatment (**B**, **T**, **B+T**). For each treatment, we included 4 replicates across 7 timepoints (generations 0, 1, 2, 6, 7, 8, 9; generations 3-5 were excluded because only 2 of 4 replicates were available). Each replicate used its own independent founder pool as the generation 0 measurement. We randomly assigned the tracked allele at each locus to avoid polarization bias. Sites were filtered to require depth $\geq$ 20 in all samples and minor allele frequency $\geq$ 0.05, yielding ~776,000 SNPs per treatment. Allele frequency changes were computed in 100 kb genomic windows with confidence intervals estimated using block bootstrapping to account for linkage disequilibrium. The standard bias correction assumes binomial sampling with $n = 2N_{pool}$ chromosomes, where $N_{pool}$ is the number of diploid individuals in the sequencing pool (80 in our experiment). However, unequal DNA contribution per individual in pooled sequencing creates overdispersion beyond the binomial expectation. We estimated the effective pool size ($N_{eff}$) using two independent approaches. First, the four founder pools (F1-F4) were used as independent draws from the same large colony (no drift between them). The between-founder variance, divided by the expected binomial variance, gives an overdispersion parameter $\psi = 2.78$, corresponding to $N_{eff} = 80/\psi$ or ~29 diploids. Next, we used an estimate inferred from the temporal covariance patterns for an independent measure of $N_{eff}$. The k=1 off-diagonal of the temporal covariance matrix (covariance between adjacent time intervals sharing a sampled timepoint) reflects sampling noise at the shared measurement. The observed k=1 covariance was 1.72x more negative than expected from binomial sampling with $N_{pool} = 80$ and de-duplicated sequencing depths, resulting in $N_{eff}$ of about ~24 diploids. These two estimates agree within 15% of each other ($N_{eff}$ is ~24-29). We present results here with both N=80 and $N_{eff}$=29 (founder-verified) corrections (Figs. S19, S22). The relationship between mean window depth and corrected covariance should be flat, at around zero.

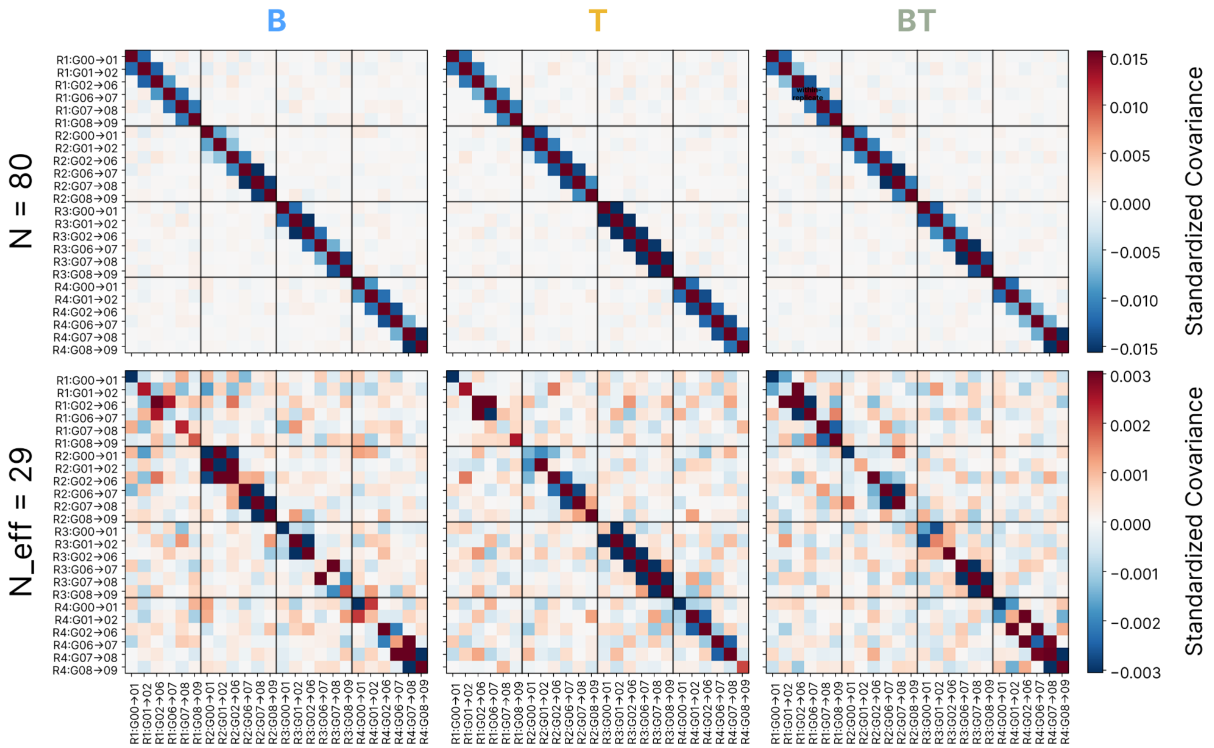

**Figure S19.** Temporal-replicate covariance matrix for each treatment (**B**, **T**, **B+T**), before (top row; pool size N = 80) and after (bottom row; effective pool size 29) the overdispersion correction. Each cell is the standardized covariance between Δp at two (replicate, interval) pairs. Under the uncorrected model, the k = ±1 off-diagonal (adjacent intervals within a replicate) showed a negative bias from pool-sampling noise at the shared timepoint. This was partially removed after correction with the founder-validated effective pool size. Between-replicate covariance blocks were near zero for all three treatments.

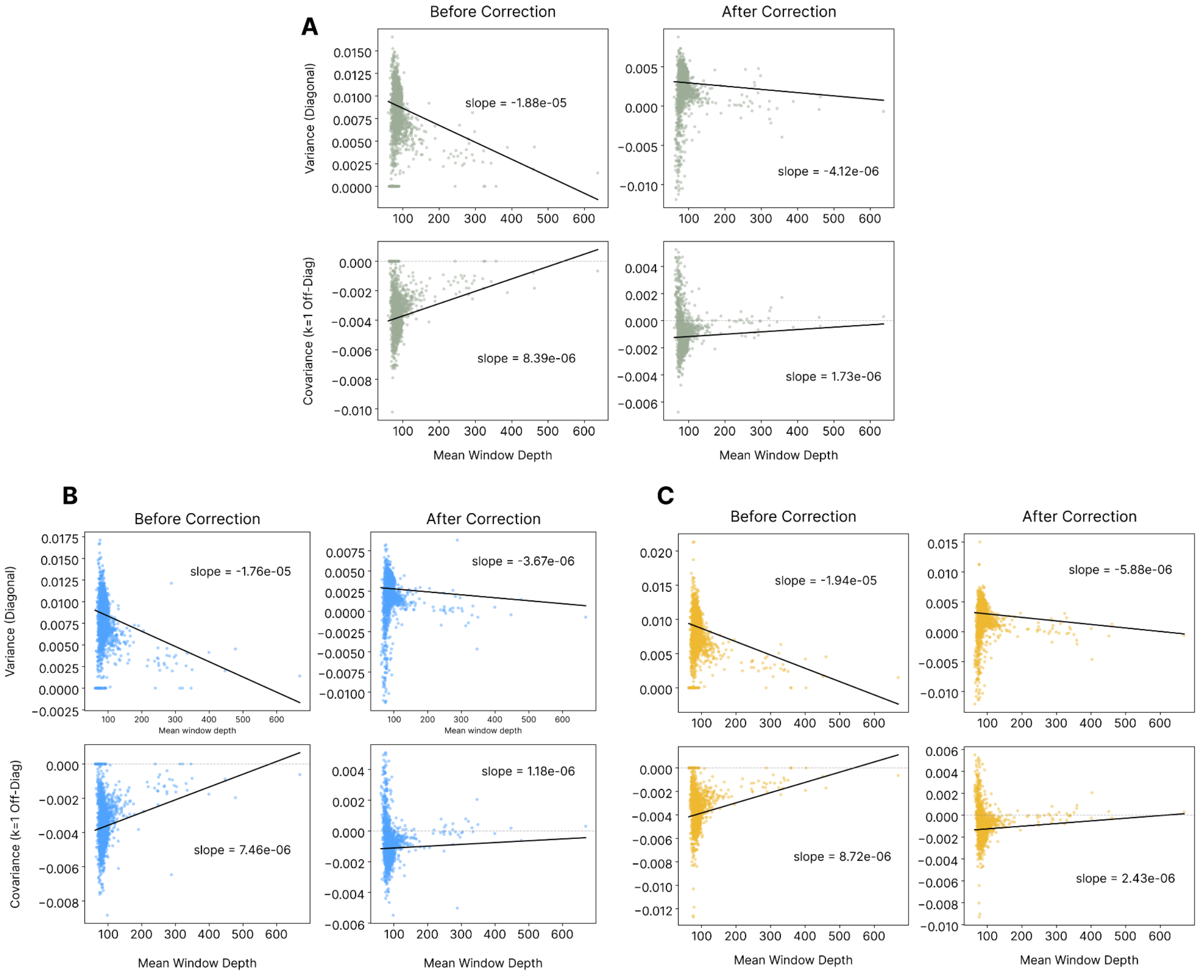

**Figure S20.** Bias correction diagnostic plots for the temporal covariance analysis. Each panel shows per-window (100 kb) variance (top row of each panel; the diagonal of the temporal covariance matrix) and adjacent-interval covariance (bottom row of each panel; the k=1 off-diagonal) plotted against mean sequencing depth, before (left) and after (right) the binomial sampling noise correction. (A) **B+T** treatment. (B) **B** treatment. (C) **T** treatment. Each point is one 100 kb genomic window; the black line is a regression of the metric on depth.

We next used the G statistic, including the k=±1 off-diagonal that requires the most sensitive bias correction. Because the k=1 correction is insufficient with N=80 and overcorrects with $N_{eff}$=29, we report G computed from only the k ≥ 2 off-diagonal elements, which are unaffected by the pool-size correction (they do not share a sampled timepoint). With the k≥2 G statistic (Fig. S21), only the **B+T** (spatially varying selection) treatment shows a positive trajectory across replicates (mean G = +0.063, one-sided p = 0.045). The **B** and **T** treatments show G indistinguishable from zero (**B**: -0.034, p = 0.76; **T**: +0.041, p = 0.25).

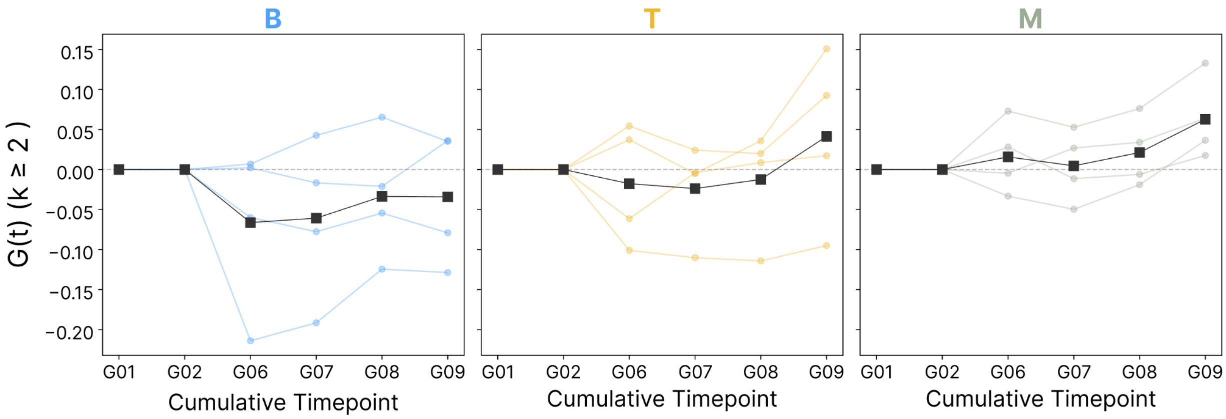

**Figure S21.** Cumulative G statistic (k≥2 off-diagonal only) per treatment, with individual replicate trajectories (colored lines) and replicate mean (black). G(t) is computed at each cumulative timepoint from generation 1 through 9. Only **B+T** showed a consistently positive trajectory across replicates.

We next computed the convergence correlation, which measures whether allele frequency changes in different replicates are correlated at the same sites [37]. Under the standard definition, the mean of the full convergence correlation matrix $C(t, s)$ across all $(t, s)$ time-interval pairs and all replicate pairs, the convergence correlation is sensitive to the pool-size correction. With $N = 80$, no treatment showed significant convergence. With the founder-validated correction ($N_{eff} = 29$) (Fig. S22), **B** and **B+T** values did not overlap zero (**B**: 0.022, 95% CI [0.014, 0.042]; **B+T** 0.012, CI [0.007, 0.034]), while **T** did not (0.009, CI [−0.002, 0.023]).

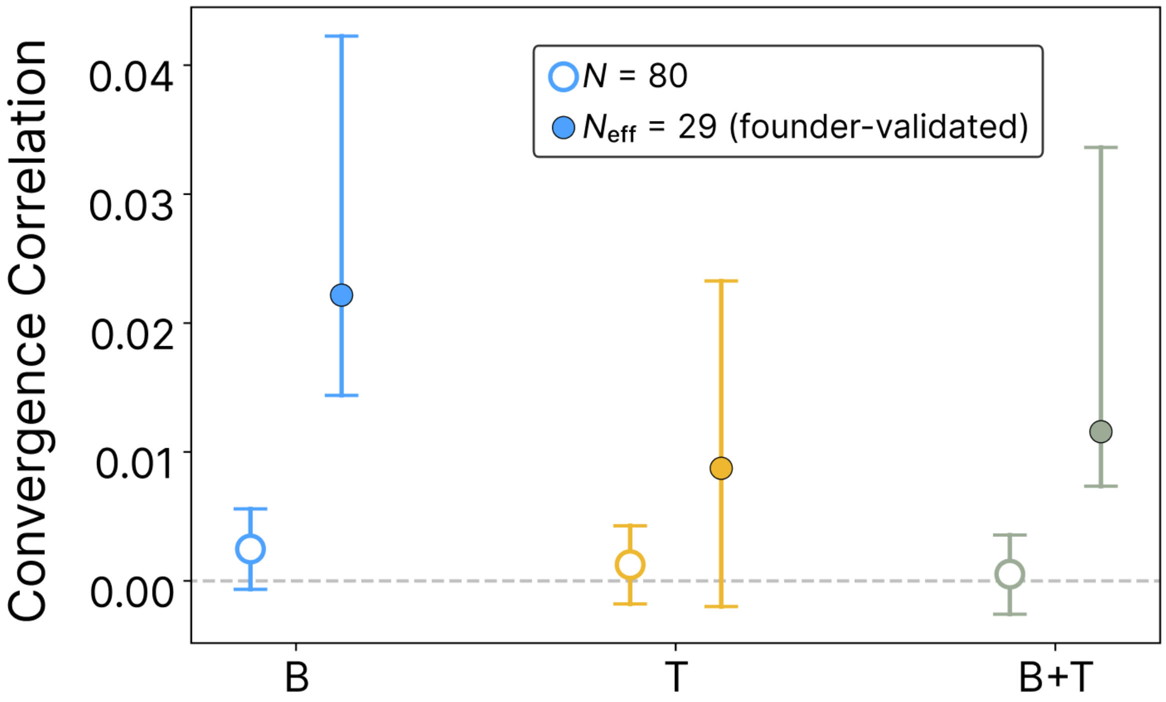

**Figure S22.** Convergence correlation estimated with pool size ($N_{pool}$=80) and founder-validated effective pool size ($N_{eff}$=29), with 95% block bootstrap confidence intervals. With $N_{eff}$ correction, **B** and **B+T** show confidence intervals that do not overlap zero.

The full-matrix convergence correlation is a conservative summary that averages over both within-interval (diagonal, $t = s$) and across-interval (off-diagonal, $t \neq s$) entries. Because the off-diagonal entries were dominated by noise in this dataset, the full-matrix mean diluted the within-interval parallelism signal. Decomposing the matrix corroborated the G statistic result above. The within-interval (diagonal) convergence correlation was strongest in **B+T** (0.053), moderate in **B** (0.035), and negative in **T** (-0.010). Therefore both the G statistic and the within-interval convergence correlation identified **B+T** as the treatment with the most consistent within-generation parallel allele frequency changes across replicates.

An alternative summary, the total-trajectory convergence correlation (computed from the cumulative allele frequency change $\Delta p_{G01 \rightarrow G09}$ rather than per-interval changes), gave higher values under a $N_{eff}$=29 correction: **B**=0.108, **T**=0.076, **B+T**=0.022. We found that using the total-trajectory approach there was less susceptibility to noise dilution arising from off-diagonal entries. These values suggested stronger detectable parallelism than the per-interval summary reported in Fig. S22.

| **Statistic** | **B** | **T** | **B+T** |
|---|---|---|---|
| Full-matrix cc (standard, reported above) | 0.022 | 0.009 | 0.012 |
| Within-interval cc (diagonal only) | 0.035 | -0.010 | 0.053 |
| **Total-trajectory cc ($\Delta$p G01→G09)** | **0.108** | **0.076** | **0.022** |

We assessed our power to detect convergence correlations by simulating 200,000 loci across 20 chromosomes with the same experimental design (4 replicates, 7 timepoints, $N_e$=250, de-duplicated depth=83) and a known convergence correlation, then ran the full cvtkpy analysis with block bootstrap confidence intervals. We varied the true convergence correlation (0-0.20), the number of replicates (4-20), the effective pool size (24-80), and the population effective size (100-500) to assess power. This analysis suggested that the minimum detectable convergence correlation is approximately 0.15-0.20, which is well above the 0.003-0.022 values estimated from our data. We found that under the variance structure present in our dataset, a total of 15 replicates would be needed to detect a 0.05 convergence correlation. The power analysis above uses the standard per-interval convergence correlation (full-matrix mean). If instead the total-trajectory convergence correlation is used (which gives values of 0.02-0.11 in our data), the estimated signal approaches the 0.15-0.20 detection threshold, suggesting that about 6-8 replicates may be needed to achieve sufficient power.

The primary limitation is the ratio of biological signal (drift variance, ~0.0005 per generation) to measurement noise (pool sampling variance, ~0.006 per timepoint). When each allele frequency is measured by pooled sequencing, we estimated that the measurement carries ~12x more noise than the single-generation drift signal it captures. When increasing the effective pool size from 24 to 80 (perfect binomial sampling), we found minimal improvements in power. When increasing the population effective size from 250 to 500, we found more substantial power improvement (Fig. S23).

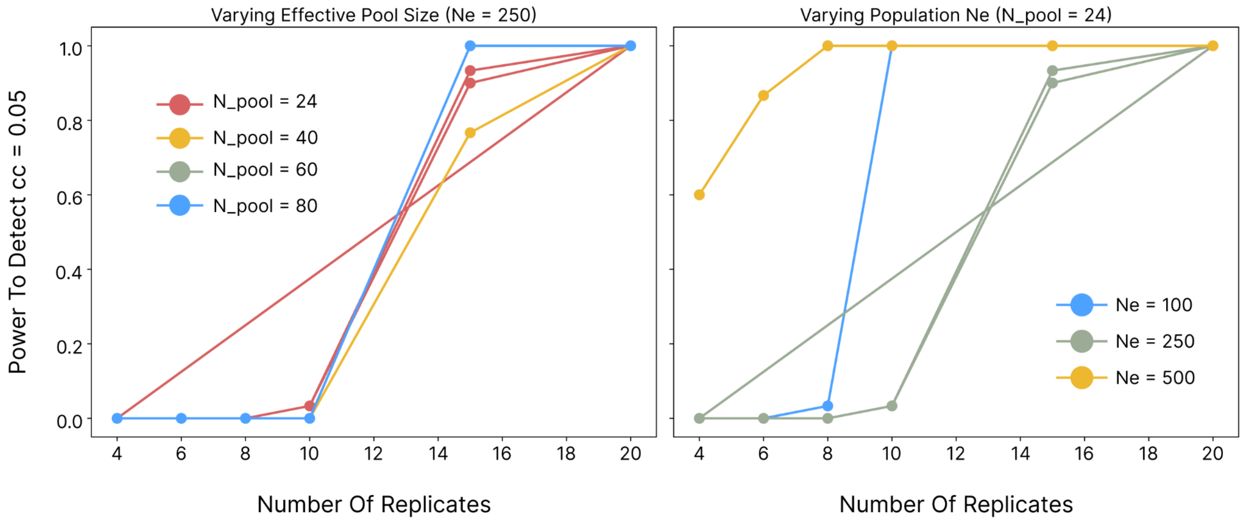

**Figure S23.** Power to detect a true convergence correlation of 0.05 as a function of the number of replicates, relative to our experimental parameters (4 replicates, $N_e$=250, $N_{pool}$ = 24). (Left) Different effective pool sizes ($N_e$ = 250 fixed). (Right) Different population effective sizes ($N_{pool}$=24 fixed).

Given the limited power of the temporal covariance framework, we more directly investigated the allele frequency changes between treatments for signatures of selection. To test for concordant vs. antagonistic selection between treatments, we computed the genome-wide and per-window (100 kb) correlation of mean allele frequency change (averaged across 4 replicates per treatment) between **B** and **T** from generation 1 to generation 9. We avoided the shared founder timepoint (generation 0) to prevent artifacts from shared sampling noise. Significance was assessed using z-transform per window, with FDR correction (Benjamini-

Hochberg). The mean allele frequency change from G01 to G09, averaged across 4 replicates, is positively correlated between **B** and **T** treatments at r = +0.021 (p = 0.0002), suggesting that alleles increasing on the **B** host also tended to increase on the **T** host. We conducted a windowed analysis (100 kb) and found that 75.7% of the genomic windows exhibited positive BvT correlation, with 45.8% significantly positive (p < 0.05) and only 6.5% significantly negative (Fig. S7). The dominant genome-wide selection signal was concordant across treatments, suggesting adaptation to shared laboratory conditions.

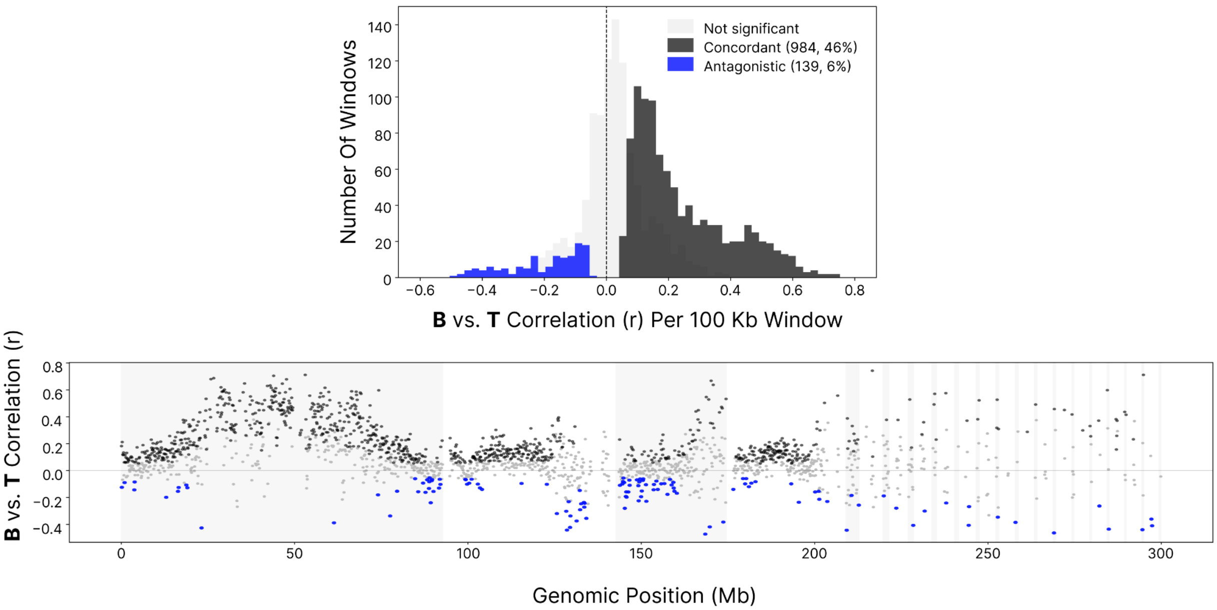

**Figure S7.** (Top) Distribution of windowed (100 kb) correlation of allele frequency change between **B** and **T**. The distribution is right-shifted (mean = 0.11, 76% positive), suggesting pervasive concordant selection for **B** and **T** (BvT) treatments. Significantly negative windows (blue, 6.5%) represent candidates for antagonistic host-specific selection for BvT. (Bottom) BvT allele frequency correlation along the three scaffolds with the highest concentration of antagonistic windows (chr_ScDA7r2_439, _597, _110).

The 139 windows (6.5%) with significantly negative **B** vs. **T** correlation represent candidate regions for host-specific antagonistic selection, or loci where alleles favored on one host are disfavored on the other. These antagonistic windows were concentrated on three scaffolds: chr_ScDA7r2_439 (39 windows), chr_ScDA7r2_597 (29 windows), and chr_ScDA7r2_110 (24 windows), which together account for 66% of the antagonistic signal (Fig. S7). After FDR correction, 33 windows remained significant at FDR < 0.10.

The **B+T** treatment imposed subdivided selection (250 flies on **B** + 250 on **T** each generation), so allele frequencies at strictly host-specific loci should have approached a balanced equilibrium and produced zero temporal autocovariance. In contrast, alleles beneficial in both environments (the convex region of Levins' fitness set) (see above) should have shown persistent directional change and positive temporal autocovariance. The finding that **B+T** was the only treatment with significantly positive $G(t)$ is consistent with persistent selection on generalist alleles that improved fitness across both host environments. This was further supported by the within-interval convergence correlation, which was high in **B+T**. The dominant concordant signal genome-wide (75% of windows showing positive BvT correlation) was consistent with selection on alleles whose effects were aligned across both host environments, likely a combination of shared laboratory adaptation and cross-host generalist alleles. A smaller set of windows (6.5%) were associated with antagonistic BvT selection and represent candidate loci where selection acted in opposite directions on the two hosts.

#### Haplotype structure and scale of linked selection in pool-sequenced time series

To characterize the scale of linked selection in our *S. flava* evolve-and-resequence experiment, we combined two complementary analyses. First, we detected haplotype blocks from the pool-seq time series using the haplovalidate program [38] and second, we measured linkage disequilibrium using 128 individually-sequenced founders and 64 individually-sequenced T2G07 (**T** flies, replicate population 2, generation 7) evolved flies using the ngsLD program [39].

We first identified candidate SNPs using a modified Cochran-Mantel-Haenszel (CMH) test implemented in the ACER R package [40], which corrected the variance of the standard CMH test statistic for both genetic drift and pool-seq sampling noise. We measured the extent of pool-seq overdispersion in the previous section ($N_{eff}$ = 29 diploids per pool). Because ACER requires two separate parameters that are frequently conflated (population effective size and pool sample size), we calibrated this analysis to the genome-wide patterns. We did this by scanning $N_e$ values from from 100 to 750 with pool size fixed at the founder-validated effective haploid count (58 = 29 diploid × 2). We found that $N_e$ = 250 produced a null p-value distribution closest to the theoretical expectation (median -log10(p) = 0.305 vs expected 0.301 under uniformity, 5.5% of sites at $p < 0.05$ vs expected 5.0%). We applied three parameter sets to test the sensitivity of this method. First, we used a pool size of 80, then an effective pool size, which propagated the previously measured overdispersion through ACER's pool-seq variance term and finally a BH q<0.05 candidate filter to identify the best blocks. Blocks were then clustered by haploReconstruct with haplovalidate's correlation-threshold selection, restricted to the four largest scaffolds (chr_ScDA7r2_597, _110, _439, _126, total 204 Mb). The minimum founder minor allele frequency was 0.05 and the minimum coverage was 20 in all 28 per-treatment samples. Candidates were capped at 15,000 per scaffold. We ran six treatment configurations: three single-treatment runs (**B**, **T**, **B+T** with 4 replicates each) and three pairwise runs (BxT, Bx[**B+T**], Tx[**B+T**] with 8 replicates combined). The pairwise runs have higher statistical power to detect shared selection signals but by construction detect only loci where replicates from both treatments respond in the same direction.

Haplovalidate detected 81 blocks in **B**, 78 in **T**, and 43 in **B+T** (Figure S10A). Single-treatment median block sizes differed among regimes: **B** and **T** were statistically indistinguishable (1.11 Mb and 0.78 Mb respectively; Wilcoxon p = 0.49), but **B+T** blocks were significantly shorter (0.43 Mb, **B** vs **B+T** p = 0.022, Figure S10A, bootstrap 95% confidence intervals shown as vertical bars). The smaller **B+T** blocks matched the biological prediction for the multiple-host regime where each leaf-mining *S. flava* larva experienced host-specific selection on only one of two hosts per generation, effectively reducing selective pressure at the population level and per locus by roughly half. However, we also found that the **B+T** block size distribution had a longer upper tail and the largest outlier block sizes among the three treatments.

The pairwise analysis runs (B×T, B×(B+T), T×(B+T), combining 8 replicates each) produced more but shorter blocks (medians 0.43-0.44 Mb) all identified as low-confidence-boundaries in the haplovalidate output. This was likely a power artifact of the 8-replicate design. With more replicates, within-scaffold correlations were always significantly above the cross-scaffold background, so the data-driven threshold could not be calibrated. Blocks from pairwise analyses should be interpreted as lower-resolution confirmations of parallel signals rather than as independent discoveries of new signals.

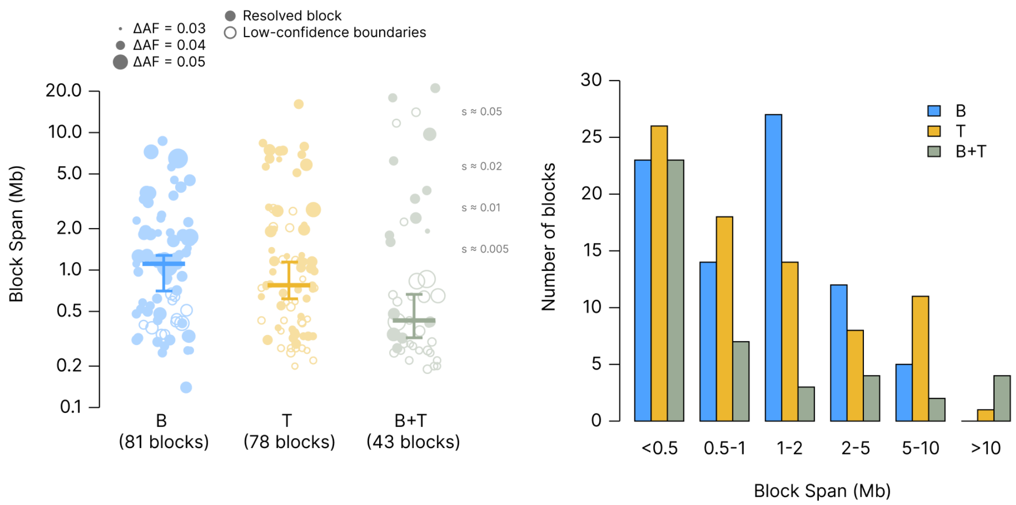

**Figure S10.** Haplotype-block sizes differed across treatments. (A) Block span (log scale) per treatment for all dominant blocks detected by haplovalidate under the calibrated parameter set ($N_e$ = 250, pool size = 58). Each point is one block; point size is proportional to the block's mean polarized ΔAF between G0 and G9. Filled circles show blocks with resolved correlation thresholds while open circles show blocks with low-confidence boundaries (where within-chromosome block correlations always exceeded the cross-chromosome background, so the algorithm could not calibrate a focal-vs-background threshold). Thick horizontal bars show the median block span per treatment; vertical bars span the 95% bootstrap confidence interval on the median (2000 resamples). Dotted reference lines show block-size predictions under classical sweep hitchhiking (block ~ $s \times \ln(N_e) / r$, with $N_e$ = 250 and r = 2 cM/Mb). (B) Block counts stratified by span class for each treatment. **B** and **T** have nearly all blocks in the 0.5-10 Mb range. **B+T** is shifted toward both very short (<0.5 Mb) and very long (>10 Mb) blocks, the latter reflecting regions where weaker per-locus power in the multi-host environment causes haplovalidate to merge what **B** and **T** resolve as multiple smaller blocks.

We considered block sizes in terms of simple hitchhiking theory where our median 1 Mb blocks correspond to selection coefficients of approximately s = ~0.01 under $s/r \times \log(N_e)$ with r = ~2 cM/Mb and $N_e$ = 250 (Figure S10A). Blocks of 2-5 Mb correspond to s ~0.02-0.05, consistent with moderate directional selection. **B+T**'s significantly smaller blocks (median 0.43 Mb, s ~ 0.005) are consistent with the diluted selection pressure of the Levene multi-host environment. However, **B+T** also possessed four blocks spanning more than 10 Mb (up to 21 Mb on scaffold 597), which neither **B** nor **T** produced (Figure S10A-B). These regions contain 10-12 **B**-specific and 3-10 **T**-specific blocks within their boundaries (i.e., they are regions where **B** and **T** both detected dense block-level signal, but **B+T**'s weaker per-locus power causes haplovalidate to cluster them into single megaregions). The larger haplo-blocks therefore may reflect scale-of-resolution differences across treatments with varying power.

Each block's polarized mean allele frequency (SNPs oriented to the rising allele, averaged across replicates and SNPs) changed by a median ΔAF ~ 0.04 from G0 to G9 (Figure S24A), consistent with s ~0.015-0.02 under a simple linear-change approximation. To characterize trajectory shape without imposing an arbitrary early/late split, we computed the area under each normalized trajectory curve (AUC; 0.5 = linear, > 0.5 = saturating, < 0.5 = accelerating; illustrated in Figure S24). All three treatments' median AUC fell significantly below 0.5 (**B** = 0.397, **T** = 0.401, **B+T** = 0.375; Wilcoxon $p < 10^{-11}$ in each case), indicating that by G9 selected haplotypes were still rising and had not reached equilibrium. **B+T** accelerated later than **B** or **T** (**B+T** vs **B** Wilcoxon $p < 10^{-4}$), again consistent with diluted Levene-like selection. These patterns suggest that ΔAF and block sizes we report are lower bounds on equilibrium response, which we likely did not reach by the end of the experiment given the patterns observed in this analysis.

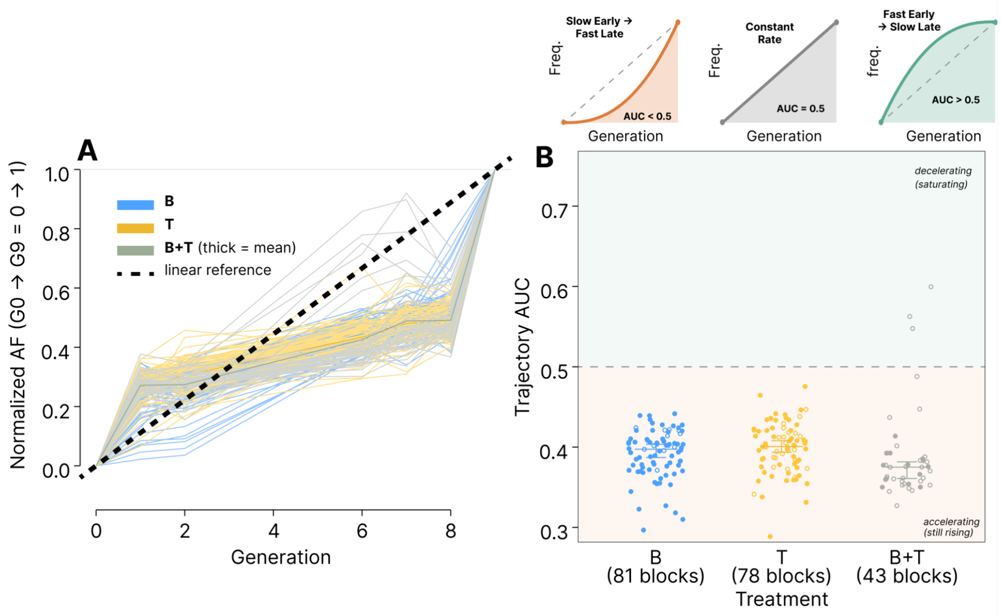

**Figure S24.** Block trajectories were still accelerating at generation 9. Cartoon schematics illustrate trajectory shape summarized by AUC, the area under each normalized trajectory curve: a linear rise from G0 (0) to G9 (1) gives AUC = 0.5; slow-then-fast (accelerating) curves fall below 0.5; fast-then-slow (saturating) curves exceed 0.5. (A) Every haplovalidate block's mean polarized allele-frequency trajectory, rescaled so G0 = 0 and G9 = 1. The dashed black diagonal is the linear reference. All three treatments' per-block trajectories sit below the linear reference on average, indicating slow early and fast late change. (B) Per-block trajectory AUC values. All three treatments' median AUC is significantly below 0.5 (Wilcoxon vs 0.5, $p < 10^{-11}$ each; median **B** = 0.40, **T** = 0.40, **B+T** = 0.38), indicating blocks had not reached equilibrium frequencies by generation 9. **B+T** trajectories were the most accelerated (**B+T** vs **B**: Wilcoxon $p < 10^{-4}$), consistent with a diluted per-generation selection effect under the multiple host environment.

#### Linkage disequilibrium (LD) using individually-sequenced experimental flies

The analyses discussed thus far focus on LD structure inferred from temporally replicated pooled sequencing and identified scaffold 439 as hotspot for parallel host-selection signals. We addressed this directly by measuring LD in 128 founder individuals and 64 T2G07 (**T** of replicate 2 at generation 7) evolved individuals. Each individual was sequenced to ~0.5× coverage and the resulting sequencing was used to compute genotype likelihoods for LD estimation (ANGSD + ngsLD). To reduce potential confounding effects of SNP ascertainment, we restricted all comparisons to the intersection of SNP positions passing ANGSD's SNP filter in both cohorts. This yielded 102,211 / 109,340 / 126,130 / 166,870 shared SNPs across scaffolds 110, 126, 439, and 597 respectively, and produced pair-count-matched comparisons between founder and T2G07 $r^2$ in every distance by region comparison.

Within scaffold 439, the hotspot (2-12 Mb) had lower $r^2$ than the flanking region in both cohorts and at every distance bin (Figure S8A; hotspot/flanking ratios in founders: 0.894 at <1 kb, 0.934 at 1-10 kb, 0.934 at 10-100 kb; T2G07: 0.910, 0.990, 0.999). A segregating inversion spanning the hotspot would produce the opposite pattern (i.e., uniform elevated $r^2$ across the arrangement interval, particularly at intermediate-to-long distances where inversion-suppressed recombination dominates over intra-arrangement decay).

We then compared cohorts on the matched SNP to show that the hotspot's LD deficit relative to flanking closes in T2G07 (Figure S8B). At 1-10 kb the ratio shifted from 0.934 (founders) to 0.990 (T2G07). At 10-100 kb, it shifts from 0.934 to 0.999. Absolute $\Delta r^2$ at the hotspot was small ($|\Delta| \leq 0.0015$ at intermediate and long distances and a ~-5.7% change at <1 kb and because the starting LD in the outbred founders is already near the background rate at all but the shortest distances). The ngsLD analysis therefore validates both the no inversion interpretation and the pool-seq-inferred scale of linked selection.

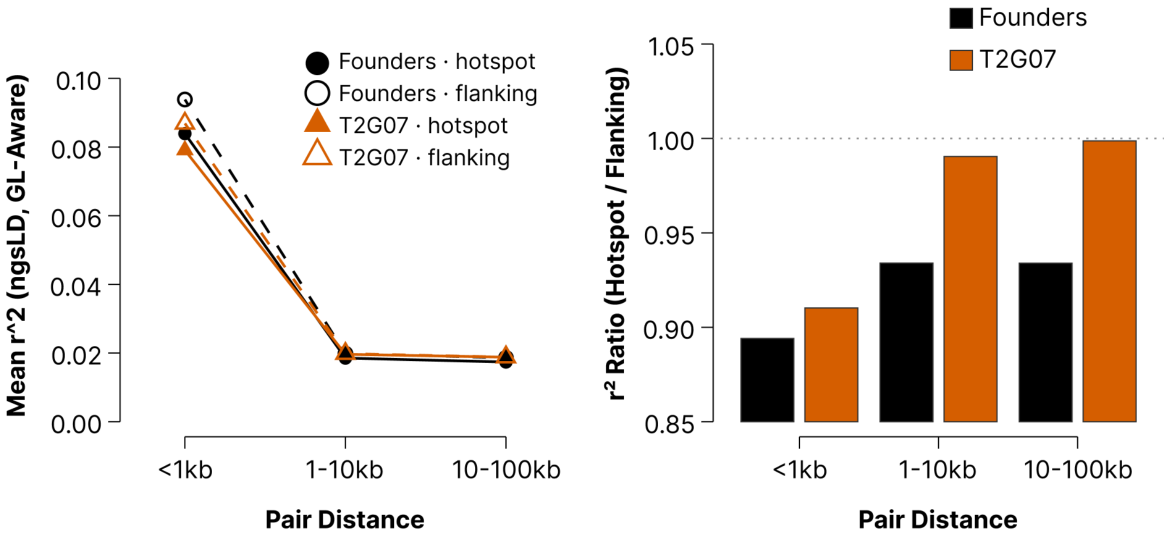

**Figure S8**. LD comparisons at matched SNPs on scaffold 439 using genotype likelihoods and ngsLD on 128 individually-sequenced founders and 64 T2G07 individuals (~0.5× coverage each). Restricted to 126,130 SNPs called in both cohorts. (A) Mean $r^2$_ExpG (ngsLD's genotype-likelihood-aware estimator) versus inter-SNP distance bin for four comparisons: founders x hotspot (2-12 Mb), founders x flanking, T2G07 x hotspot, T2G07 x flanking. Founders in black, T2G07 in orange. Hotspot in solid lines/filled symbols, flanking in dashed lines/open symbols. $r^2$ in the hotspot is lower than in the flanking region in both cohorts at every distance. (B) Hotspot/flanking $r^2$ ratio per cohort and distance bin. Dotted reference line at 1.0 marks no hotspot deficit. The ratio rises from ~0.93 in founders to ~0.99 in T2G07 at 1-10 kb and 10-100 kb.

The hotspot-level result above aggregated ~126,000 SNPs and millions of LD pairs across 10 Mb, making small changes in $r^2$ detectable. At the scale of individual haplovalidate blocks (0.5–5 Mb, 20–500 SNPs each), the signal falls below the noise of single-timepoint $r^2$ estimates. We confirmed this by demonstrating that per-block within-block $\Delta r^2$ (T2G07 - founders) showed a positive trend relative to matched flanking windows, but no block-level comparison reached statistical significance. Per-block PCA on individual genotype likelihoods (PCAngsd on 192 individuals) also found no haplotype cluster structure (PC1 explained only 4-5% of variance per block, where founders and T2G07 were indistinguishable), as expected given the short number of generations included in the experimental time series. The founders were derived from a large-$N_e$ wild population where background LD decayed rapidly. After only 7–9 generations of cage evolution with $N_e$ ~250, selection-driven $r^2$ at distances that match the block lengths we detected (0.1–10 Mb) is $10^{-4}$ per SNP pair, which is below the sampling noise of 64-128 individuals at 0.5× coverage. In contrast to the single-generation and single-population measures of LD, the haplovalidate method detected blocks using coordinated allele-frequency change across 10 time points and 4 replicate populations, giving a ~40-fold increase in effective observations compared to a single $r^2$ snapshot. Genome-wide windowed $r^2$ from individual sequencing was flat at ~0.017 across all four scaffolds in both founders and T2G07, with no elevation at haplovalidate block positions and no detectable difference between cohorts (Figure S9). The elevated $r^2$ windows that did appear corresponded to low-SNP-density regions (< 50,000 pairs per 500 kb window) where repeat content inflated $r^2$ estimates. Block-level validation of haplovalidate output will require either deeper individual sequencing (≥ 5× per individual) or long-read haplotype phasing.

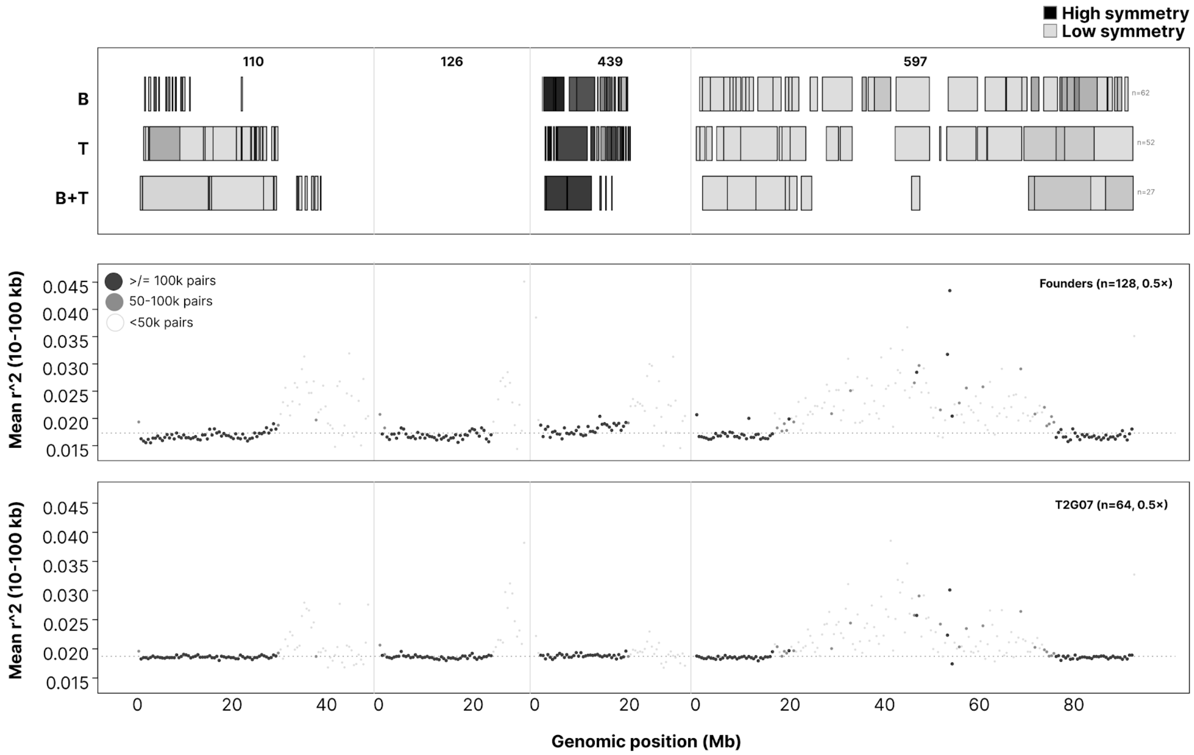

**Figure S9.** Temporal replication reveals haplotype structure not detectable with single population LD mapping. (Top) Haplovalidate blocks detected from pool-seq allele-frequency trajectories across 10 generations and 4 replicates per treatment, shown as three horizontal tracks (**B** / **T** / **B+T**). Each block is a horizontal bar at its genomic position. Bar shade represents the PBS symmetry score on a grayscale gradient. Block counts per treatment are **B** = 62, **T** = 52, **B+T** = 27. These blocks come from a different haplovalidate run than the calibrated set in Fig. S10, which detected 81, 78 and 43 blocks. Scaffold 439 between ~2 and 12 Mb is dominated by high-symmetry blocks in all three treatments. (Middle) Genome-wide windowed $r^2$ (500 kb non-overlapping windows, 10–100 kb inter-SNP pairs) from 128 individually-sequenced founder flies (~0.5× coverage each), computed via ngsLD on the shared-SNP set. Point shade indicates estimation confidence based on pair count. Dashed line marks the genome-wide mean of well-estimated windows ($r^2$ ~0.017). No elevation was detected at haplovalidate block positions, including the scaffold 439 hotspot. (Bottom) The same as the middle panel for 64 T2G07 evolved individuals (~0.5× each). The $r^2$ landscape was indistinguishable from founders locally but is elevated genome-wide, suggesting the build-up of LD over the experimental time series. No block-level or hotspot-level signal emerged from single-timepoint LD at this coverage. The contrast between panels demonstrates the ~40-fold information advantage of temporal–replicate pool-seq (top) over single-timepoint individual sequencing (middle, bottom) for detecting selection-driven haplotype movement in populations founded from a high-Ne source with fast ancestral LD decay.

#### Generalised linear mixed model and parametric-bootstrap likelihood-ratio test

To test at each individual SNP whether the trajectory of allele-frequency change across the ten generations of selection differed between the two single-host treatments, we fit a per-SNP binomial generalised linear mixed model (GLMM) and assigned a p-value to the generation x treatment interaction by parametric bootstrap from the fitted null. Allele counts were computed as $success_{i,j} = AF_{i,j} \times N_{eff,i,j}$ and $total_{i,j} = (AF_{i,j} \times N_{eff,i,j}) + ((1 - AF_{i,j}) \times N_{eff,i,j})$ and applies three pre-filters per site (every retained sample has total ≥ 15, ≥ 6 samples have success > 0, and ≥ 3 distinct success/total ratios are observed across samples.

For each SNP, we performed two binomial GLMMs, the null and full model. Both used the binomial family with weights equal to total read depth, a logit link, and a random intercept by replicate cage (four **B** cages and four **T** cages). Generation was treated as a continuous covariate ($t_1$-$t_{10}$) and treatment was treated as a two-level factor (**B** vs **T**) with **B** as the reference. The fit used the Nelder–Mead optimiser (fast=true). We assigned a p-value to each LRT statistic by parametric bootstrap from the fitted null. For each of n_Sims = 200 simulations (LRT_NSIMS = 200), we simulated a per-observation success vector under the null model fit, refit both the null and full models to the simulated counts retaining the original design matrix and weights and then recorded the simulated LRT. The p-value is therefore the number of simulations whose simulated LRT met or exceeded the observed LRT and n_valid is the number of simulations whose null and full-model refits both converged. We reported the p-value only when n_valid > 0.5 x n_Sims (SNPs failing this convergence threshold were removed). The final GLMM table contained 813,431 SNPs (slightly more than the 812,670-SNP canonical set because of processing differences) with 812,590 (99.9%) returning OK, 841 returned ObservedFitFailed, and 22.5% of OK SNPs carrying the singular flag (random-intercept variance estimated at zero, suggesting small between-cage variance relative to the time series).

#### Concordance vote among methods for identifying signatures of antagonistic selection

We localized antagonistically-selected loci genome-wide by building a per-window vote-based scan that integrated five independent statistical tests in 200-kb non-overlapping windows across the four chromosome-scale scaffolds (chr_ScDA7r2_110, _126, _439, and _597). The five tests, in the order shown in Figure 3A, are: (1) is the most-negative tail of the per-window allele-frequency-change covariance cov($\Delta\bar{p}_B$, $\Delta\bar{p}_T$), where $\Delta\bar{p}_X$ is the per-SNP mean of AF($t_{10}$) - AF($t_1$) across the four replicates of treatment X covariance is computed across SNPs within each non-overlapping 200 kb window via numpy.cov (unbiased, ddof=1; minimum 50 SNPs per window); (2) the z-standardized slope of the grenedalf temporal $F_{ST}$ (**B**, **T**) statistic across $t_1$-$t_{10}$, fitted per window by a linear mixed-effects model; (3) the empirical p-value from a permutation test (all 70 four-versus-four relabellings of the eight pools) on $t_{10}$ **B** vs. **T** $F_{ST}$; (4) the per-window mean of per-SNP generation x treatment LRT_chisq from the GLMM in the previous subsection; and (5) the per-window maximum of the per-SNP BayPass C2 contrast statistic, taken across all three treatment contrasts (**B** versus **T**, **B** versus **B+T**, and **T** versus **B+T**), among the twelve t10 pools, conditional on the among-pool covariance matrix Ω. All five tests are summaries of the same laboratory time series under different model assumptions. They are therefore not statistically independent of one another, which is why we require agreement among several of them rather than treating any single test as decisive. Each test was thresholded at the genome-wide top-5% percentile (or $p \leq 0.05$ for the permutation $F_{ST}$), calibrated by $10^4$ density-stratified permutations conditional on per-window n_snps to control for the confounding between window-level test scores and SNP density. Three haplovalidate-block tracks (per-treatment **B**, **T**, **B+T**) are included in Figure 3A as orthogonal evidence of linked selection but are not included in the vote count.

The $t_{10}$ **B** vs. **T** permutation $F_{ST}$ test (test 3 above) included eight $t_{10}$ pools labelled B1–B4 and T1–T4 (there are 70 combinations to partition them into two groups of four and the true **B** vs **T** labelling is one of those 70). For each 4/4 split we computed the mean between-group Hudson $F_{ST}$ per 200-kb window using pool-seq corrected effective chromosome counts ($N_{eff}$ = 58 = 29 diploids x 2, matching the founder-overdispersion estimate from the variance-decomposition section above). The per-pair Hudson $F_{ST}$ numerator and denominator follow $num_{\{i,j\}} = (p_i - p_j)^2 - p_i(1 - p_i)/(N_{eff} - 1) - p_j(1 - p_j)/(N_{eff} - 1)$ and $den_{\{i,j\}} = p_i(1 - p_j) + p_j(1 - p_i)$. Per-window aggregation is a sum-of-numerator divided by the sum-of-denominator (ratio-of-averages form). For each of the 70 4/4 splits of the cross-group pair set, we produced a per-window and per-split $F_{ST}$. The empirical p value per window is the rank of the observed (true **B**/**T**) $F_{ST}$ among the 70 splits. The smallest achievable p is 1/70 ~0.014. The null preserved all drift, shared-laboratory-adaptation, and within-pool sampling noise at $t_{10}$ and only relabeled which four pools were considered as **B** and which were considered as **T**.

The temporal $F_{ST}$ slope test (test 2 above) was computed upstream by a pool-seq $F_{ST}$ pipeline using grenedalf $F_{ST}$ to compute Hudson's pairwise $F_{ST}$ between **B** and **T** pools at each generation, followed by a linear mixed-effects model fitting $F_{ST}$ on generation per window with treatment-pair as a per-(**B** replicate x **T** replicate) random-intercept term. The slope coefficient and its z-statistic (slope / $SE_{slope}$) were stored per window. The permutation $F_{ST}$ and slope tests were non-redundant. The slope test integrated information across all ten generations and favors windows where between-treatment divergence accumulates monotonically with time, while the permutation $F_{ST}$ used only the $t_{10}$ endpoint and favored cumulative divergence regardless of trajectory shape. A window with steady early divergence that plateaus before $t_{10}$ will pass the slope test but may not pass the permutation $F_{ST}$, and vice versa.

The per-window vote count was votes = $\Sigma$ $IND_t$ summed over the five binary indicators, with maximum possible vote = 5. A single 200-kb window (chr_ScDA7r2_439_HRSCAF_779:2,800,000–3,000,000) reached the unique genome-wide maximum of 4/5 votes. The four hits at this window are the antagonistic cov($\Delta p_B$, $\Delta p_T$) tail, the $t_{10}$ permutation $F_{ST}$, the per-SNP generation x treatment LRT and the BayPass $C_2$ treatment contrast among the generation-10 pools. The fifth test ( z-scaled temporal $F_{ST}$ slope) placed the same window at the 84.5th genome-wide percentile (slope_div_z = 12.05 vs top-5% threshold 13.69). Independent recovery of the same window across four of the five summary statistics was parsimoniously explained by a single locus under host-antagonistic selection at this region.

#### Signal block, founder LD block, and gene-track annotation

Two nested intervals on chr_ScDA7r2_439_HRSCAF_779 are referenced throughout the rest of this section. The signal block is the 200-kb window scaffold 439:2,800,000–3,000,000 that reaches the 4/5 vote in the multiple-test scan above. The founder LD block is the surrounding ~1-Mb haplotype block scaffold 439:2,640,000–3,610,000 defined by ngsLD on the 128 individually-sequenced founder flies (see Haplotype structure and scale of linked selection above). The signal block contains the 200-kb LRT peak. The LD block extends both flanks of the peak to capture the haplotype background that hitchhiking is expected to span. All polarised AF-trajectory plotting (Figure 3B) was restricted to the signal block, and the 339-SNP top-LRT subset of the q* iteration (Figure 3C) was restricted to the LD block. The per-replicate diversity test used the LD block as the focal interval. We extracted protein sequences for every annotated CDS in scaffold 439:2.50–3.75 Mb (the LD block plus ~140-kb flanks on each side) by parsing the v2 GFF for CDS records on chr_ScDA7r2_439, sorting and concatenating the genomic CDS sequences, reverse-complementing minus-strand genes, and translating in the standard reading frame. Proteins were compared against the *D. melanogaster* UniProt reference proteome (UP000000803, 13,826 sequences) using diamond blastp v2.1.10 in --more-sensitive mode, retaining the top-1 hit per query at e-value ≤ $10^{-5}$.

### Wild-lab parallels

#### Wild host-associated differentiation in *Scaptomyza flava*

The evolution experiment was initiated from a founding colony of n = 130 wild-emerged *S. flava* adults collected at a single New Hampshire locality (58 emerged from leaves of *B. vulgaris*, 72 from *T. glabra*). Specifically, we collected third-instar *S. flava* larvae that had formed mines in the leaves of naturally established *B. vulgaris* (**B** flies) and *T. glabra* (**T** flies) host plants at the three sites (AV, PS, RM) within the same general geographic area, bagged them on the parental leaf, and then reared them to adult emergence. Between 22 and 60 emerged adults per host and site were pooled and sequenced with paired-end short reads to produce six wild pool-seq libraries (n = 22–60) that we used as a natural genomic context for interpreting the laboratory selection experiment.

| **Site** | **B wild** | **T wild** |
|---|---|---|
| AV | AVB | AVT |
| PS | PSB | PST |
| RM | RMB | RMT |

All six pools were aligned to the *S. flava* reference genome assembly (sfla_v2.fa) with BWA-MEM and deduplicated with Picard as in the experimental evolution samples. All wild analyses use the experimental variant set of 812,670 biallelic SNPs called jointly from the lab **B**/**T**/**B+T** pools. We restricted the wild pools to this set for two reasons. First, every site was biallelic and polarized to the same set of reference and alternative allele pairs, allowing read-count to be comparable across wild, founder, and G10 experimental pools. Second, this was the same site set used by every other analysis in this study (lab BayPass, PBS, grenedalf diversity, haplovalidate).

For each wild pool we generated an mpileup at those 812,670 positions and counted reference and alternative reads, masking indel and read-start regions before counting alleles. We excluded sites with fewer than five reads in any wild pool. Because we tested only sites that were already segregating in the lab founders, the wild analyses are blind to variants private to the wild. This biases toward the relevant set for the wild to lab comparisons. We asked to what extent wild *S. flava* carried allele-frequency differences between *B. vulgaris*- and *T. glabra*-associated cohorts after controlling for population structure among the three sampling sites. Given that pairwise $F_{ST}$ (Figure S11) showed that population structure was at least as large as within-site host differentiation, we used BayPass [41] to estimate a genome-wide population-covariance matrix Ω and to test for per-SNP host-associated shifts conditional on Ω (Figure S25).

Between-site $F_{ST}$ (median 0.004–0.005) was at least as large as within-site **B**-vs-**T** $F_{ST}$ (median 0.003). AV had the strongest within-site **B** vs **T** differentiation ($F_{ST} = 0.010$). PS was near zero ($F_{ST} = -0.002$, i.e. the **B** and **T** pools at PS were more similar than random within-site pairs). RM was intermediate ($F_{ST} = 0.003$). There was no uniform **B** > **T** or **T** > **B** pattern across sites. Because between-site $F_{ST}$ was as large as within-site $F_{ST}$, any outlier scan that did not correct for population structure was dominated by site-level noise.

Table S6. Wild-pool pairwise $F_{ST}$ stratified by pair category. Within-site **B** vs **T** (n = 3 pairs) median $F_{ST}$ was comparable to between-site same-host (n = 6 pairs) median $F_{ST}$, indicating that geographic structure across the three New Hampshire localities (AV, PS, RM) was at least as large as within-site host-associated differentiation.

| **Category** | **Pairs** | **Median $F_{ST}$** | **Range** |
|---|---|---|---|
| Within-site **B** vs **T** | AVB vs. AVT, PSB vs. PST, RMB vs. RMT | 0.003 | [-0.002, 0.010] |
| Between-site, same host | AVB vs. PSB, AVB vs. RMB, PSB vs. RMB, AVT vs. PST, AVT vs. RMT, PST vs. RMT | 0.005 | [0.000, 0.010] |

| Category | Pairs | Median $F_{ST}$ | Range |
|---|---|---|---|
| Between-site, different host | AVB vs. PST, AVT vs. RMB, etc. (9 pairs) | 0.004 | [-0.000, 0.008] |

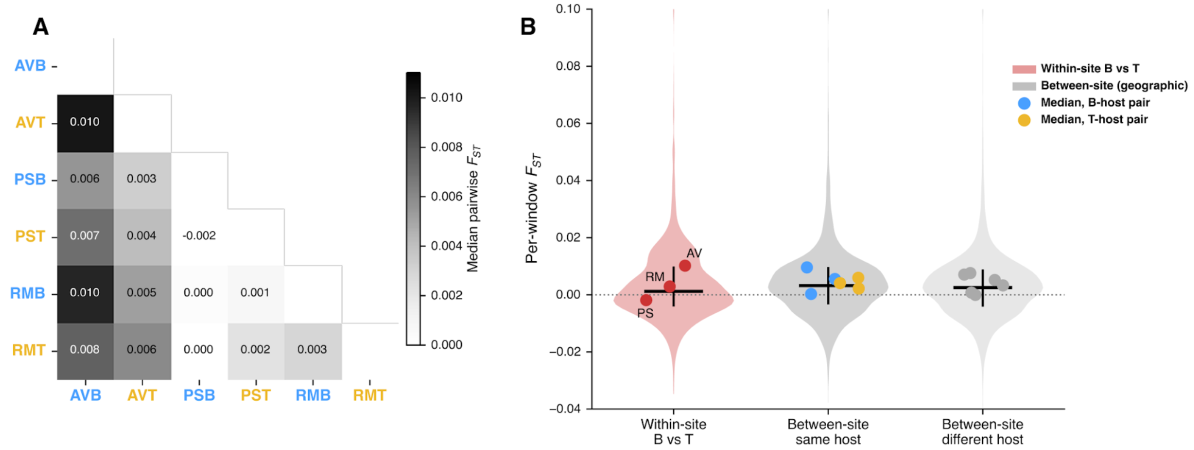

**Figure S11:** (A) Heatmap of median pairwise $F_{ST}$ among the six wild pools on 390-kb non-overlapping windows. AVB was the most divergent pool genome-wide (top row 0.006–0.010 to all others). PSB vs PST had a slightly negative median $F_{ST}$ (-0.002), suggesting no host-associated differentiation at PS at this genomic scale. (B) Per-window $F_{ST}$ distributions pooled by pair category (within-site **B** vs **T**, between-site same-host, between-site different-host), with per-pair medians overlaid.

We therefore conditioned on an estimate Ω of the genome-wide among-pool covariance in order to isolate SNPs whose host-associated differentiation exceeded the drift and sampling noise background (the component of the **B**-vs-**T** comparison that was shared across all three sites). Our BayPass run on the wild pools produced three outputs from a single Ω. We considered the Ω core model (pool-drift model fit on a thinned SNP subset), $\beta_i$covariate model (per-SNP regression of scaled allele frequency on a host covariate, Ω fixed), $C_2$contrast model (per-SNP Bayesian contrast between the three **B**-host pools and the three **T**-host pools, Ω fixed). We thinned the 812,670 SNPs to every 16th site (--thin-step 16) and used BayPass default MCMC settings (20 pilot runs of 500 iterations, a 5,000-iteration burn-in, and 1,000 post-burn-in samples thinned every 20 iterations). The result was the 6x6 scaled population-covariance matrix used as the null for the two SNP-level tests below (Figure S25A). Before testing per-SNP host-associated changes conditional on Ω, we checked that Ω had correctly absorbed the drift covariance. Under the Ω-only null, the per-SNP differentiation statistic $X^tX_{st}$ was expected to follow a $\chi^2$ distribution (6 d.f. = 6 pools). We ran the core model on the thinned SNP set and compared the observed distribution to the theoretical (Figure S25B). The observed quantiles were close to theoretical in the bulk of the distribution (observed median 4.83 vs. $\chi^2$ median 5.35), suggesting that Ω captured the genome-wide drift covariance. At the upper quantiles the observed values shifted above theoretical (8.8% of SNPs above the $\chi^2$ 5% tail, 3.3% above the 1% tail), suggesting an excess of host-plant driven population genomic differentiation above the drift null expectation.

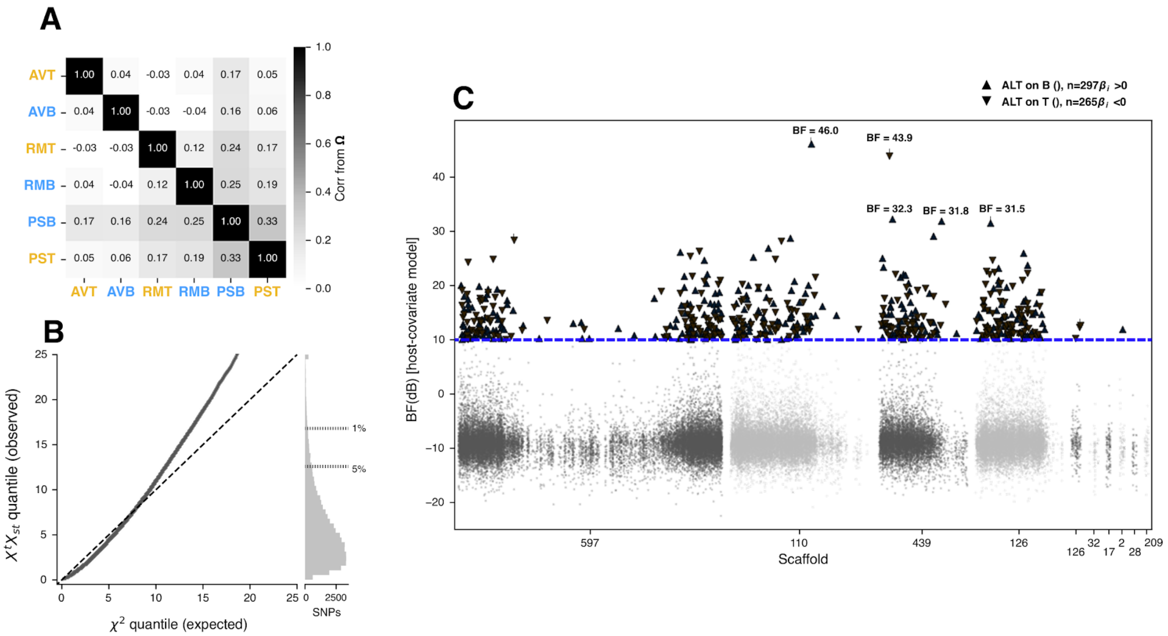

**Figure S25.** Wild BayPass scan for host-associated differentiation conditional on Ω. (A) Correlation matrix derived from Ω, reordered by hierarchical clustering with **B**-host pool labels in blue and **T**-host labels in yellow. AVB and AVT were the most divergent pools. (B) Q-Q plot of observed $X^tX_{st}$ quantiles from the Ω model (thinned SNP set, n = 50,792) against $\chi^2$ expected quantiles with null expectation as dashed line. The bulk of all SNPs sit on the line (observed median 4.83 ~ $\chi^2$ median 5.35) which confirmed that Ω absorbed drift covariance. The upper-right tail rises above the line (8.8% of SNPs above the $\chi^2$ 5% tail, 3.3% above the 1% tail), suggesting a small excess of host-plant specific differentiation. The histogram on the right shows the raw distribution of observed $X^tX_{st}$ values at the same y-coordinates. (C) BF(dB) per SNP from the $\beta_i$ host-covariate model (conditional on Ω) across the top-10 scaffolds of the *S. flava* assembly. Dashed line at BF = 10 marks the threshold used in BayPass to identify significant regions (decisive SNPs). Decisive SNPs that are triangle-up = $\beta_i > 0$ are those in which the alternative allele was enriched on **B** (n = 297) and those that are triangle-down = $\beta_i < 0$ were those in which the alternative allele was enriched on **T** (n = 265). The top-5 SNPs by BF were annotated with their BF values.

We next coded a binary host covariate c with AVB = PSB = RMB = +1 and AVT = PST = RMT = -1 and ran the function g_baypass against the full 812,670 SNPs. The per-SNP columns that were produced were: $\beta_i$ = posterior mean of the standardized regression coefficient of allele frequency on the host covariate, BF(dB) = Bayes factor for the host-covariate model versus the Ω-only null. Under our coding (**B** = +1, **T** = -1), a positive $\beta_i$ was the alternative allele enriched on **B** hosts. A BF(dB) > 10 cutoff corresponded to decisive SNPs. We used the $C_2$ contrast fit to the same data under the same Ω but re-analyzed with host as a discrete group assignment (AVB / PSB / RMB vs AVT / PST / RMT), producing per-SNP contrast statistics. The $\beta_i$ covariate and the $C_2$ contrast tested independent hypotheses. The covariate model treated host as a continuous predictor and penalized cross-site variance in $\beta_i$, whereas the contrast model treated the two hosts as fixed groups. We reported both below and used BF(dB) for genome-scan ranking.

| **Statistic** | **Value** |
|---|---|
| SNPs tested | 812,670 |
| BF(dB) range | [-23.0, +46.0] |
| BF(dB) median | -9.26 |
| SNPs with BF(dB) > 10 (decisive) | 574 (0.07%) |
| SNPs with BF(dB) > 15 | 174 |
| SNPs with BF(dB) > 20 | 49 |
| Mean $\lvert\beta_i\rvert$ among BF(dB) > 20 | 0.0057 |
| $C_2$ contrast $-\log_{10}(p) \geq 3$ | 1,316 |
| $C_2$ contrast $-\log_{10}(p) \geq 5$ | 14 |
| Spearman(BF, $-\log_{10}p$) | 0.24 |

| **Statistic** | **Value** |
| --- | --- |
| Top-5% SNP-set overlap (BF ∩ $C_2$) | 21.6% |

Of 812,670 tested SNPs, 574 had BF(dB) > 10, 174 had BF > 15, and 49 had BF > 20 (about 0.07% of the genome at the decisive threshold). About 99% of SNPs had BF < 0 (more likely under the Ω-only null than under the host-covariate model), suggesting that any signal of host-driven differentiation after one generation of selection in the wild was sparse. The $C_2$ contrast returned 1,316 SNPs at $-\log_{10}(p) > 3$ (about 0.16%) and 14 at $-\log_{10}(p) > 5$. Spearman rank correlation between BF(dB) and $C_2$ $-\log_{10}(p)$ was $\rho = 0.24$ (Pearson $r = 0.29$). The two top-5% sets overlapped at 21.6% (above the null 5%). At the extreme BF tail (BF > 20, n = 49) the mean absolute standardized effect size was $|\beta_i| = 0.006$, suggesting that the top-BF SNPs were often outlier-like (high posterior confidence, near-zero effect), whereas $C_2$ re-weighted by group-wise mean difference and so preferentially highlighted SNPs with visibly shifted group means. We used BF(dB) for genome-scan ranking but note that the joint tail of $\beta_i$ and $C_2$ provides additional evidence of host-based differentiation.

In summary, we found that population structure among the three New England sites was at least as large as within-site differentiation between flies reared on different host plants (median between-site $F_{ST}$ ~0.005 vs within-site $F_{ST}$ ~0.003). In addition, the direction of the associations were not consistent across wild sampling sites ($F_{ST}$ = 0.010 at AV, ~0 at PS, intermediate at RM). Therefore, any test of host-driven differentiation, especially after the 1 generation of selection in the wild captured here, would need to condition on a model of among-pool covariance to avoid being dominated by population structure. BayPass conditioning on a 6×6 genome-wide covariance matrix Ω addressed this, with the the bulk of the $X^tX_{st}$ quantiles tracking the $\chi^2$ null but with an upper tail rising above expectation (8.8% above the 5% tail, 3.3% above the 1% tail), which we interpret to be a small but systematic excess of host-driven differentiation. Third, after conditioning, the per-SNP signal is sparse with only 574 SNPs (0.07%) reaching BF(dB) > 10. The standardized effect sizes were small (mean effect = 0.006 at BF > 20). These patterns suggest that wild

*S. flava* on *B. vulgaris* and *T. glabra* are weakly differentiated at a small set of SNPs once population structure is removed.

#### Sign concordance of lab-to-wild allele-frequency correlations

We compared signatures of laboratory antagonism between **B** and **T** flies to wild allele frequency divergence between flies reared from the same host plants in the wild. For each SNP we computed lab effect direction (sign of **B** vs **T** allele-frequency change at $t_{10}$) and wild effect direction (sign of **B** vs **T** $\Delta p$ averaged across the six wild pools). We then asked whether sign agreement was enriched on the X-chromosome ortholog (Muller A) relative to the four autosomes. Density-matched bootstrap null distributions were drawn per Muller element (n = 1,000 size-matched draws) to test for enrichment. We separately fit a per-SNP linear regression of wild $\Delta p$ on lab $\Delta p$, restricted to the scaffold 439 LD-block and to a genome-wide background, to test whether the focal hotspot drove the lab-to-wild correspondence.

We started from the lab GLMM SNP table (per-SNP gen×trt interaction LRT and bootstrapped p-values) and applied three additional filters. First, we retained SNPs in the top 50% of the genome-wide generation x treatment interaction LRT distribution. This removed the bottom half of the LRT distribution where lab effect-size signs were dominated by per-replicate slope sampling noise rather than by reproducible between-treatment differences. Second, we required $|z| \geq 2$ on both $s_B$ and $s_T$, where z = estimate / SE and the SEs are the per-replicate scipy.stats.linregress standard errors of the logit slopes for each replicate. This removed SNPs with sign change between treatments that were poorly resolved by the lab time series. At the SNP level, with only 4 **B** and 4 **T** cages and 10 generations, $|z| < 2$ on either coefficient was considered a “no information” site. Third, we required founder pool minor-allele frequency in [0.05, 0.95]. Outside of this interval, the pool-seq detection floor (binomial sampling at per-pool $N_{eff} \sim 29$) dominates the per-SNP $\Delta p$ estimate and the resulting wild-AF and lab-$\Delta p$ signs were unreliable. After applying these three filters, the background SNP set was n = 4,050 SNPs, with 709 (Muller A / X), 695 (B / 2L), 786 (C / 2R), 790 (D / 3L), and 999 (E / 3R).

For each background SNP, lab Δp is the unweighted mean over the four lab replicates where we subtracted the alternate-allele frequency at $t_{10}$ in **T** treatment from the alternate-allele frequency at $t_{10}$ in **B** treatment: $lab_{\Delta p} = (1/4)\ \Sigma_{\{r=1..4\}}[AF(B_{r,t10}) - AF(T_{r,t10})]$. Wild Δp is the unweighted mean over the three wild collection-site pairs (AVB/AVT, PSB/PST, RMB/RMT) of $p_B$-$p_T$. We calculated $wild_{\Delta p} = (1/3)\ \Sigma_{\{site \in \{AV,PS,RM\}\}}[AF(site_B) - AF(site_T)]$. Both Δp values are unpolarized (alternate-allele coding as called in the joint VCF). The unweighted mean is appropriate because the three sites have comparable per-pool sequencing depth.

For each SNP we computed the indicator $I_{concord} = 1[sign(lab_{\Delta p}) = sign(wild_{\Delta p})] \in \{0, 1\}$, treating SNPs with lab Δp = 0 or wild Δp = 0 as ties. Under independent unsigned lab and wild Δp the chance expectation is 0.50. We reported binomial standard errors $\sqrt{(p(1-p)/n)}$ on observed fractions throughout. Each background SNP was assigned to a *D. melanogaster* Muller element via synteny. For Muller A, we overlaid the concordance fractions at the two scaffold 439 candidate regions. This included the signal block (scaffold 439:2,800,000–3,000,000, n = 28 background SNPs, observed concordance 0.71 ± 0.09) and the surrounding ngsLD-defined LD block (scaffold 439:2,640,000–3,610,000, n = 67 background SNPs, observed concordance 0.58 ± 0.06). To assess whether these values were unusual given size and founder-MAF distribution, we constructed an AF-matched bootstrap null per candidate region and per Muller element. For each of 1,000 bootstrap iterations and each candidate SNP, we sampled with replacement one control SNP from the Muller's background pool whose founder MAF was within ±0.05 of the candidate's. We then computed the iteration concordance fraction across the n_target sampled controls. The bootstrap median and 2.5/97.5 percentile interval were reported as the per-Muller control background reference in Figure 3D. Both the signal block (0.71) and LD block (0.58) showed concordance fractions above the 95% upper bound of every Muller's matched control bootstrap, and within Muller A alone the bootstrap medians (0.43 signal-block-sized, 0.43 LD-block-sized) were well differentiated from the observed values, confirming that the elevated scaffold 439 concordance is not a Muller A artifact.
